# Environmental DNA analysis to search for an endangered desert fish

**DOI:** 10.64898/2026.05.20.726406

**Authors:** Alanna Fulkerson, Alexsandre Gutiérrez-Barragán, Alejandro Varela-Romero, Douglas K. Duncan, Mariana Mateos

**Affiliations:** Department of Ecology and Conservation Biology, Texas A&M University, College Station, Texas, USA; Laboratorio de Ecología Molecular, Departamento de Investigaciones Científicas y Tecnológicas (DICTUS), Universidad de Sonora (UNISON), Hermosillo, Sonora, México

**Keywords:** Cienega, Sonoran Desert, Madrean Archipelago, Poeciliidae, endangered, environmental DNA, Mifish, metabarcoding, conservation, Leuscicidae

## Abstract

We conducted field surveys over two consecutive years (2022 and 2023) of freshwater habitats in the Rio De La Concepcion (several tributaries and sites) and one site in the Rio Santa Cruz (tributary of the Gila River) in the state of Sonora, Mexico. Our primary goal was to detect the presence of the Rio Concepcion Topminnow (*Poeciliopsis jackschultzi*), an endangered microendemic livebearing desert fish whose external morphology is indistinguishable from several sympatric congeners. Using an environmental DNA (eDNA) metabarcoding approach (“MiFish” locus), we failed to detect evidence of *P. jackschultzi*, implying that it is extinct or present at abundances below our detection ability. Applying a collective evidence approach including visual/eDNA surveys and past records, we discuss the strengths and limitations of the eDNA metabarcoding approach. In the Rio De La Concepcion, we confirm the presence of the other known native and previously reported introduced teleosts, and reveal more recent introductions. In the Rio Santa Cruz site, we detect three putative non-natives (Asexual Hybrid Topminnow, Yaqui Sucker, and Mexican Roundtail Chub). We recommend further monitoring of habitats and fish taxa, and implementation of practices that improve groundwater recharge and the quality and quantity of treated wastewater in the study area.

## 2. Introduction

We are currently confronting what is regarded as the sixth global mass extinction because the current rate of biodiversity decline far exceeds pre-anthropogenic background estimates [1, 2]. This biodiversity crisis is most severe in freshwater ecosystems [3]. Even though they cover less than 1% of Earth’s surface, freshwater habitats contain about one third of all vertebrate species and 40% of all fish species [4–8]. Due to their natural water scarcity, freshwater ecosystems in arid lands are particularly vulnerable to anthropogenic activity, including climate change [9, 10]. Fishes and other aquatic taxa native to arid environments have been greatly impacted, and many native species in desert ecosystems have declined or have been extirpated from parts of their native ranges due to these and other anthropogenic stressors, such as establishment of non-native species [11–15].

As documented by Miller [16], Southwestern North America, housing the Great Basin, Sonoran, and Chihuahuan Deserts, typifies the arid region experiencing the longest and most drastic anthropogenic disturbances affecting freshwater ecosystems. This region is characterized by low species richness but high endemism of fishes [14]. Between the end of the nineteenth century and 1961, Miller [16] reports that at least six fish species in this region had become extinct, and at least 13 “additional forms” were at “levels from which they might not be able to recover”. By 1989, at least 20 fish species in this region had become extinct [17], and ∼120 of the remaining 200 taxa were considered of special concern [18]. In the same region, approximately three decades later, 32 fishes and 23 aquatic invertebrates are considered extinct, and five fishes extinct in the wild [14]. As expected, species that occupy more restricted ranges (a.k.a. narrow endemics or microendemics) are more susceptible to anthropogenic stressors such as water scarcity and introductions of nonnative species.

Efforts to prevent extinction in the wild rely on monitoring efforts to document the status and trends of target wild populations. Up until recently, capture-based surveys (using gear such as seines, electrofishing, minnow traps, and dip nets) were the only practical approach available for monitoring desert fish populations. Furthermore, in systems where external morphology is insufficient for distinguishing sympatric species, lethal (whole specimen sampling) or non-lethal (e.g. fin tissue sampling from live specimens for genetic analysis) was required, which could further jeopardize persistence of endangered populations by removal or injury of individuals, respectively. Environmental DNA (eDNA) has emerged as an alternative or complementary non-invasive approach for monitoring populations of many taxa in many types of environments [19]. Environmental DNA refers to the pool of DNA isolated from environmental samples (e.g. soil, water, air, sediment, biofilm), which includes both organismal and extra-organismal, including extracellular, DNA [20]. The collected material is typically passed through filters, meshes or nets, aimed at the retention of organisms, organismal debris (e.g. cells and free DNA) or particles of a desired size [20], from which DNA is subsequently isolated.

Most eDNA studies employ one of two Polymerase Chain Reaction (PCR)-based approaches. One approach uses quantitative PCR (qPCR) and can detect one or a few target species using taxon-specific primers (and probes). The other approach, known as metabarcoding, uses conventional PCR with “universal” primers containing unique DNA barcodes for each sample, followed by high throughput sequencing. Consequently, metabarcoding can identify many more taxa within an ecosystem than qPCR [21]. Metabarcoding is thus useful when the appropriate reference sequence libraries are available and the targeted genetic marker contains adequate taxon-diagnostic variation [22, 23]. However, metabarcoding is reported to exhibit amplification bias towards some non-target species [23, 24], and appears to have lower ability to detect low concentration targets compared to qPCR methods [24]. One major drawback of eDNA studies, along with any PCR-based study that aims to detect target DNA that is in low amounts and/or highly degraded (e.g. ancient DNA), is that it is particularly susceptible to contamination from DNA not present in the collected sample. Thus, rigorous sterile procedures and controls are required to prevent or account for amplification of non-target DNA [21, 25, 26]. In addition, eDNA metabarcoding approaches are reported to yield sequences that are not present in the sample through method-induced artifacts [27], which should be considered during data interpretation. The present study uses eDNA metabarcoding to survey teleost taxa in filtered water samples collected within the known range of a microendemic, endangered, and morphologically cryptic desert fish (*Poeciliopsis jackschultzi*, Rio Concepcion Topminnow), and nearby habitats (Fig. 1).

**Figure 1.**
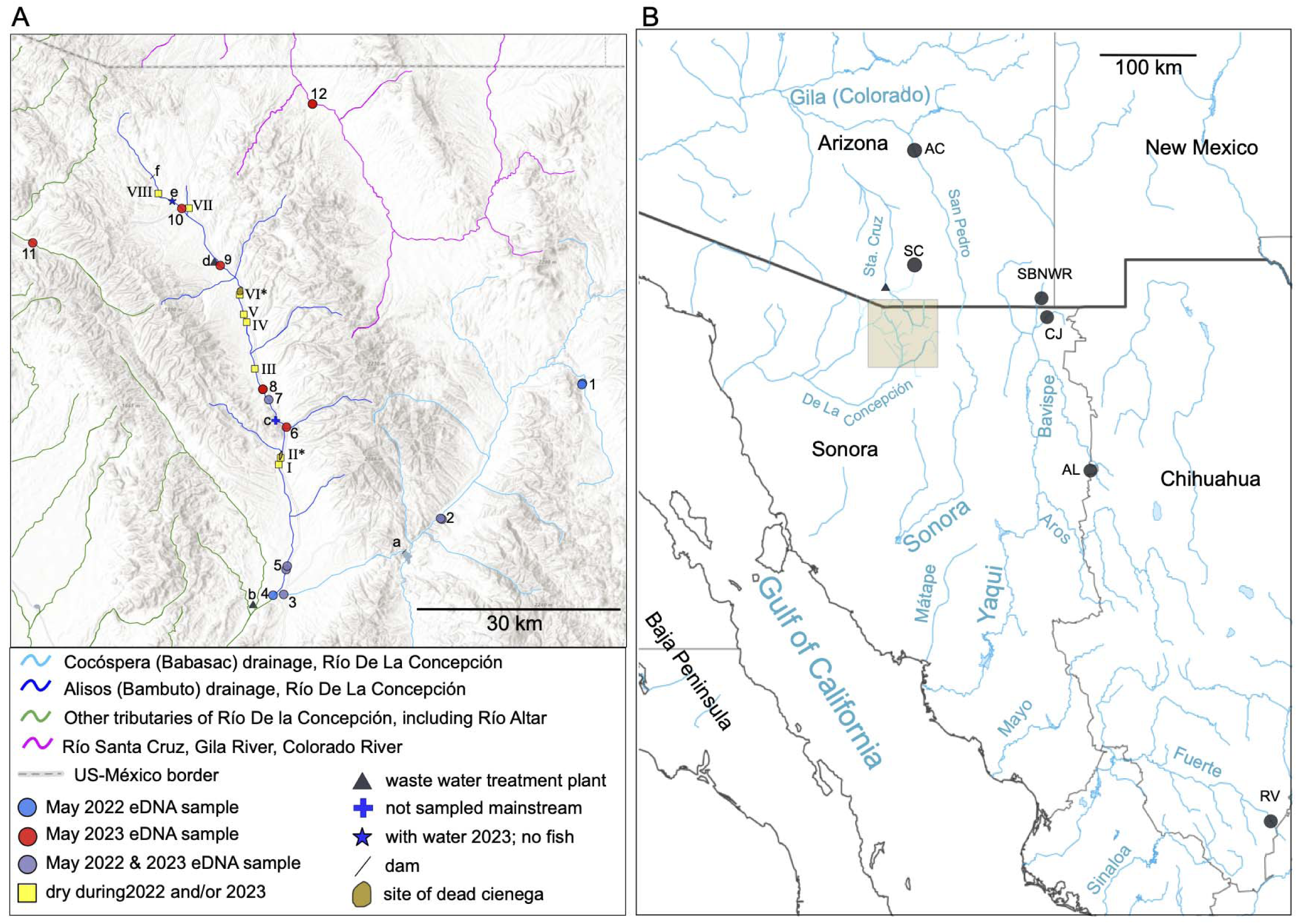
Maps of the study area. **A.** Some tributaries of the Rio Concepción (turquoise, blue and green) and the Rio Santa Cruz (magenta) are depicted, most of which are intermittent. Environmental DNA samples were collected at the following sites (indicated by filled circles). 1 (2 sites): Cuitaca Upstream and Cuitaca Downstream. 2: Aribabi (2 sites): Ciénega and Mainstream. 3: Ímuris-Cocóspera. 4: Bambuto-Cocóspera (i.e. confluence of the Cocóspera and Alisos branches). 5: Ímuris-Bambuto (2 sites): Mainstream and Sidestream. 6: Agua Caliente. 7: La Atascosa. 8: La Cieneguita. 9: Cíbuta (∼1 km downstream of Los Alisos Wastewater Treatment Plant = d). 10: La Mesa. 11: La Arizona: 12. Mascareñas (Río Santa Cruz). The following localities of interest are indicated. a = Presa (dam) Comaquito. b = Presa (dam) El Muro. c = site, not sampled, along the mainstream that likely maintains permanent flow. e = short stretch with water but no fish May 2023. Yellow squares with Roman numerals indicate sites that were dry during our 2022 or 2023 visit (Dataset S11). Dry sites II* and VI* are within or adjacent to what former cienega habitats [28, 45]. Photos/videos of the habitat during our field expedition are provided in Dataset S14. **B.** Beige shaded rectangle indicates the area depicted in panel A. Turquoise font indicates the Gulf of California and relevant rivers/tributaries. SC = approximate location of Sonoita Creek and its headwater Monkey Spring. AC = Aravaipa Creek. The following three localities of the Bavispe basin of the Rio Yaqui are indicated: SBNWR = San Bernardino National Wildlife Refuge, San Bernardino Creek; CB =Arroyo Cajón Bonito; and AL = Arroyo El Largo (Rio Yaqui site for *Gila minacae* reference sequence). Triangle shows Nogales International Wastewater Treatment Plant, which treats some of the wastewater from Nogales, Sonora. Maps were generated with the ArcGis Online application [Sources: Esri, TomTom, Garmin, FAO, NOAA, USGS, (c) OpenStreetMap contributors, and the GIS User Community], including the North American Environmental Atlas - Lakes and Rivers layer [46]; and edited with Inkscape v.1.4.2 [47].

*P. jackschultzi* was described in 2019 based on only a handful of specimens [28], even though it was discovered by R.C. Vrijenhoek in the 1980’s. *P. jackschultzi* has a highly restricted and discontinuous distribution along a ∼30 km stretch of the Río De La Concepción (hereafter Rio Concepcion), state of Sonora, Mexico [28] (Fig. 1). *P. jackschultzi* prefers spring-fed cienega habitats [29], and the cienega site where it exhibited the highest abundance (in the vicinity of site 7; Fig. 1) was obliterated as result of the federal highway lane expansion sometime between 1986 and 1994. Before the present study, the last field monitoring efforts reporting the presence of *P. jackschultzi* were in 1999-2001, during which its relative abundance in remaining habitats was very low [28]. *P. jackschultzi* is currently listed as endangered in the IUCN Red List [30], and in Mexico’s proposed updated list of threatened and endangered species (Table S1). Like most narrow endemic desert fishes, the major threat to *P. jackschultzi* appears to be the reduction of adequate habitat caused directly by human infrastructure (e.g. highway construction) and activities (e.g. trampling/pollution caused by cattle ranching), or indirectly through decreased quantity/quality of ground and surface water, likely exacerbated by climate change [28]. Non-native fishes, which have been documented in the region [e.g. Tilapia, the Western Mosquitofish Gambusia affinis, and others; 31, 32, 33], and are considered threats to other similar native species (e.g. *Gambusia* for *Poeciliopsis occidentalis*, [34]), may also contribute to the demise of *P. jackschultzi*.

A challenge to monitoring efforts is that *P. jackschultzi* is practically impossible to distinguish based on external morphology from its sympatric congeners, including its much more broadly distributed close relative *Poeciliopsis occidentalis sensu lato* (hereafter Gila Topminnow), which is listed as threatened in Mexico and endangered in the US (Table S1). The historical range of *Poeciliopsis occidentalis s.l.* spanned the Gila River Basin (part of the Colorado River) and its tributary the Santa Cruz River (US and Mexico), and the following Mexican rivers (Ríos): Concepcion (including the entire known range of *P. jackschultzi*); Sonora; Yaqui (spans US and Mexico); Matape; and Mayo (Fig. 1B). *Poeciliopsis occidentalis s.l.* differs from *P. jackschultzi* at numerous mitochondrial and nuclear markers [35]. *P. jackschultzi* also co-occurs with all-female hybrid forms (biotypes) of *Poeciliopsis* that arose and persist via a peculiar asexual mechanism known as hybridogenesis [36, 37]. Hybridogenetic *Poeciliopsis* are effectively “perpetual” diploid F1 hybrids, whose maternal genome (derived originally from a *P. monacha* female) is inherited in a clonal manner, and whose paternal genome is “borrowed” from a sexual species (the sexual host) each generation and discarded during oogenesis via an abnormal meiosis (Fig. S1). The eggs of these hybrids carry one set of chromosomes and mitochondria derived from *P. monacha* and can be fertilized by sperm from distant sexually reproducing congeners of *P. monacha*, including *P. occidentalis s.l.* (producing the biotype known as *P. monacha-occidentalis*) and *P. jackschultzi* (producing the biotype known as *P. monacha-jackschultzi*) [29, 38]. Because half of their genome is inherited in a clonal fashion (i.e., the maternal genome), hybridogenesis is also known as hemiclonality. Because both the maternal and paternal genomes are expressed, these hybrids exhibit traits from both progenitors, effectively rendering them morphologically indistinguishable from their sperm donor species. The maternal progenitor of all *Poeciliopsis* asexual hybrids, *P. monacha*, is a sexually reproducing species whose range does not overlap with *P. jackschultzi* but does overlap with *P. occidentalis* in the Rio Mayo (i.e., the southern limit of *P. occidentalis* and the northern limit of *P. monacha*), where presumably the hybrids arose [39]. Because hybridogenetic *Poeciliopsis* carry *P. monacha*-derived mitochondria, mitochondrial markers can be reliably used to distinguish asexual hybrids from their sympatric sexual hosts (e.g. *P. jackschultzi* and *P. occidentalis*).

The primary goal of this study was to evaluate the status of *P. jackschultzi,* by surveying the condition of freshwater habitats and documenting the fish taxa present (native and non-native), within its known range and in nearby localities and tributaries. The following features of this study system and goals suggested that eDNA metabarcoding would be suitable as the primary method to survey fishes: (1) availability of a primer set [MiFish; 22] and a relatively complete reference library targeting a mitochondrial genetic marker that is diagnostic for all/most fish taxa expected to be present in the study area (*P. jackschultzi* is 3/167 bp different from its closest sympatric relative *P. occidentalis*; see Results); (2) inadequacy of external traits to diagnose *Poeciliopsis* species and hybrid biotypes; and (3) our intent to minimize harm to native fish populations, most of which have some degree of protected status (Table S1).

A secondary goal of this study was to document teleost species that are currently present in the Mexican stretch of the Rio Santa Cruz (Fig. 1), a tributary of the Gila River (which is a tributary of the Colorado River). The entire Colorado River has been considered outside of the Asexual Hybrid Topminnow’s native range [40], but recent surveys of the US side of the Santa Cruz, downstream of the US-Mexico border, have recorded its presence since at least 2017 [41], hypothesized to reflect a recent human introduction from a yet unknown Mexican location.

An additional goal of this study was to search for evidence consistent with the existence and presence of an undescribed cienega-microendemic chub (genus *Gila*), sympatric with *P. jackschultzi*. According to DeMarais *et al.* [42] and Hendrickson and Juárez-Romero [32], samples taken between 1971 and 1983 from the same cienega sites preferred by *P. jackschultzi*, contained a chub morphologically distinct from *G. ditaenia* (from the Rio Concepcion)*, G. purpurea* (from the Rio Yaqui), *G. eremica* (from the Rios Sonora and Matape), and *G. intermedia* (from the Gila River). This distinct chub, which is currently considered at native undescribed species (hereafter *Gila sp.*; [43]) was initially thought to represent *G. purpurea* introduced from the Rio Yaqui and subsequently referred to as *G. eremica* (i.e., the species it most closely resembles; [42]). A subset of specimens from these samples, which exhibited intermediate phenotype between *Gila* sp. and *G. ditaenia*, were deemed hybrids [42]. No reference sequence is available for this putative cienega-endemic chub, but its MiFish haplotype is expected to be distinct from *G. ditaenia*, and possibly more similar to those of *G. eremica*.

## 3. Materials and Methods

### 3.1 Study Area

Metadata for this study has been documented following [44] under Dataset S15. Because our primary goal was to monitor the presence of *P. jackschultzi*, most of the sampling efforts targeted sites within its known distribution (i.e., the Alisos-Bambuto branch and the lowest part of the Cocóspera branch) of the Río Concepción (between the border city Nogales and the town of Ímuris; Fig. 1). Additional sampling localities included other parts of the Cocóspera branch, as well as another tributary of the Río Concepción (Rio Altar), and one site in the Rio Santa Cruz, a few kilometers upstream from the US-Mexico border (Fig. 1). Unless otherwise noted, distances between sites are measured “along the river” based on tracing the path within the GoogleEarth Pro software application.

### 3.2 Sample collection

Prior to field work, sample collection kits were prepared in a “clean” lab (i.e., a room dedicated to pre-PCR processing of eDNA and ancient DNA samples; see Protocol S1A). The filter apparatus consisted of an analytical filter funnel (Cat# 28198-861; VWR) in which we replaced the original filter with a Whatman glass microfiber filter (1.5um particle retention; Fisher Scientific Cat# 09783DD). Each sample kit also included gloves, plastic zip bags containing desiccant beads (e.g. Orange Indicating Silica Gel Bulk Desiccant Beads, Industry Standard 2-5mm from Interteck Packaging) for storing the filter after collection, and sterile forceps (Cat# 76501-426; VWR).

Upon arrival at a sample site, and before any team member or gear was introduced into the water, one team member handled the filter funnel using the gloves contained in the corresponding kit. The filter funnel was connected to a tubing that connected to the pump (Geotech Geopump II Peristaltic Sampling Pump w/ Pump Head, AC/DC Power Accessories & Case, 91352113, Geotech Environmental Equipment, Inc.). On the effluent end of the pump, pumped water was collected into a graduated bucket to monitor and record filtered volume. At most sites, the first sample to be collected was a field negative control, referred to as “blank”, which consisted of pumping 30 seconds of air, rather than water, through a filter. For each collection site/time, we collected 2–3 filtered water replicates, each with an individual kit. For each replicate, we aimed to pump a maximum of 5L of water through the filter, but filter volumes tended to be lower because filters would usually clog before the maximum target volume was achieved (see Results). To minimize contamination from footwear and gear, at sites where there was a current, the mouth of the filter holder was placed facing upstream, whereas the person handling the filter stood downstream. At sites without a current, the person collecting the sample stood as far as possible from the filter holder, and where feasible, the filter holder was attached to a long pole, so as to avoid entering the water. After filtering the blank or each replicate’s volume, sterile forceps were used to remove the filter from the holder, and to fold the filter once or twice with the collecting surface facing inward. The folded filter was placed inside a bag containing the desiccant beads, and stored at room temperature while traveling, and at −80°C upon arrival to the lab.

### 3.3 DNA isolation

Samples were processed in two separate batches; one in 2022 and one in 2023. Batch 1 included all 2022 collections except one sample (i.e., MV22-01 Blank; Cienega Rancho El Aribabi), whereas Batch 2 included MV22-01 Blank plus all the 2023 collections (Table S2, Dataset S1, and Dataset S15).

The DNA extractions and set up of first-round PCRs were performed in the aforementioned clean lab. To isolate DNA, we used half of each filter (filters were cut with a sterile razor in a sterile Petri dish). The other half was returned to the respective sample bag and placed back into −80°C storage. The filter half was placed into a Qiagen Investigator Lyse and Spin column (Cat # 19598) with 475 ul ATL buffer and 25 ul of Proteinase K, and incubated overnight for ∼14 h at 55°C, in a Multi Therm Heat Shake (Benchmark Scientific) on the lowest shake setting. After incubation, we centrifuged the columns containing the filters. The resulting volume was processed with Qiagen’s DNeasy Blood & Tissue Kit following the user’s manual protocol for muscle tissue extraction. We performed two DNA elutions, each in a volume of 100 ul AE buffer that were pooled into the same tube, and processed to with the Zymo’s OneStep PCR Inhibitor Removal kit (Cat # D6030) following the user’s manual.

### 3.4 First-round PCR

We performed two rounds of PCR. The first round used the MiFish-U primers [22], which target a ∼180 bp fragment of the 12S rRNA mitochondrial gene. For each sample replicate and blank, we performed 4 replicate first-round PCR reactions (reagents, volumes, and thermocycler settings are indicated in Protocol S1B). The Platinum SuperFi II Green MasterMix (Life Technologies) was used because it is reported to reduce annealing bias [48]. After PCR, reactions (3 ul mixed with GelRed; Cat#41003; Biotium) were electrophoresed on a 2% agarose gel (Tris-Boric Acid-EDTA; TBE) and visualized under UV light. Successful PCR amplification runs fulfilled the following two requirements: (1) lack of evidence of amplification of the PCR master mix negative control; and (2) positive amplification of a positive control (i.e., template known to work under such PCR conditions). Any PCR amplification runs that failed at least one of the above criteria were discarded and repeated. Each of the four replicates per sample first-round amplification were subjected to the second-round PCR (see below).

### 3.5 Second-round PCR

The second round used the same MiFish-U primers to which unique “barcodes” (10bp unique index sequences) had been added on the 5’ end (Table S2 and Dataset S1). This dual unique indexing enables pooled sequencing and subsequent bioinformatic separation (i.e., demultiplexing) of the individual samples according to their unique barcode sequence. Within each of the two batches, each sample and blank was assigned a different pair of barcoded primers (Table S2 and Dataset S1). Each of the four first-round PCR replicates was used separately as template (undiluted) for four second-round PCR reactions (reagents, volumes, and thermocycler settings are indicated in Protocol S1C. The same success criteria as noted above for the first-round PCR were applied to the second-round PCR. In Batch 1, we used DNA from one of several *Gambusia affinis* specimens as the positive control for PCRs. However, because *Gambusia* was present as an invasive taxon in many of our sites, and thus could serve as a source of contamination for our experiments, for Batch 2, we switched to DNA from a *Heterandria formosa* specimen as the positive control for PCRs (see Table S2 and Dataset S1); all other conditions were identical between the first and second batches.

### 3.6 Post second-round PCR protocols

The four replicate 2nd-round PCR reactions per sample were pooled in equal volumes, and then purified with Zymo’s DNA Clean & Concentrator-5 kit (Cat # D4013). DNA concentration was measured with Qubit’s High Sensitivity DNA assay (Cat # Q33230). For each batch separately, equimolar amounts of each sample were then pooled into a single tube. For the field negative controls (samples collected as “blank”) and the single extraction negative control, which are generally expected to yield low or no amplicons, we used 8 ul, reflecting the average volume used from the other samples. The tube of pooled samples was shipped on dry ice to Genewiz, Inc., where it was subjected to purification with AMPpure (Beckman Coulter) to size-select DNA to enrich the amplicons for the target size. A single library (2 x 150 bp paired-end layout) per batch was prepared and sequenced on an Illumina platform (HiSeq 4000 for Batch 1 and MiSeq for Batch 2).

### 3.7 Bioinformatics Analyses

Details on Bioinformatics analyses and associated files can be found in Protocol S2. Because some of the samples in Batch 2 shared the same barcodes as those in Batch 1 (Table S2 and Dataset S1), initial trimming and demultiplexing of the raw sequence reads received from GeneWiz, Inc. was performed separately on each batch, with two rounds of Cutadapt v3.5 [49] and concatenated (Protocol S2). The concatenated files from batch 1 plus batch 2 were used as input for the QIIME2 pipeline [50] implementing the parameters found in Protocol S2. The main output of the above QIIME2 pipeline is a set of “features” (also known as Amplified Sequence Variants; ASVs), which are expected to represent actual DNA sequences present in the sample, along with their estimated frequencies in each sample. We used two complementary approaches to assign each feature to a taxon. One approach used blastn [51] against NCBI’s non-redundant database, as implemented in Geneious Prime® v2023.2.1 to extract the top hit. Using the species identification of the top hit, the features were separated according to whether they were identified as teleosts vs. non-teleosts (i.e., Bioinformatics Filter #1).

We aligned the teleost-only features, plus the sequence of their blastn top hit(s), as well as publicly available and newly generated (Table S3) MiFish region sequences of teleost species we expected to find in the study area. Distance-based trees (e.g. uncorrected p; Neighbor-Joining) were used to verify that features fell within the clades of the species name that matches their top blastn hit (see Results). Those that did not were further scrutinized by checking additional blast hits and visually examining the sequence within the alignment. Features with substantial indels and/or of apparent chimeric origin (e.g. resembling one species on one end, and another species on the other end) were discarded (i.e., Bioinformatics Filter #2). All discarded features, along with the criteria used to discard them, are in Dataset S3. Retained teleost features were used to compute the collective eDNA abundance (adding the frequency of all features assigned to a particular species) per species per sample replicate and subjected to Bioinformatics Filter #3, which aimed to account for contamination based on the results of negative controls (see Results). Relative and absolute frequencies of species were plotted in the Microsoft Excel® software.

The above QIIME2 pipeline, which is a de novo method, could potentially discard sequences incorrectly flagged as chimeric [reviewed in 52]. To verify that the absence of *P. jackschultzi* features revealed by the QIIME2 pipeline (see Results) reflected absence of *P. jackschultzi* reads, we mapped to the known sequence of *P. jackschultzi*, the post-Cutadapt reads from the sites 3–10 (Fig. 1A), i.e., the subset of sites within or nearby the known range of *P. jackschultzi* (see Protocol S2).

## 4. Results

### 4.1 Assessment and mitigation of contamination and method-induced artifacts

We received 511,987,600 and 1,157,193,283 sequence read pairs (hereafter reads), for Batch 1 and 2, respectively. All 73 samples, including one extraction negative control and 18 field negative controls (i.e., blanks; ratio of field blanks to field samples was 18.1% = 13/72), yielded reads after the Cutadapt filtering/trimming step. Collectively, a total of 812.3 million reads were retained after the Cutadapt filtering/trimming step, and subjected to the QIIME2 pipeline (Table S2 and Dataset S1), which retained 535.8 million reads. With few exceptions, lab and field negative controls yielded very few reads. Excluding negative controls and one outlier sample (i.e., La Mesa, MV23-2#2; site 10; which only yielded 124 reads), QIIME2-retained reads per sample averaged 10 ± 4.8 (std. dev) million (range: 0.48–22.5 million; Table S2 and Dataset S1). The sum of QIIME2-retained reads from sites within (or nearby) the known range of *P. jackschultzi* was ∼348 million.

QIIME2 generated 2,925 unique features (Dataset S2), of which 2,774 had a teleost taxon as their top blastn hit (i.e., they passed Bioinformatics Filter #1). Among the 151 features that failed to pass Bioinformatics Filter #1, 80 were assigned to cattle (*Bos* spp.), which collectively accounted for the most abundant non-teleost features in our samples (> 8 million). This finding is not surprising because cattle were abundant in the study area and the 12S rDNA gene sequence of *Bos taurus* (AY526085.1) has a perfect match for the MiFish-U-R primer and a high match (two mismatches) with the MiFish-U-F primer, expected to yield a 216 bp amplicon. The remaining 71 non-teleost features included blastn hits to reptiles, other mammals, birds, bacteria, and viruses (Dataset S3).

The sequences of the 2,774 teleost features were separated into datasets by blast-assigned taxonomy (e.g. Poeciliidae, Leuciscidae [formerly Cyprinidae]) and aligned to their closest blast hit and other reference sequences (i.e., available sequences from the same or closely related taxa). Final assignment of features to a particular taxon (in most cases, a species) generally followed the >97% similarity threshold used by many studies, but also took into account whether the feature fell within the clade of the putative species in the NJ tree, and whether the MiFish region is known to be diagnostic for the putative species versus its close relatives. The preliminary NJ trees revealed several “rogue” features (i.e., those that did not group neatly within their expected clade; not shown). Upon further visual scrutiny in sequence alignments, we identified and removed 252 features with large indels or that appeared to be chimeras (i.e., Bioinformatics Filter #2).

The remaining 2,522 teleost features represented 71 putative species (see “counts_by_final_taxon” sheet in Dataset S3). Based on the taxon identity (perfect vs. imperfect sequence match to reference, marine vs. freshwater; likelihood of being present in study area) and its frequency patterns (summed over all samples > 6,000; and consistent occurrence across replicates), we filtered out 50 putative taxa; retaining 21 for further analysis (see “counts_by_final_taxon2_ro” sheet in Dataset S3). The 21 retained taxa were classified into native (N), introduced, non-native or invasive (I), and likely spurious (S). Native refers to taxa that are known to occur natively in the broader study area (i.e. Arizona-Sonora border area), even though they may not be native to the particular tributary or basin where its eDNA was detected. Similarly, introduced (I) are those taxa known to have been introduced into the study area from outside the broader region. Spurious (S) refers to taxa that are unlikely to inhabit the study area (e.g. field contaminants, lab contaminants, etc.), but were detected at substantial frequencies. The most abundant spurious taxon was the Monterey Spanish Mackerel; detected at one replicate of Agua Caliente MV23-13 (site 6) at very high abundance ∼8.8 million (magenta bar in Fig. S2). This marine pelagic species inhabits the Gulf of California, which is the nearest marine environment to our study area (Fig. 1B), and is widely used regionally in “ceviche” (raw fish dish treated with lime juice). This, along with the fact that, to our knowledge, neither Monterey Spanish Mackerel nor its close relatives have ever been processed in our laboratory’s building, leads us to suspect that it is a field, rather than a lab, contaminant (possibly reflecting discarded ceviche in the sampled water body).

The 2nd-most-abundant likely spurious taxon was assigned to the Eurasian Minnow or a congener (genus *Phoxinus*; abundance ∼800K; 99.9% of which was in the blank of Imuris-Bambuto sidestream MV22-5C; site 5; Fig. S3). We consider this detection a lab contaminant because it is very unlikely that its DNA would be present in the study area [see 53], it had a perfect match to two sequences in NCBI (i.e., unlikely to be the result of a method-induced mutation; see below), and because other labs in our building work with closely related leuciscids. Five of the remaining six spurious taxa appear to be lab contaminants, because four (three leuciscids and one centrarchid) were almost exclusively recovered from the Aribabi Cienega 2022 blank (abundance range ∼ 7–68K; site 2), and one leuciscid (Stone Loach *Barbatula barbatula*; abundance ∼37K; Fig. S3) was almost exclusively recovered from the La Atascosa 2022 blank (site 7). Similar leuciscids and centrarchids have likely been processed within our building. The least abundant spurious taxon (assigned to Patzcuaro Chub *Algansea lacustris* by blastn; abundance ∼6K), is likely a method-induced artifact (e.g. mutation or chimera) of the highly abundant leuciscids in our sample (e.g. Longfin Dace and Sonora Chub), as it was more evenly distributed across the 2022 and 2023 samples (31 out the 73 samples; abundance range per sample 2–1,067; mean = 199; median = 115), and did not perfectly match its closest blastn hit. Hereafter, the features and frequencies from the aforementioned spurious taxa were excluded from downstream analyses, leaving six native (N) and seven introduced (I) taxa represented by 2250 (distinct) features whose collective abundance is 515,931,851.

The number of features per each of the thirteen non-spurious teleost species varied widely (mean = 173.08 ± 118.09 S.D; range 1–337). As observed in the frequency histograms (Fig. S4), the frequency of the top 1–2 most abundant features assigned to each species was substantially higher than the remaining features assigned to that species (i.e., typically accounting for > 95% of the cumulative frequency). In addition, the top 1–2 most abundant features per species typically had a perfect match to the available reference sequence(s). The remaining (less frequent) features within each species typically differed from the reference or the most abundant feature(s) by one nucleotide substitution (e.g. Fig. S3 and Fig. S5). Furthermore, a positive (exponential) association is observed between the number of features assigned to a species and their collective frequency (Fig. 3). For example, the Longfin Dace and Sonora Chub had the highest eDNA abundances (>150 million each) and the highest number of features (337 and 325, respectively). The observation that species with higher inferred eDNA abundance had a higher number of unique features can be explained by two hypotheses: (1) that the majority of the low frequency features are not artifactual and thus, species with a higher abundance of eDNA harbor higher genetic diversity, possibly reflecting larger effective population sizes; or (2) that the majority of the low frequency features reflect artifactual single-base mutations that occurred during the PCR or library preparation procedures. Support for the latter hypothesis is provided by Illumina metabarcoding analyses of known bacterial “mock” communities [27], which demonstrated that most sequence variants inferred in a sample are artifacts, that erroneous variants accumulate at a roughly constant rate as a function of the number of Illumina reads, and that convergence is not achieved in rarefaction curves. Because with the currently available data, it is impossible to confidently determine which of the lower frequency features are artifacts, we did not discard features based on their frequency. Retaining artifactual features resulting from method-induced ∼1–2 bp substitutions should not bias our inferences of a species’ eDNA abundance, but we refrain from using them to make inferences about within-species genetic diversity, unless other evidence supports such inferences (see below).

We excluded the two replicates and the blank from Aribabi Cienega 2022 (MV22-1) because of the high abundance of teleost features (native, introduced, and spurious) in the blank, and because of very different relative abundances in the two replicates (see Fig. S2). The remaining 70 samples and blanks were deemed “acceptable” because after removal of the spurious taxa, the relative frequencies of the replicates within each site/year were similar to each other, and the blanks had a very low abundance of teleost taxa.

To account for contamination detected in the remaining negative controls (17 field blanks and 1 extraction negative control), we applied the following filter (Bioinformatics Filter #3) to the remaining 52 non-negative-control samples. For each of the 13 non-spurious teleost species, we identified the negative control with the highest abundance (hereafter “all blanks threshold”), which ranged from 0 to 1,293. We then converted to zero the abundance of the corresponding species for every sample that did not have an abundance above the “all blanks threshold” (see “counts_by_final_taxon2_ro” sheet in Dataset S3).The resulting corrected abundance values (13 species and 52 non-negative controls) were used to compute the sum of replicates per site/year (see “cnts_by_final_taxon2_ro_no_blnk” sheet in Dataset S3). Sum of replicates per site/year > 0 were classified as “effectively present”, whereas those below as “effectively absent”. These values were also used to plot the relative frequencies of each taxon per site/year (Fig. 2).

**Figure 2.**
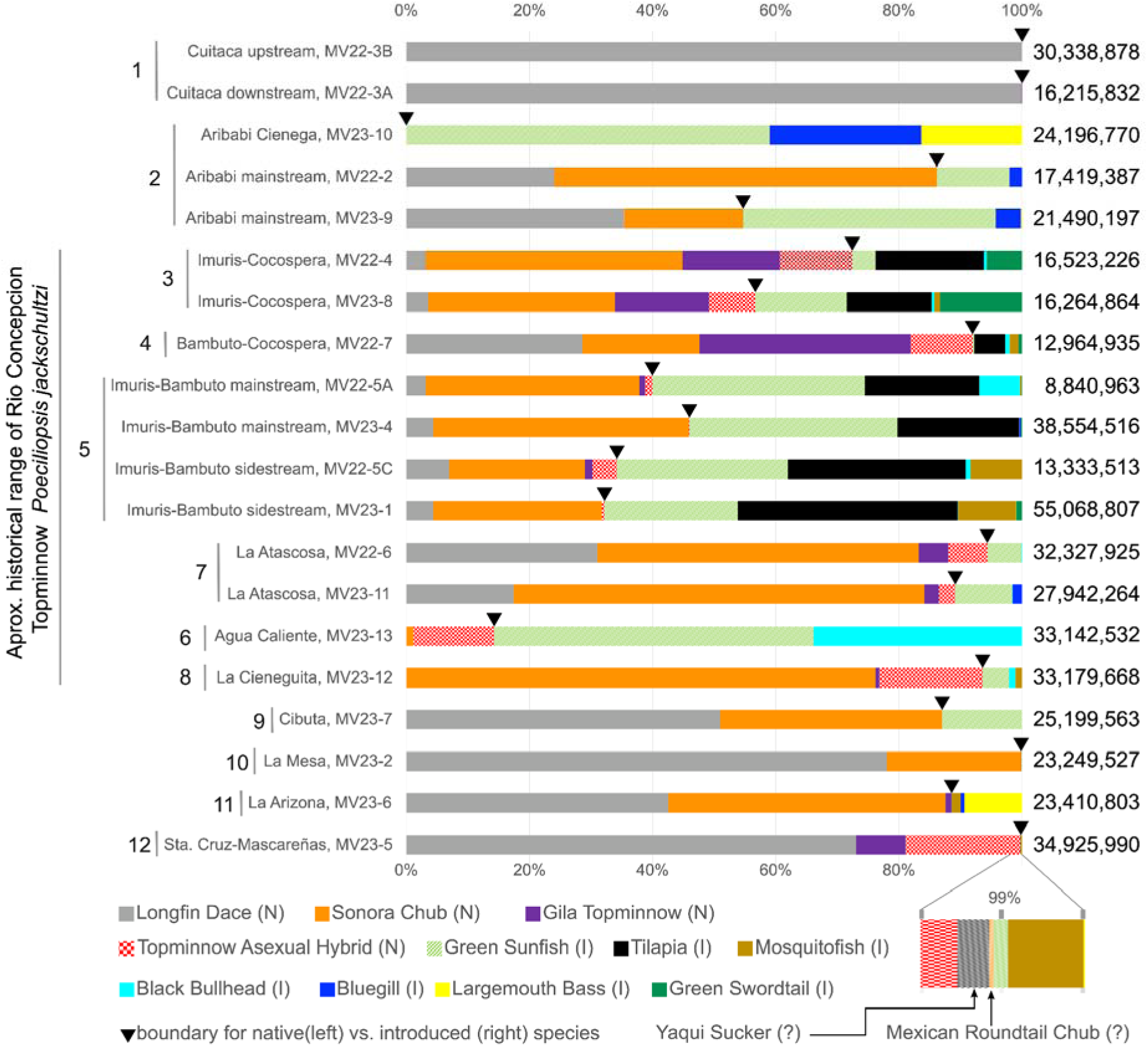
Relative eDNA frequencies per site/year. Relative eDNA frequencies of the thirteen non-spurious teleost taxa per each site/year sample (summed over replicates); excluding blanks and Aribabi Cienega 2022. Numbers to the left of the site/year labels correspond to the site numbers of Fig. 1A. Sites 1–11 are Rio Concepción. Sites 1-2 are Cocospera above the Comaquito dam. Site 3 is Cocospera below Comaquito dam. Site 4 is the confluence of the Cocospera and Bambuto tributaries. Sites 5–10 are considered within the Bambuto tributary. Site 11 is the only Altar tributary sample. Site 12 is the only Gila River (Mexican side of the Río Santa Cruz) sampled. Numbers to the right of each of the depicted 20 samples indicate the collective abundance (= frequency) used to estimate the depicted relative frequencies. The inverted black filled triangles indicate the proportion of native (to the left) vs. introduced (to the right) species. For the Santa Cruz sample, the 98-100% region is expanded to show the relative frequencies of the Yaqui Sucker (0.08%) and the Mexican Roundtail Chub (0.01%). N = native; I = introduced;= native to the Arizona-Sonora border region but not to the site where it was detected (see main text). Original graph generated in sheet “cnts_by_final_taxon2_ro_no_blnk” in Dataset S3, and edited with Inkscape. A version of this figure showing each replicate separately is Fig. S11.

As recommended by Goldberg *et al.* [54], rather than using a strict dichotomous approach to determining presence/absence of a taxon at a site/year based solely on whether their eDNA abundance was above/below an established threshold (i.e., whether they were “effectively” present/absent as defined above), we considered the strength of the evidence. Such evidence included the eDNA abundance (higher abundances = higher strength of evidence), but also assumed that taxa whose collective abundance among samples was high, were more likely to serve as contamination sources for other samples, leading to false positives or inflated abundances. We also considered whether the taxon was visually detected (based on individuals captured in seine or minnow trap, or simply observed in the habitat) at the time of eDNA collecting (i.e., “Observed”), and whether it is expected based on: (1) previously recorded at the site; or (2) previously recorded at a nearby site from which individuals are likely to have colonized (e.g. upstream or downstream from a known site) or eDNA is likely to have traveled (e.g. from an upstream to a downstream site). Previous records were based on our own observations (e.g., fieldnotes of Varela-Romero, Mateos, and R.C. Vrijenhoek since ∼1980’s), the “SONFISHES” database [55], peer-reviewed literature [e.g. 32, 33], and the GBIF database [56]. The collective evidence was then used to assign a species as absent (A) vs. present (P) under the “inference” category (“Inf” in Table S4). Out of 260 cells (13 species x 20 sites/years), three eDNA inferences were considered likely false positives (inference = Absent), and whereas three others were deemed potential false negatives (inference = Present; Table S4).

### 4.2 No evidence of Rio Concepcion Topminnow (*Poeciliopsis jackschultzi)* or the putative cienega-endemic Rio Concepcion Chub

None of the features recovered by the QIIME2 pipeline had a perfect match to the only known MiFish haplotype of *P. jackschultzi* (based on 5 specimens), which differs from known haplotypes of *P. occidentalis* and *P. monacha* at 3 and 13–14 nucleotide positions (out of 167), respectively. To further verify the absence of *P. jackschultzi* eDNA in our recovered sequences, we mapped the post-Cutadapt raw reads from localities 3–10 to *P. jackschultzi* reference sequence (Protocol S2). This procedure also did not recover a single read pair that perfectly matched the sequence of *P. jackschultzi*, further confirming that none of our recovered sequences derive from *P. jackschultzi* eDNA.

None of the eDNA features recovered were sufficiently distinct from the Sonora Chub (*G. ditaenia*), and geographically restricted to the Rio Concepcion to suggest that they represent the cienega-endemic chub that was presumably recorded in these habitats (e.g. our La Atascosa and La Cieneguita sites) between 1971 and 1983 [32, 42].

### 4.3 Patterns of native and introduced teleosts eDNA abundance per site/year

The relative eDNA abundance of native (N) to non-native (I) teleosts varied widely among sites/years (mean= 69 ± 32% SD; Fig. 2 and Table S4) from exclusively natives (Longfin Dace) in the Cuitaca (most upstream Cocospera) sites to exclusively non-natives at Aribabi Cienega (Cocospera). The native Longfin Dace was the most geographically widespread teleost, as it was only effectively absent from three sites, which had lentic characteristics (i.e., La Cieneguita and Agua Caliente in Alisos-Bambuto tributary, and Aribabi Cienega in the Cocospera). Its eDNA relative abundance where/when present ranged = 3.1–100% (mean = 35.7 ± 33.7% SD; Table S4). The Longfin Dace total eDNA abundance (∼150 million) was just slightly lower than that of the native Sonora Chub (∼153 million; Fig. 3). Sonora Chub eDNA was effectively present at all sites except for the Rio Concepcion upper Cocospera sites (Cuitaca upstream and Aribabi Cienega), where the collective evidence also suggests it was absent (i.e., inference = Absent). Based on our strength of evidence criterion, we consider that the detection of Sonora Chub eDNA at the Cuitaca downstream and the Rio Santa Cruz sites is probably the result of contamination (i.e., false positive; thus, inference = Absent; Table S4). The Santa Cruz (and the entire Colorado River) is considered outside of the native range of the Sonora Chub. The Sonora Chub eDNA relative abundance where/when present (excluding the records deemed false positives) ranged 1.2–76.2% (mean = 37.3 ± 19.8% SD).

**Figure 3.**
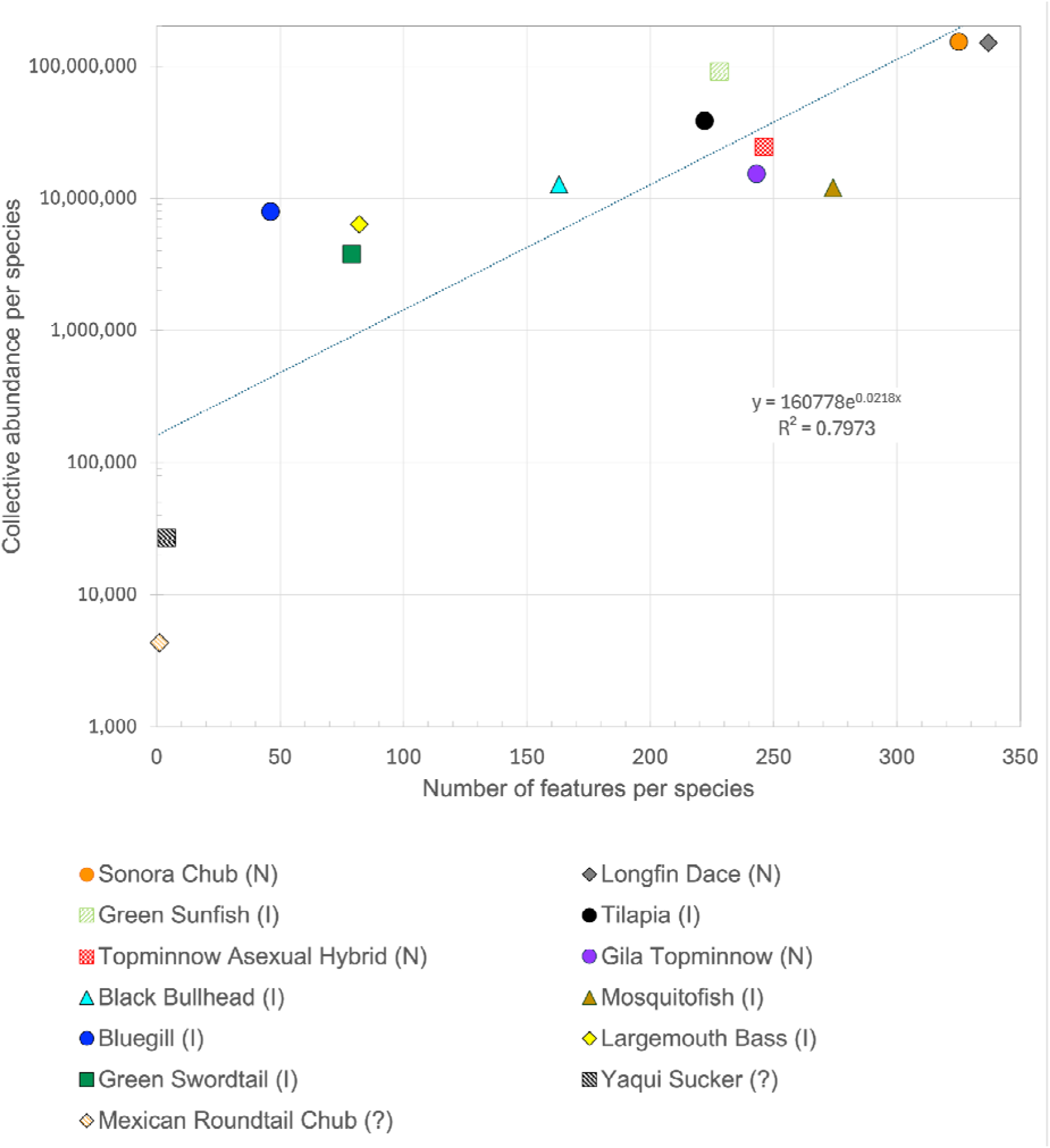
Feature abundance vs. feature frequency. Feature abundance vs. feature frequency for the thirteen non-spurious teleost taxa and linear regression trendline. The Y-axis is depicted in logarithmic scale (base 10). Data used and original graphs found in sheet “features_passed_filters1&2_cnts” of Dataset S3. N = native; I = introduced;= native to the Arizona-Sonora border region but not to the site where it was detected (see main text).

The Green Sunfish was the 3rd-most frequently detected teleost (∼90 million) and the most geographically widespread non-native, as it was effectively detected in all sites/years except for Cuitaca and La Arizona (eDNA relative abundance where/when present range: 0.03–59%; mean = 19.6 ± 18.7% SD; Table S4). Its congener, the Bluegill (∼6 million) was detected at only six site/year Rio Concepcion samples (eDNA abundance range 0.26–24.6% mean ± SD = 5.5 ± 9.45%), of which all except La Arizona yielded also Green Sunfish eDNA, which was consistently at a higher abundance relative to that of the Bluegill.

The 4th-most frequent teleost detected, the Africa-native Cichlidae to which we collectively refer to as Tilapia (∼40 million), had a somewhat geographically restricted distribution; detected at all seven site/year samples along the stretch spanning the lower Cocospera to the lower Bambuto, including their confluence just below the town of Imuris (relative eDNA abundance range = 5.6–35.7%; mean = 19.9 ± 10% SD). Surprisingly, whilst eDNA from Tilapia was effectively absent at La Arizona, we inferred Tilapia as present (Table S4) because it was captured in a seine haul.

The non-native poeciliids Green Swordtail (∼4 million; max relative frequency = 13.3% at Imuris-Cocospera 2023) and Mosquitofish (∼7 million; max relative frequency = 9.4% Imuris-Bambuto sidestream 2023) were effectively detected at most site/years at which Tilapia eDNA was detected. Mosquitofish appears to have a broader distribution than the Green Swordtail and Tilapia, as we detected its eDNA (albeit at relatively low frequencies) at sites more upstream in the Bambuto, at La Arizona (Rio Altar, a separate tributary of the Rio Concepcion), and at the Rio Santa Cruz. The non-native Black Bullhead was also detected at most sites/years that harbored Tilapia (max relative frequency in this stretch = 6.6%), but it was also detected in the two lentic sites of the Bambuto, reaching its highest relative frequency (33.9%) at the Agua Caliente site. Of the non-native teleosts, Largemouth Bass accounted for the lowest eDNA abundance (∼3 million), and was detected at only four site/years of the Rio Concepcion; it was most abundant at Aribabi Cienega 2023 (16.4%), followed by the geographically distant site of La Arizona (9.3%).

The fifth and sixth-most frequent (respectively) teleosts were the Topminnow Asexual Hybrid (∼19 million; relative eDNA abundances range = 0.05–18.6%; mean = 7.7 ± 6.6%) and its sexual host the Gila Topminnow (∼15 million; relative eDNA abundances range = 0.03–34.2% (mean = 7.04 ± 10.3%), which are both considered native to the Rio Concepcion. With one exception that we considered a false positive eDNA detection (i.e., Gila Topminnow at Cuitaca downstream site), both were effectively (and inferred) absent from all Cocospera sites upstream of the Imuris site. One or both of these topminnows were inferred present in the remaining sites of the Concepcion, because *Poeciliopsis* sp. was visually confirmed albeit eDNA not effectively present (e.g. Cibuta), or because eDNA was effectively present. The highest relative abundance of Gila Topminnow eDNA was found in the Bambuto-Cocospera confluence followed by the Imuris-Cocospera site. The highest relative abundances of Topminnow Asexual Hybrid eDNA in the Concepcion occurred at the two lentic sites (La Cieneguita = 16.75%; and Agua Caliente = 13.1%). Topminnow Asexual Hybrid eDNA was effectively absent from the most upstream sites of the Bambuto (Cibuta and La Mesa) and from the single Altar tributary site sampled (La Arizona). In the Rio Santa Cruz, the relative eDNA abundance of Gila Topminnow (considered native) was 8.1%, whereas that of Topminnow Asexual Hybrid (considered non-native) was 18.6%, which was the highest relative frequency of this taxon among all the site/years samples.

Two additional teleosts, a sucker (Catostomini) and a chub (*Gila*) that is not Sonora Chub were detected, exclusively in the Rio Santa Cruz, albeit at low abundances. The most abundant sucker feature (i.e., 6e7ec8e5e4e6637ea39b5e01a542e824; frequency = 25,145; relative freq = 0.08%; Fig. S6) was identical to the only known haplotype of the Yaqui Sucker *Catostomus bernardini* (OL878383; from site RV in (Fig. 1B), which is ∼700km away straight-line in the Rio Fuerte basin), and different (at 3/170 and 12/170bp, respectively), from the only known native suckers to this part of the Gila River: the Sonora Sucker *Catostomus insignis* (sampled from site AC in the San Pedro River; Fig. 1B) and Desert Sucker *Pantosteus* (formerly *Catostomus*) *clarkii* (sampled from sites AC and SC in the Santa Cruz; Fig. 1B). The most abundant eDNA feature of the chub that we exclusively found in the Rio Santa Cruz (i.e., d06fd679e55d16876f12fe5b88d61436; 4,317; 0.01%) was identical to the only haplotype known from Mexican Roundtail Chub *Gila minacae*, sampled from the Rio Yaqui site AL (Fig. 1B; ∼270km away straight-line), and was different from: (a) all available members of the *Gila robusta* complex (for taxonomic status see [57–60]); (b) the Yaqui Chub (*Gila purpurea*); and (c) the Desert Chub (*Gila eremica*) and its close relatives from Sonora and Sinaloa (Mexico) (Fig. S3).

### 4.4 Inferences about intraspecific variation at the MiFish region

#### 4.4.1 Asexual Hybrid Topminnow

For the Asexual Hybrid Topminnow, we have independent evidence that two MiFish haplotypes exist in the Rio Concepcion. Their diagnostic MiFish SNPs are linked to Cytochrome b (cytb) SNPs previously used to distinguish two *P. monacha*-derived mitochondrial haplotypes referred to as “A” and “D” (GenBank AF047344 and AF047343, respectively). A previous study [61] and our own data (unpublished) revealed that cytb haplotype A is more common and geographically widespread (present in all rivers from the Mayo to the Concepcion, and recently found in the Rio Santa Cruz), than haplotype D, which was only known from the Rio Concepcion (see Fig. S5). Our eDNA results revealed that haplotype A was the most frequent Asexual Hybrid Topminnow feature (collective abundance = 18,675,350), followed by haplotype D (5,365,477). Nonetheless, the relative frequencies of the two haplotypes varied among sites, with haplotype A being more common in all sites/years except for Imuris-Cocospera, where the frequency of haplotype D was consistently higher in all samples from both 2022 and 2023 (range 58.3–77.24%; Fig. S7). We hypothesize that most or all of the remaining 244 features assigned to Asexual Hybrid Topminnow (Fig. S5) reflect method-induced artifacts of haplotype A or D, including the three low-frequency features (absolute frequency range = 227–4164) that differed at 1 bp from haplotype A and perfectly matched the following samples from geographically distant localities: (a) a sexual *P. monacha* (GenBank NC_032074) and a hybridogenetic *P. monacha-lucida* (strain Cw S68-4) from the Rio Fuerte; (b) a *P. monacha-occidentalis* from the Rio Mayo; and (c) a gynogenetic triploid *P. monacha-lucida-lucida* (strain M61-31) from the Rio Fuerte [62].

#### 4.4.2 Gila Topminnow

For the Gila Topminnow, we did not have any independent evidence of the existence of more than one MiFish region haplotype (GenBank Acc. Nos. NC_028294.1 and MK860198), which is shared by all members of *P. occidentalis s.l.* examined to date, including the Yaqui Topminnow *Poeciliopsis sonoriensis* (MK860197) [63]. As expected, the most frequent Gila Topminnow feature (collective abundance = 14,487,940) perfectly matched the reference sequence. However, the 3rd-most frequent Gila Topminnow feature (f27c8509b944165b2eb1cc8a4dd1e416; collective abundance = 44,792; Fig. S5 and Fig. S8) was exclusively and consistently, detected in the five samples (2 from 2022 and 3 from 2023) from La Atascosa (site 7). Its frequency, relative to the two most frequent Gila Topminnow features at these five La Atascosa samples, ranged 0.9–4.3% (mean = 2.3 ± 1.2% SD). Such a non-random pattern of geographic/temporal distribution strongly suggests that the Gila Topminnow 3rd-most frequent feature represents a real haplotype found at relatively low frequency at La Atascosa. Because the 2nd-most frequent feature (collective abundance = 94,301; Fig. S5 and Fig. S8) did not have such an obvious non-random geographic pattern, it is unclear whether it is a real haplotype or whether it reflects a common method-induced mutation.

#### 4.4.3 Longfin Dace

The most abundant eDNA feature assigned to Longfin Dace (bbda9ad3bc84c50cfb5eff67c91d825a; 146544257; 69 samples; (Fig. S3) was identical to two Longfin Dace reference haplotypes: one from Rancho El Aribabi (Acc. No. PX746530) and one from New Mexico (AF081838; San Francisco River), and 4/176bp different from a Longfin Dace from an unknown locality (AY216528). Two eDNA features assigned to Longfin Dace exhibited a non-random pattern of presence: 3e74807cf24c44fd159109e41cd83625 (collective abundance 82,914) was exclusively recovered in the three Santa Cruz samples (Fig. S3); and 97.3% of the abundance of 05712217e29b0bc70fc2b6fa766044ca (i.e., the 2nd-most frequent Longfin Dace feature; collective abundance 442,060) was contributed by the three Rio Santa Cruz samples (Fig. S9; Dataset S7). The geographically biased presence of these features suggests that they might represent real haplotypes, rather than method-induced mutations. The possible existence of Santa Cruz exclusive/predominant haplotypes is consistent with the finding of significant and large *Fst* values (range 0.53–0.56) in comparisons from two Concepcion sites vs. one Santa Cruz site in/nearby our sampled sites, based on one nuclear (*Rag1*) and several mitochondrial (*Cytb*, *COI*, and *ND1*) markers [64].

#### 4.4.4 Sonora Chub

The most abundant eDNA feature assigned to the Sonora Chub (fa93e21884d734dc87b38f66d1dce538; collective abundance 148,092,536; 67 samples) had a perfect match to the reference haplotype from Rancho El Aribabi (Acc. Nos. PX746531 and PX746532), and 1 bp difference from a Sonora Chub specimen (AF038476) from a headwater site of the Rio Concepción Altar branch in Arizona (Sycamore Creek; see Fig. S3). A non-random pattern of presence and frequency was observed for the Sonora Chub’s 2nd-most frequent feature (7e54acc862386d9e6c1f9609ba0e0525; 1,389,879) at Aribabi Mainstream samples from both years (n=5), which collectively accounted for 70.5% of the abundance of this feature (Fig. S10; Dataset S7). This feature also reached its highest frequency among all sites/years at Aribabi Mainstream (range = 5.6–8.7% relative to the top four Sonora Chub features). Thus, feature 7e54acc862386d9e6c1f9609ba0e0525 may represent a real haplotype.

#### 4.4.5 Mosquitofish

For the Western Mosquitofish, we have independent evidence that, at least in its native range, two haplotypes exist that differ from each other at 5 out of 170 bp (uncorrected p = 2.94%). The most frequent Mosquitofish feature recovered (8fdf39f489879d13e0670f471f411c2a; 7,110,293) perfectly matched GenBank ID NC_004388 and a Sanger-sequenced specimen sourced from a stream in College Station, TX. We hereafter refer to this feature as “haplotype (hap) 1”. The 2nd-most frequent Mosquitofish feature (adbb5430e02b82ee5b2b181409c44034; 3,849,476), hereafter “hap 2”, perfectly matched two Sanger-sequenced specimens also sourced from a College Station stream. We are uncertain whether both Mosquitofish haplotypes are present in the study area for the following reasons. Firstly, one or more of the three aforementioned Sanger-sequenced College Station specimens were used as PCR positive controls for all the samples processed in batch 1 (i.e., all MV22 samples except for MV22-1 blank, from Aribabi Cienega). Consequently, large amounts of Mosquitofish MiFish amplicons were produced in our lab in 2022, which could have contaminated other samples. An encouraging finding, however, is that we did not detect a single feature attributable to *Heterandria formosa*, the positive PCR control used in samples processed in the 2023 batch. In other words, for the 2023 batch, we are certain that our sample handling practices, which were the same as those applied in 2022, prevented contamination from the positive control.

Secondly, 99.6% of hap 2’s abundance was contributed by samples processed in the 2022 batch, of which 99.9% was contributed by the two non-blank Aribabi Cienega 2022 samples (MV22-1.1 and 1.2; Fig. S12). The blank for this site/year (MV22-1blank), which was processed with the 2023 batch, had a large abundance of hap 1 (444,809) and ∼30X lower abundance of hap 2 (14,654). As mentioned above, due to the large abundance of teleost features in the MV22-1 blank, and to the discrepant relative abundances of MV22-1.1 and 1.2 (Fig. S2), the blank and samples from this site/year were considered unacceptable, and thus excluded from inferences of presence/absence of teleosts. In contrast, the 2023 samples from Aribabi Cienega (MV23-10blank, 10.1, 10.2, 10.3) had zero copies of hap 2 and extremely few of hap 1 (i.e., hap 1 absolute abundance = 6, 6, 10, 18, respectively), thus, effectively no evidence of Mosquitofish eDNA. Thus, most of the hap 2 abundance detected reflects either lab contamination, or its 2022-only presence at Aribabi Cienega. Unfortunately, we did not keep track of which of three Mosquitofish specimens (and thus which haplotype) was used as a positive control for each of the PCRs performed in the 2022-processed batch, which could have helped discern the source (i.e., present in the field vs. lab contaminant) of hap 2 features.

In contrast to Mosquitofish hap 2, hap 1 was recovered more consistently between samples collected and processed in both years, and its consistent recovery at high absolute and/or relative teleost abundance at the site where our visual surveys during both years confirmed the abundant presence of Mosquitofish (i.e., Imuris-Bambuto sidestream), increases our confidence that hap 1 is the only (or the predominant) Mosquitofish haplotype in the study area. The third-most frequent Mosquitofish feature (e35ea82ef6b0d6554d0ffc0682141185; abundance = 737,933) differs from hap 1 at one nucleotide position, and was only recovered where hap 1 was also recovered. These observations suggest that the third-most frequent Mosquitofish feature is either a real low-frequency haplotype present in the study area, or that it reflects a method-induced single base substitution of hap 1.

## 5. Discussion

The primary goal was to use eDNA metabarcoding to identify localities where the recently described, microendemic, endangered, and morphologically cryptic Rio Concepcion Topminnow (*P. jackschultzi*) is extant. Over two sampling expeditions (May 2022 and May 2023), we attempted to survey as many mainstream and tributary sites as possible within the known historical range of *P. jackschultzi* (i.e., between sites 3 and 8; Alisos and lower Cocospera), as well as nearby localities within in the same tributaries or in separate tributaries that have nearby headwaters. At least two sites where *P. jackschultzi* was recorded in the past were completely dry: the type locality, which is a small tributary that meets the mainstream immediately downstream of La Atascosa (site 7); and dry site II* (Fig. 1), which is a dead cienega. Our failure to detect evidence of eDNA from this species implies that it was truly absent or that its abundance in our samples was below the detection limits of our methodology. We consider that we can rule out PCR amplification biases against *P. jackschultzi* because it exhibits a perfect match to the primers used (GenBank PX582708), and the four individuals collected in 1986 that we subjected to the MiFish PCR and Sanger sequencing amplified successfully (GenBank PX556618-PX556621). We also have no reason to believe that *P. jackschultzi*’s eDNA is shed at a lower rate or is less stable than that of its sympatric congeners, which have similar size, life history, and ecology. Considering only the sampled localities that perfectly match or are geographically very close to localities where *P. jackschultzi* was historically reported (i.e., 3 and 5–8), and using the most stringent filtering (i.e., values shown in Fig. 2), we estimate that if present in our samples, the eDNA concentration of *P. jackschultzi* would be lower than 1 in 8,840,963–55,068,807 (i.e., the range of collective abundances of samples taken at sites 3 and 5–8), or lower than 1 in 275,178,278, if we consider the collective abundance of teleost eDNA in sites 3 and 5–8). The possible ways to increase our detection power using the same eDNA metabarcoding approach include: (a) increasing sequencing depth of the libraries; (b) increasing the number of PCR replicates per sample (we performed four PCR replicates); or (c) increasing the number of samples per site/year (we collected 2 or 3). Alternatively, a metabarcoding approach that uses primers specific to *P. jackschultzi*, or at least specific to *Poeciliopsis,* could increase the target taxon reads without requiring increased sequencing effort per sample. It is unclear to us whether the use of an *P. jackschultzi*-specific quantitative PCR assay (traditional or digital PCR) would enable detection of *P. jackschultzi* at concentrations as low as we predict (e.g. lower than 1 molecule in 275 million) in a cost-permissive manner. To our knowledge, digital PCR assays, which are regarded as the most sensitive [e.g. 65], compartmentalize on average 0, 1 or very few DNA molecules per compartment/microchamber. Accordingly, to detect one eDNA molecule of *P. jackschultzi*, > 275 million such compartments would have to be examined, which may be cost-prohibitive.

### 5.1 Inferences about changes in distribution of fish taxa and habitats compared to past (non-eDNA) surveys

Most of our eDNA-based inferences of effective presence or absence were not surprising, but the few exceptions (i.e., those deemed false negatives and false positives in Table S4) raise concerns about the use of this methodology to confidently determine presence/absence of taxa (discussed in detail below). Despite these drawbacks, our results should provide a baseline against which to compare future documentation of presence/absence or eDNA-based abundances in the study area, particularly in light of the increasing presence of non-natives, and the loss or degradation of native fish habitat in this region.

#### 5.1.1 Upper Cocospera (Cuitaca area; site 1; 2022; 1260 meters above sea level; masl)

To our knowledge, ours is the first documentation of fish taxa in the Cuitaca area. Above our sampling site, the stream was completely dry. This stretch is intermittent and highly dependent on the water table levels. Local inhabitants told us this stretch, “effectively dries at times when the nearby mine extracts high amounts of groundwater for its operations”. According to the Mexican government’s official records [66], this aquifer has six subterranean water exploitation concessions, of which 82% (=1.5 million m^3^/year) are for industrial purposes and assigned to a single mining company. We hypothesize that local fish populations along this headwater stretch regularly go extinct, and that our detection of only Longfin Dace (assuming that our very low eDNA detections of Sonora Chub and Gila Topminnow are false positives) reflects that Longfin Dace is a highly opportunistic early colonizer that can travel long distances in short time periods and persist under low water conditions [67, 68]. We predict that any of the species present at the mainstream of site 2 (Rancho Aribabi), including the native Sonora Chub, occasionally (re)colonize the Cuitaca stretch.

The inferred absence of eDNA from *Poeciliopsis* sp. at both Cuitaca (site 1) and Aribabi (site 2) contrasts with the 1986 record of *Poeciliopsis* sp. at a site approximately halfway [∼14 km upstream of Aribabi; i.e., site 17 in 32]. Because those specimens were fixed in formalin, hampering DNA-based identification, their species (or hybrid) identity has not yet been determined, but they likely included at least the Gila Topminnow. In addition, co-author D.K. Duncan observed the characteristic black nuptial males of *P. occidentalis* at Rancho Aribabi in 2006. However, a 2015 fish survey at Rancho Aribabi [69] did not record *Poeciliopsis*, but did record Mosquitofish. We infer that *Poeciliopsis* has gone extinct from at least the Aribabi to Cuitaca stretch, and possibly along the ∼5km stretch between Aribabi and the Presa (dam) Comaquito (a in Fig. 1). This 3,200,000 m^3^ capacity storage dam initiated operations in 1981 [70]. While the 1986 capture-based fish survey from this dam [32] only recorded four other teleosts, all non-natives (i.e., Green Sunfish, Bluegill, Black Bullhead, and Channel Catfish), Largemouth Bass was observed albeit not captured (A. Varela-Romero, pers. obs.). *P. occidentalis* and *P. monacha-occidentalis* have been consistently collected at the Imuris-Cocospera site (site 3; ∼15km downstream of Presa Comaquito) since the early 1980’s, and were unequivocally detected with our eDNA methods both years. Due to safety concerns, we did not visit the Cocospera stretch between Imuris and Presa Comaquito, but we predict that when water is present, *Poeciliopsis* sp. are likely able to disperse upstream from the Imuris area, possibly as far as immediately downstream of Presa Comaquito. The dam, however, must be an effective upstream dispersal barrier for all teleosts. Thus, re-establishment of *Poeciliopsis* above the dam would require assisted migration. Fortunately, our 2022–23 eDNA and capture surveys suggest that the non-native Mosquitofish, which was last recorded at Rancho Aribabi in 2015 [69], may have gone extinct in the Cocospera stretch above the dam. Three non-native teleosts (Green Sunfish, Bluegill, and Largemouth Bass), however, continue to persist above the dam; they collectively accounted for 100% of the teleost eDNA at the Aribabi Cienega, and 14 and 45% at Aribabi Mainstream (2022 and 2023, respectively). We predict that fish present at Aribabi Cienega and mainstream can disperse downstream to the dam, which has varied widely in volume during the last few years, with extremely low levels and unavailability for irrigation reported in July 2025 [71], but excessive levels requiring water releases reported in March 2020 [72].

#### 5.1.2 Lower Cocospera to lower Bambuto in the Imuris area (sites 3–5)

The sampled stretch spanning the lower Cocospera to the lower Bambuto around the town of Imuris (sites 3–5) had relatively similar teleost species composition based on our eDNA results. This stretch was the only one where eDNA from the non-natives Tilapia and Green Swordtail was effectively detected. Site 3 is the only Cocospera site where *P. jackschultzi* was ever recorded; with the last record in 2000, albeit at very low abundance (i.e., 1 out of 150) relative to the Gila Topminnow and Topminnow Asexual Hybrids [28]. The Imuris-Cocospera (2022 and 2023) and the Bambuto-Cocospera confluence (2022; elevation 821 masl) eDNA samples were the ones with the highest abundance of Gila Topminnow, and of Gila Topminnow and Topminnow Asexual Hybrid collectively, relative to other teleosts. We hypothesize that there is high connectivity for fishes along this stretch, but habitat/water availability are threatened. Firstly, despite the summer 2021 onset of operation of the updated wastewater treatment infrastructure for the town of Imuris (b in Fig. 1; [73]), at the time of sampling (May 2022), we detected what appeared to be raw sewage in the lowest stretch of the Cocospera, immediately upstream from where it meets the Bambuto (i.e., between site 3 and 4; see Dataset S14). Secondly, the ongoing construction (initiated in 2023) and future operation of the new train tracks between Imuris and Nogales, which essentially track the Cocospera from its confluence with the Bambuto (site 3) up to Rancho Aribabi (site 2), is expected to have numerous negative impacts for this tributary and further downstream, which may further compromise the viability of native fish species (and water availability for human use). Thirdly, this stretch harbors a high number and relative eDNA abundance of non-native teleosts. Among the most eDNA abundant non-natives in this stretch, Tilapia, Green Sunfish, and Black Bullhead have been known to be present in the region since at least 1986 [32]. In contrast, the presence of the Green Swordtail was not recorded in previous fish surveys of this stretch (the most recent in 2000) or of the larger study area. Thus, its introduction must have occurred within the last ∼20 years, possibly as a pet release from an Imuris area household. For the Mosquitofish, the earliest tentative record of presence in the entire Rio Concepcion appears to be from a site ∼20 km upstream (in the Middle Bambuto; near site c; Figure 1) during a 2009 capture-based survey, but identity of these specimens awaits to be confirmed (C.B. Dillman, pers. comm.; see Dataset S13). Capture-based surveys conducted by R.C. Vrijenhoek and/or coauthors Mateos and Varela-Romero, between the 1980’s and 2001 in Bambuto and Lower Cocospera area, which were mostly focused on *P. jackschultzi* [see 28], would likely have detected Mosquitofish or Green Swordtail had they been present.

#### 5.1.3 Middle Bambuto (also known as Alisos): sites 6–9

The ∼39 km “Middle Bambuto” stretch between sites 5 and the wastewater treatment plan (PTAR) Los Alisos (immediately upstream of site 9), contains the remaining historical range of *P. jackschultzi*, whose most upstream (and northernmost) record (from 1986) is likely around site 8 (La Cieneguita). Also in 1986, a now-disappeared spring-fed cienega adjacent to site 7 (La Atascosa), harbored the highest frequency (∼58%) of *P. jackschultzi* relative to other *Poeciliopsis* ever recorded. The last record of *P. jackschultzi* occurrence was in 2001 from the type locality; a short spring-fed stream that drained from the east (under the federal highway bridge currently known as “Puente La Atascosa”) to ∼0.5km downstream of our mainstream Bambuto eDNA site 7 [28]. Along this ∼39 km Middle Bambuto stretch, we sampled four sites (6–9) in 2023, and only one of these (i.e., a mainstream site at La Atascosa; site 7) in 2022. The relative eDNA abundance of non-natives (dominated by Green Sunfish) along the Middle Bambuto stretch was low (6–13%), except at site 6 (Agua Caliente); a pond (max depth ∼1m) bordered by trees and other vegetation, which is completely surrounded by highway lanes (maximum length x width of this cienega-like island ∼400 x 100m; see 10.6084/m9.figshare.30649319; Dataset S14). Site 6 is where one replicate detected a substantial eDNA abundance of mackerel, which we attributed to a field contaminant (see Results). Based on their high eDNA relative abundances in this pond, Green Sunfish (52%) and Black Bullhead (24%) likely outcompete the natives; only Asexual Hybrid Topminnow (13%) and Sonora Chub (1%) were effectively detected. Minnow trap surveys at the time of eDNA sampling only detected *Poeciliopsis*. A drain under the highway is apparent, which likely enables fish movement to and from the mainstream, at least at times of high precipitation. A 2020 fish survey of where this Agua Caliente tributary meets the mainstream (USON-1426; 30° 57.150’N, 110° 51.383’W) recorded Longfin Dace [64], Sonora Chub, and *Poeciliopsis* (Varela-Romero pers. obs.). Therefore, Longfin Dace are likely unable to successfully establish in this pond due to the intermittent nature of its connection to the mainstream and lack of current.

Of all the sites we visited along the Alisos-Bambuto, site 8 (La Cieneguita; elevation 1000 masl) appeared to retain the most cienega-like characteristics (see photos/videos at 10.6084/m9.figshare.30651914; Dataset S14). Therefore, we considered that this site had the highest likelihood of supporting *P. jackschultzi* (and the putative cienega-endemic chub *Gila* sp.). The eDNA sample from La Cieneguita was unique in that it had the highest relative abundance of Sonora Chub of all samples (76%), and an effective absence of Longfin Dace, which is not surprising given that thick-bodied chubs such as the Sonora Chub tend to be more abundant in ciénegas than elsewhere [74]. The 2nd-most eDNA-abundant (∼17%) fish at La Cieneguita was the Asexual Hybrid Topminnow, but its sexual host was only detected at 0.7%. While at low relative eDNA frequencies, the Green Sunfish, Black Bullhead and Mosquitofish were also detected. Seine and minnow trap surveys at the time of eDNA sampling recorded at least 10 *Poeciliopsis* sp. including 1 male, and 14 “minnows” (i.e., Longfin Dace or Sonora Chub).

The most upstream site sampled along the Middle Bambuto stretch (site 9 at El Cíbuta) was ∼1 km downstream of where the PTAR Los Alisos releases its treated effluent. Whereas visual surveys at site 9 revealed the presence of Longfin Dace, Sonora Chub, *Poeciliopsis* sp., and at least one Mosquitofish individual, eDNA effectively detected only Longfin Dace (51%), Sonora Chub (26%), and Green Sunfish (13%). Non-capture-based visual surveys performed by walking alongside the stream starting at the effluent discharge site revealed the presence of fish (likely Longfin Dace) beginning ∼40 m downstream, implying that the water quality was adequate for fish survival.

#### 5.1.4 La Mesa (Upper Bambuto; site 10)

Within the Bambuto, the highest elevation site where we found fish was at Arroyo La Mesa (site 10; elevation 1174 m asl), where Longfin Dace had the highest eDNA frequency (78%) followed by Sonora Chub (21.85%). Both native species were also recorded at this site in a 2009 capture-based survey (C.B. Dillman, pers. comm.; see Dataset S13), and were captured in a seine at the time of eDNA collection. Also captured in the seine were a substantial number of *Poeciliopsis* sp., including males of Gila Topminnow, as well as a smaller number of Mosquitofish, but the eDNA abundance of these poeciliids was extremely low (0.06 and 0.01%, respectively). According to our eDNA results, we did not detect Topminnow Asexual Hybrid above the threshold; and it was thus deemed effectively absent.

Approximately 1 km upstream of site 10, we found a short stream stretch containing water (site e; elevation 1190 m asl; Figure 1), but visual inspection revealed that it lacked fish. No water was observed above site e; the highest point surveyed was ∼2 km upstream of site e (i.e., dry site VIII; ∼1207 m asl; Figure 1). A small dam ∼2.6 km upstream of dry site VIII (site f; Fig. 1) exists since at least 1975 [75], but no records on water levels or flow exist for this tributary.

#### 5.1.5 La Arizona (Rio Altar; site 11)

The Rio Altar tributary of the Rio Concepcion (Fig. 1) is not considered within the known range of *P. jackschultzi*, but its headwaters come very close to headwaters of the Alisos-Bambuto, and few surveys documenting *Poeciliopsis* sp. in the Rio Altar exist [40]; Hendrickson, 1990 #1527’; and 1976 collections by R.C. Vrijenhoek}. We sampled this site because it was close to the Bambuto, accessible to us, and had water. This was a small stream immediately upstream of a small reservoir stocked with Largemouth Bass, of which eDNA was detected in our sample, and juveniles were captured in the seine. Our lack of detection at site 11 of eDNA from Tilapia, which was captured in the seine, highlights the limitations of our eDNA methodology. Similarly, our failure to detect eDNA from Topminnow Asexual Hybrid at site 11 could be a false negative, or may truly reflect its absence in the sample. Moore, Miller and Schultz [40] used dentition to distinguish Gila Topminnow from Topminnow Asexual Hybrid at four Rio Altar sites (sampled years: 1941, 1950, and 1955), and documented very low proportion (0–3%) of asexual females compared to Gila Topminnow females. R.C. Vrijenhoek (unpublished; pers. comm.) used allozymes to distinguish asexuals from Gila Topminnow from a 1976 survey of Rio Altar at Oquitoa (∼82 km downstream of site 11). Vrijenhoek documented very different relative frequencies of asexuals (i.e., 64.4 and 4.3%, respectively) from two samples; one taken at a side pool and one taken in the mainstream (n = 104 and 114, respectively). This, along with Moore, Miller and Schultz [40] findings from 36 other site/year samples from four other rivers, indicates that frequency of asexuals varies substantially over space and time. The most recent *Poeciliopsis* sample taken from the Rio Altar (1986 in [32]; fixed in formalin) did not distinguish between sexuals and asexuals.

#### 5.1.6 Santa Cruz (site 12)

Another goal of this study was to determine what teleost species are currently present in the Mexican stretch of the Rio Santa Cruz. Our eDNA results are the first to confirm the presence of Asexual Hybrid Topminnow in this stretch. The Rio Santa Cruz, and the entire Gila (and Colorado) River system have been considered outside of the Asexual Hybrid Topminnow’s native range. Moore, Miller and Schultz [40] state “Over 1000 specimens of *P. occidentalis* from scattered locations in the Gila River system have been examined with no evidence of the aberrant dentition characteristic of all-female forms”. These authors, however, did not specify if they had examined the only (to our knowledge) accessions of *Poeciliopsis* from the Mexican side of the Santa Cruz available at that time [i.e., Catalog Nos. 157241 (n = 11; 1940) and 162670 (n = 115; 1950) of the University of Michigan Museum of Zoology (UMMZ)]. The earliest documented presence of this asexual form in the Santa Cruz (and in the US) is from 2017 at Tubac and Tumacacori, which are ∼42 km downstream of our eDNA sampling site. The asexual is currently recorded as far downstream (and North) as Tucson (∼70 km from Tubac), where it likely arrived via inadvertent introduction along with Gila Topminnow from the Tubac-border area, as part of a conservation action in October 2020 [41].

To our knowledge, prior to this study, the few post-1950 records documenting *Poeciliopsis* in the Mexican stretch of the Santa Cruz (i.e., USON-0268 and USON-0274 from 1988, USON-01032 from 1996, a 2019 survey of the Madrean Discovery Expeditions [76] and a 2021 survey by co-author A. Varela-Romero), did not distinguish between Gila Topminnow and Asexual Hybrid Topminnow, and the specimens (if available) are preserved in formalin. Unless such specimens can be reliably diagnosed [e.g. with the dentition traits used by 40], it is impossible to know whether these samples/records contained asexuals. There are two possible hypotheses to explain the recent detection of the asexual topminnow in the Santa Cruz. One hypothesis is that the asexual is native to the Gila River, and persisted at least in certain sites (e.g. the Mexican side), but was not detected. Occasional water connections allow dispersal (at least downstream) into the US. The 2009 and 2013 upgrades to the Nogales International Wastewater Treatment (Fig. 1B) improved the quality and quantity of the Santa Cruz stretch downstream of the effluent discharge [77, 78]. The enhanced habitat may have enabled colonizers to persist, whereas prior colonizers would die out. It is worth noting that by the time Robert R. Miller (who initially discovered the asexual forms of *Poeciliopsis*) and collaborators were surveying the Gila River in Arizona, the fish habitats were highly degraded by human activities. It is thus possible that asexual topminnows were previously present in the US but extirpated. The Mexican side, which appeared to retain more viable habitat, may have served as a refugium. An alternative hypothesis is that the asexual was introduced by humans into the Santa Cruz (Mexican or US side) from its native range (e.g. nearby tributaries of the Rios Concepcion, Yaqui, or Sonora).

Our eDNA results from the Santa Cruz unequivocally detected the presence of *monacha* mitochondrial haplotype A, which is broadly distributed as far south as the Rio Mayo, and we have documented it in 17 specimens from Arizona (unpublished data). Whilst our Santa Cruz eDNA results also detected the presence of *monacha* mitochondrial haplotype D (considered endemic to the Concepcion), its frequency relative to haplotype A was < 6%, raising concern that it could be a false positive. Detection of haplotype D in at least one individual specimen from the Rio Santa Cruz, would dispel such concern. If confirmed, the presence of haplotype D in the Santa Cruz would be consistent with a human introduction from the Concepcion, or with a past natural connection between the Santa Cruz and Concepcion. Although a past connection that enabled the distribution of Longfin Dace across these drainages likely existed, the timing is unclear. Paredes Gallardo [64] detected genetic differentiation of Longfin Dace between the Santa Cruz and the Concepcion, but several haplotypes are shared between both drainages. Additional insight into the origin of the Santa Cruz asexual topminnow population may be gleaned from genetic variation in the clonal (i.e., *monacha*-derived) nuclear genome of the asexuals from throughout the distribution. Similarly, if asexuals were introduced to the Santa Cruz, such introductions likely included Gila Topminnow. Examination of variable nuclear markers of the Gila Topminnow (e.g. microsatellites or many SNPs) may reveal the source locality(ies) of topminnow putative introductions into the Santa Cruz.

The other relevant finding for the Santa Cruz was the detection of eDNA from what appear to be the Mexican Roundtail Chub (*Gila minacae*) and the Yaqui Sucker (*Catostomus bernardini*), neither of which is native to the Gila, Concepcion, or Sonora basins. The nearest localities known to harbor these two species are in the Río Bavispe branch of the Rio Yaqui [Varela-Romero, pers. obs.;79], including: (a) San Bernardino Creek (e.g. SBNWR = San Bernardino National Wildlife Refuge; in Fig. 1B), which is ∼150km straight-line distance from our Santa Cruz site; and (b) Arroyo Cajon Bonito (CB in Fig. 1B). The Bavispe branch also harbors/ed Yaqui Chub (*Gila purpurea*) and the endangered Yaqui Catfish *Ictalurus pricei* [80]. To our knowledge, there are no records of translocations from the Bavispe branch to the Río Santa Cruz. Nonetheless, in 1899, Yaqui Catfish was successfully introduced (presumably from the Rio Sonora) into a reservoir system fed by Monkey Spring (a headwater of Sonoita Creek; labeled “SC” in Fig. 1B), which meets the Santa Cruz mainstream ∼25km downstream from our sampled site. This introduced population of Yaqui Catfish persisted for half a century [67]. It is possible that this Yaqui Catfish introduction was from the Rio Yaqui and was accompanied by other Yaqui native fish, which dispersed to, and became established at, the Mexican side of the Santa Cruz. Further verification of the presence and identity of the putative Mexican Roundtail Chub and Yaqui Sucker whose eDNA we exclusively detected in the Santa Cruz is needed, including specimen-based surveys. We also recommend verification that the MiFish region is adequately diagnostic by examining additional specimens and localities representative of the entire range of the relevant species (e.g. *Gila purpurea*, *Gila minacae*, *Gila robusta* complex, *Catostomus insignis*, *Catostomus bernardini*).

### 5.2 Conservation Implications and Outlook

Several observations raise serious concerns about the conservation status of native fishes in the study area, most of which have some type of protected status in Mexico’s NOM-059, USA’s ESA, and/or the IUCN Red List (Table S1). Numerous sites that harbored native fish populations in the past do not appear to provide viable habitat at present (e.g. sites found dry during our 2022-2023 surveys; Fig. 1). The most concerning are the extensive loss of cienega habitats. Site II* was described in Vrijenhoek’s March 1984 fieldnotes as “A clear water cienega with many macrophytes. A good current ran through its center portion and across the road. The cienega ranges from running to swamp with rushes. It is bordered by willows and trees that look like aspens”. This site continued to harbor aquatic habitat and fishes (at least in the mainstream) during R.C. Vrijenhoek’s visits in November 1986, November 1989, September 1994, April 1999, and May 2000. However, photographs and fieldnotes from 1999 and 2000 reveal that the habitat at this site was no longer a cienega (see Dataset S14), and that the spring area and pool sampled in prior visits were completely dry. Finally, during our 2022 and 2023 visits, the entire area around site II* was completely dry.

Approximately 10 km north of site II*, at La Atascosa area (site 7; Fig. 1), a July 1980 photograph taken by Gary Meffe shows the “interior view of a mature cienega habitat” Figure 149 in Minckley *et al.* [81]. This Atascosa Cienega site was visited by R.C. Vrijenhoek in January 1981 and November 1986 (i.e., the collection with the highest frequency of *P. jackschultzi* relative to other *Poeciliopsis*). Subsequent surveys, beginning in 1989 (after the expansion of the federal highway from 2 to 4 lanes), failed to find cienega habitat at La Atascosa, but a small spring-fed tributary had water in January 2001 and yielded five specimens of *P. jackschultzi*, including the holotype [see 28; and Dataset S12]. This spring-fed tributary was completely dry during our 2022 and 2023 visits, implying that the aquifer does not reach the surface at this spring any longer. Given the ∼1.5 km distance between La Atascosa (site 7) and La Cieneguita (site 8), we hypothesize that they collectively encompassed a large contiguous mature cienega habitat, of which only a very small extent remains (at La Cieneguita). La Cieneguita appears to retain patches that remain permanently moist, and has springs that, during our 2022 and 2023 visits, were the source of the water for the mainstream site 7 that we sampled. This mainstream flow, however, had disappeared by site II*. Adjacent to site 8, there is a huge greenhouse operation presumably established in 2000, which has the highest groundwater extraction concession for agriculture of this aquifer (343,178 m^3^/year; [82]).

Most surface waters in the study area are considered intermittent. Spring-fed cienegas and small tributaries have likely provided the temporary refugia that enabled these native fishes to persist in the desert for millennia prior to the arrival of humans of European descent. The habitat connectivity afforded by periods of high precipitation likely enabled dispersal between refugia habitats. Dams, such as Presa Comaquito, restrict upstream dispersal, and are typically stocked with non-native species for human consumption (e.g. Tilapia and Black Bullhead) and sportfishing (e.g. Largemouth Bass), along with other prey/game species (e.g. Green Sunfish and Bluegill). Mosquitofish were likely introduced inadvertently with the above species or purposefully for mosquito control. Green Swordtail introduction likely reflects disposal of domestic pets. The detrimental effects of non-native species include competition, predation, hybridization, disease and parasite transmission, and alteration of the habitat [reviewed in 83]. Given the substantial evidence of Mosquitofish contributions to numerous local extinctions of Gila Topminnow in Arizona [reviewed by 11], we predict similar detrimental effects on topminnows in Sonora. For the larger predatory species (Bullhead and the three centrarchids), we expect them to have negative effects on topminnows via predation, whereas the poecilids Mosquitofish and Green Swordtail likely compete with topminnows and possibly consume their juvenile stages.

Due to increasing population size and economic activity, detrimental human impacts on the aquatic habitats of the study area are expected to continue increasing. Mexico’s National Commission on Water (CONAGUA) reports the three aquifers of the study area (2613 Río Alisos, 2614 Cocóspera, and 2615 Río Santa Cruz) as “with availability”. Nonetheless, we are concerned that such criteria are based on outdated data (e.g. last piezometric data from 2009, 2006, and 2008, respectively; [84]), and do not appear to take into account the effects of groundwater extraction on spring-fed habitats that are important for native fishes. We thus discourage the issuance of further groundwater extraction concessions. We are also concerned about the potentially detrimental effects of the new train route on water quantity and quality in the Cocospera. While the Mexican government has indicated that measures to mitigate the impacts of this project will be implemented, it is unclear what they are and whether they will suffice in the short and long term. We also discourage any further conversion of riparian and adjacent habitats.

Water reclamation could play an important role in the conservation and restoration of aquatic habitats in the study area. The recently upgraded wastewater treatment infrastructure in the town of Imuris [73] is an encouraging development, but is inadequate given our observation of what appeared to be untreated sewage in the Cocospera stretch immediately above its confluence with the Alisos-Bambuto. In addition, there are news reports of citizen complaints claiming raw sewage release and leaching from the Nogales municipal landfill into the upper Bambuto [85, 86]. The construction, operation and continued expansion of the PTAR Los Alisos (Fig. 1) is a positive development, because: (a) it reduces the net amount of exported water from the Río Alisos aquifer to the Río Santa Cruz basin via the Nogales International Wastewater Treatment Plant [87]; and (b) it increases the potential to replenish the Río Alisos Aquifer, albeit no evidence exists to date that these actions have led to the replenishment of the aquifer [88]. Up until recently, PTAR Los Alisos had a capacity to treat ∼220 liters/sec, but a recently added third module reportedly increased its capacity to ∼330 liters/sec [89–92], albeit actual processed volumes appear to be lower [93, 94]. We encourage the planned fourth module, which would increase capacity to ∼440 liters/sec [95, 96]. Observation of fish beginning ∼40m downstream of the effluent discharge at the time of our visit (2023) is encouraging, but concerns about the water quality of the effluent have been raised [88]. We recommend exploring the possibility of altering the habitat immediately downstream of the effluent discharge site for establishment of an artificial cienega-like habitat, that could slow down the flow and provide refugia habitat for native fishes. Overall, any infrastructure and practices that reduce groundwater extraction and increase the quality and quantity of treated wastewater into stream habitats of the study area will likely benefit native fish populations. Such efforts may benefit from public outreach to increase awareness among local stakeholders regarding the native fishes and their habitats. Ultimately, preservation of healthy aquatic habitats should benefit local human communities by contributing ecosystem services such as water for consumption and recreation opportunities in the river.

Regarding *P. jackschultzi*, we are extremely concerned about its status given that we failed to detect its presence. We recommend further monitoring of previously surveyed sites, but at different seasons (e.g. when there is more water), and in other unexplored tributaries, including: the south-to-north draining tributary of Presa Comaquito (a in Fig. 1), any mainstream and side tributaries between localities 6 and 7, the only stretch of the Alisos-Bambuto that continues to maintain large native riparian trees. Access to most of these sites requires passage through private properties for which we have been unable to obtain permission. In terms of techniques for detecting *P. jackschultzi*, despite its drawbacks, eDNA is the least invasive approach, particularly for a species whose females are not easily distinguishable from those of females of sympatric congeners. Should *P. jackschultzi* be found, we encourage ex situ conservation measures, at least while adequate habitat is identified and preserved or restored.

## 6. Conclusion

Our 2022 and 2023 surveys of the study area revealed substantial loss of habitat for freshwater fishes, particularly cienega-like habitats and adjacent streams. Our eDNA analyses failed to detect presence of the cienega-endemics Rio Concepcion Topminnow (*P. jackschultzi*) and an undescribed chub (*Gila sp.*), implying that they are extinct or present at abundances below our detection limits. Our collective evidence (i.e., visual/eDNA surveys and past records) confirm the presence of the other known native and previously reported introduced teleosts, and reveal more recent introductions (i.e., Green Swordtail and Mosquitofish) to the Rio Concepcion. We detect the apparent extirpation of Gila Topminnow from sites above the Comaquito dam in the Cocospera tributary (Rio Concepcion). In the Rio Santa Cruz (tributary of the Gila River), we document the first record of presence of the Asexual Hybrid Topminnow in the Mexican side, as well as of two putative non-native fish species (i.e., Yaqui Sucker and Mexican Roundtail Chub). Whilst for the most part, our eDNA results agreed with concurrent and previous capture-based taxon records, a few eDNA false negative results raise concerns about the detection limits of this approach. Similarly, a small number of putative eDNA false positive results, underscore the need for appropriate controls and mitigation of contamination. We recommend further surveillance of habitats and fish taxa, and implementation of practices to enhance groundwater recharge and the quality and quantity of treated wastewater in the study area.

## Author contributions (alphabetical name order)

Conceptualization: AVR,MM

Investigation: AF,AGB,AVR,DKD,MM

Data Curation: AF,MM

Visualization: AF,MM

Resources: AF,AVR,DKD,MM

Funding Acquisition: AF,AVR,MM

Supervision and Project Administration: AVR,MM

Writing – original draft: AF,MM

Writing – review and editing: AF,AGB,AVR,DKD,MM

## Acknowledgments

Logistical support: Rancho El Aribabi (Guzman); Sky Island Alliance (Monica Montaño, Angel García, and Miguel Enriquez); and Luis Fernando Duarte (DICTUS, Universidad de Sonora). Resources: US Bureau of Reclamation (Kent Mosher), Universidad de Sonora.

Technical advice: Yale Passamanek.

The following individuals contributed DNA from specimens to generate MiFish reference sequences: Jessi Rick (University of Arizona) for *Catostomus*; Tyler Chafin, Marlis Douglas, and Michael Douglas (University of Arkansas) for *Gila purpurea*; and Carlos Alonso Ballesteros Córdova (Universidad de Sonora) for *Gila* from Mexico.

Topiltzin Contreras-MacBeath (Universidad Autónoma del Estado de Morelos) for assistance with funding acquisition.

Portions of this research were conducted with the advanced computing resources provided by Texas A&M High Performance Research Computing and by the European Galaxy server usegalaxy.eu.

Robert C. Vrijenhoek and Casey B. Dillman shared unpublished collection records.

Peter Unmack and Bill Fagan shared the database SONFISHES database

## Ethical Statement

All procedures were approved by the Texas A&M University IACUC (Protocol #2023-0106) Field work was authorized under Licencia de colecta-SPARN/DGVS-02635-2022 issued to Alejandro Varela-Romero by the government of Mexico

## Use of Artificial Intelligence (AI) and AI-assisted technologies

We have not used AI-assisted technologies in creating this article.

## Data Accessibility

All supporting data, code and other research materials can be accessed as follows:

- All novel sequencing data are available through National Center for Biotechnology Information (NCBI) repositories. Illumina raw reads available under BioProject nos. PRJNA1406547 and PRJNA1367221. Assembled sequences based on Sanger sequences are available under PX556618-PX556621, PX637987-PX637989, PX746530-PX746546, and PZ418347-PZ418350. The assembled and annotated mitochondrial genome sequence of *Poeciliopsis jackschultzi* is under PX582708.
- All other files are available as electronic supplementary material to this submission and will be deposited at the public repository required or recommended by the accepting peer-reviewed journal.

## Supporting Figure Descriptions

**Figure S1**

**The origin and maintenance of hybridogenesis**

The origin and maintenance of hybridogenesis (= hemiclonality), exemplified by the unisexual biotype *Poeciliopsis monacha-occidentalis* (MO). Each capital letter reflects a haploid genome set from *P. monacha* (M) and *P. occidentalis* (O). This biotype originated through a cross of two sexual diploid species: a female *P. monacha* (MM; white) and a male P*. occidentalis* (OO; blue). The F1 hybrid of such a cross is a diploid female (M*O; light blue to indicate its intermediate phenotype), that produces haploid M* eggs via an abnormal meiosis that excludes the paternally derived (O) chromosomes. The asterisk denotes an identical set of *P. monacha*-derived chromosomes (barring any new mutations). The dashed arrows connect the genetically identical (barring new mutations) haploid eggs (M*) produced by subsequent generations. The hemiclonal lineage persists through fertilization by males from the sexual host species P. occidentalis. Superscripts indicate different genetic composition. The black dot at the base of the dorsal fin is a diagnostic trait (dominant) of *P. occidentalis* that is present in the MO hybrids [97]. Although not depicted, the mitochondrial genome of the MO biotype is also clonally transmitted and originally derived from *P. monacha*.\

**Figure S2**

**Absolute eDNA frequencies**

Absolute frequencies of 21 teleost taxa in the 73 samples subjected to eDNA metabarcoding (N = native; I = introduced; S = spurious;= likely introduced from a nearby drainage; see text for details). Spurious taxa whose collective frequency among all samples was >6,000 are retained. Samples are grouped by year/site. The “Pass” label refers to groups whose replicates exhibit similar relative frequencies of non-spurious taxa and whose blank (where applicable) had very low frequencies of non-spurious taxa. Based on these criteria, the Aribabi Cienega MV22-1 group is considered failed, and thus excluded from downstream analyses. Site number match those in Fig. 1. Data used and original graphs found in sheet “counts_by_final_taxon2_ro” of Dataset S3.

**Figure S3**

**Neighbor-Joining (NJ) tree of eDNA features assigned to Leuciscidae and reference sequences**

Neighbor-joining (NJ) tree (p-distance; branch lengths = parsimony steps) of the retained Leuciscidae (formerly Cyprinidae) eDNA features and reference sequences. Red tip labels = most abundant features per species (or features referred to in text). Blue tip labels = references that perfectly or closely matched eDNA features. Magenta Tip Labels = reference sequences that we consider belong to the *Gila robusta* complex. Green Tip Labels = newly generated *Gila* spp. reference sequences that are closely related to the Yaqui Chub *Gila minacae*. Features tip labels include the 32 alphanumeric labels assigned by QIIME2, followed by the taxonomic assignment, the collective frequency and the number of samples in which it was found (if applicable). Reference sequence labels include voucher ID and/or NCBI accession number. Grey highlight = clade of Longfin Dace *Agosia chrysogaster* references and eDNA features. Orange branches = clade of Sonora Chub *Gila ditaenia* references and eDNA features. Yellow highlight = clade of *Gila minacae* references and the chub eDNA feature that was exclusive to the Rio Santa Cruz (site 12). Data available in Dataset S4.

**Figure S4**

**Frequency histograms of eDNA features assigned to each of twelve teleost species.** Frequency histograms of features within each of 12 teleost species, showing that the majority of features assigned to a taxon have very low abundance; > 95% of the cumulative abundance per species is contributed by ∼1–3 features. Data used and original graphs found in sheet “features_passed_filters1&2” of Dataset S3.

**Figure S5**

**Neighbor-Joining (NJ) tree of eDNA features assigned to Poeciliidae and reference sequences**

Neighbor-joining (NJ) tree (p-distance; branch lengths = parsimony steps) of the retained Poeciliidae eDNA features and reference sequences. Red tip labels = most abundant features per species (or features referred to in text). Blue tip labels = references that perfectly matched eDNA features (and reference for *Poeciliopsis jackschultzi*). Green tip labels = Other relevant reference sequences. Features tip labels include the 32 alphanumeric labels assigned by QIIME2, followed by the taxonomic assignment, the collective frequency and the number of samples in which it was found. Reference sequence labels include voucher ID and/or NCBI accession number. Additional information such as haplotype ID or locality information is given for a subset of features. Tree is rooted with *Heterandria formosa*. Data available in Dataset S5.

**Figure S6\**

**Neighbor-Joining (NJ) tree of eDNA features assigned to Catostomini and reference sequences**

Neighbor-joining (NJ) tree (p-distance; branch lengths = parsimony steps) of the Catostomini (Sucker) eDNA features (red tip labels) recovered from site 12 (Parque Mascareñas, Rio Santa Cruz, Gila River, Mexico) and reference sequences. Green tip labels = samples of representatives of the two native suckers from the Gila River: Sonora Sucker *Catostomus insignis* from the San Pedro; and Desert Sucker *Pantosteus clarkii* from the San Pedro and Santa Cruz. Blue tip label = Yaqui Sucker *Catostomus bernardini* sample from the Rio Fuerte; which perfectly matched our most abundant sucker eDNA feature(6e7ec8e5e4e6637ea39b5e01a542e824). Tree is arbitrarily rooted. Data available in Dataset S6.

**Figure S7**

**Relative frequencies of most abundant Topminnow Asexual Hybrid features**

Relative frequencies of the two most abundant eDNA sequences assigned to Topminnow Asexual Hybrid which perfectly matched the reference (i.e., *Poeciliopsis monacha*) haplotypes A and D for each sample (excluding blanks and Aribabi Cienega 2022). Numbers within or adjacent to bars are absolute abundances. Data used and original graphs found in Dataset S8.

**Figure S8**

**Relative frequencies of most abundant Gila Topminnow (*Poeciliopsis occidentali*s) features**

Relative frequencies of the three most abundant eDNA sequences assigned to Gila Topminnow for each sample (excluding blanks and Aribabi Cienega 2022). Numbers within or adjacent to bars are absolute abundances. Data used and original graphs found in Dataset S9.

**Figure S9**

**Relative frequencies of most abundant Longfin Dace (*Agosia chrysogaster*) features** Relative frequencies of the five most abundant eDNA sequences assigned to Longfin Dace in each sample (including blanks and Aribabi Cienega 2022). The X-axis only shows the 90–100% range. Numbers within or adjacent to bars are absolute abundances (where applicable). Data used and original graphs found in Dataset S7.

**Figure S10**

**Relative frequencies of most abundant Sonora Chub *(Gila ditaenia*) features**

Relative frequencies of the four most abundant eDNA sequences assigned to Sonora Chub in each sample (including blanks and Aribabi Cienega 2022). The X-axis only shows the 90–100% range. Numbers within or adjacent to bars are absolute abundances (where applicable). Data used and original graphs found in Dataset S7.

**Figure S11**

Relative eDNA frequencies of the thirteen non-spurious teleost taxa per each site/year sample excluding blanks and Aribabi Cienega 2022. N = native; I = introduced; S = spurious;= likely introduced from a nearby drainage (see text for details). Original graph generated in sheet “cnts_by_final_taxon2_ro_no_blnk” in Dataset S3, and edited with Inkscape.

**Figure S12**

**Relative frequencies of most abundant Mosquitofish *(Gambusia*) features**

Relative frequencies of the three most abundant eDNA sequences assigned to Mosquitofish for each sample. Numbers within or adjacent to bars are absolute abundances. Data used and original graphs found in Dataset S10.

## Supporting Table Descriptions

**Table S1**

**Status of native fish taxa in the study area**

Status of native fish taxa in Study Area (Gila and Concepcion), as well as several fish taxa native to the neighboring Ríos Yaqui and Sonora, which are relevant to this study. Whether or not at least one MiFish region sequence is available for comparison is indicated. P = endangered (“en peligro de extinción”); A = threatened (“amenazada”); Pr = subject to special protection (“sujetas a protección especial”); n/a = not applicable because species has not been formally described and/or because species is not native to corresponding country.

**Table S2**

**Detailed sample information**

Compressed folder: containing a tab-delimited text file with detailed sample information and a Metadata file describing contents. An Excel® Workbook (.xlsx) version of this sheet and associated sheets is provided in Dataset S1.

**Table S3**

**Sanger-sequenced specimens**

Specimens for which the MiFish region was PCR-amplified and sequenced with Sanger technology, including voucher ID, GenBank Accession number, and collection information.

**Table S4\**

**Collective evidence used to infer presence/absence of each of 13 teleost taxa**

Collective evidence used to infer presence/absence of each of 13 teleost taxa considered non-spurious. For each site/year, values = relative frequency of the sum of replicates (replicate values above “all blanks” threshold). Spurious taxa excluded from counts; and Aribabi Cienega 2022 excluded. Obs (Y/N) = whether taxon was observed during eDNA sampling. Exp (Y/N) = whether taxon is expected based on prior/nearby records. Inference = whether the taxon is inferred as present (P) or absent (A) based on the collective evidence. Red cells = deemed false positive eDNA results; Green cells = deemed false negative eDNA results. Alternative formats and metadata for this table are provided in Dataset S16.

## Supporting Dataset Descriptions

**Dataset S1**

**Alternative format for Table S2**

Compressed folder containing Table S2 Detailed Sample Information in Excel® (.xlsx) format and a metadata file.

**Dataset S2**

**Features output by Qiime2 pipeline**

Contains the 2924 eDNA features (Amplified Sequence Variants; ASV) generated by Qiime2 pipeline in fasta format

**Dataset S3**

**Feature frequencies and taxon assignment**

Compressed folder containing the Excel® Workbook used to apply Bioinformatics filters (see text) and compute the eDNA relative and absolute frequencies (= abundance) and prepare Figures 2, S2, S5, S11. Tab-delimited txt versions of each sheet (values and formulas) are also provided along with a Metadata file.

**Dataset S4**

**Leuscicidae phylogeny files**

Compressed folder containing alignments and commands used to generate Fig. S3. See included Metadata file for details.\

**Dataset S5**

**Poeciliidae phylogeny files**

Compressed folder containing alignments and commands used to generate Fig. S5. See included Metadata file for details.

**Dataset S6**

**Catostomini phylogeny files**

Compressed folder containing alignments and commands used to generate Fig. S6. See included Metadata file for details.

**Dataset S7**

**Data for Relative frequencies of most abundant Longfin Dace and Sonora Chub features** Excel® Workbook used to generate Figs. S9 and S10. Contains the eDNA abundance per sample of the most frequent features assigned to Longfin Dace (*Agosia chrysogaster*) and Sonora Chub (*Gila ditaenia*).

**Dataset S8**

**Data for Relative frequencies of most abundant Topminnow Asexual Hybrid features**

Excel® Workbook used to generate Fig. S7. Contains the eDNA abundance per sample of the most frequent features assigned to the mitochondrion of *Poeciliopsis monacha*, and thus assigned to Topminnow Asexual Hybrid.

**Dataset S9**

**Data for Relative frequencies of most abundant Gila Topminnow features**

Excel® Workbook used to generate Fig. S8. Contains the eDNA abundance per sample of the most frequent features assigned to Gila Topminnow *Poeciliopsis occidentalis* ().

**Dataset S10**

**Data for Relative frequencies of most abundant Gila Topminnow features**

Excel® Workbook used to generate Fig. S12. Contains the eDNA abundance per sample of the most frequent features assigned Mosquitofish (*Gambusia*).

**Dataset S11**

**Localities not sampled in 2022 or 2023**

Details on localities in the study area that were not sampled, including sites that were dry, most of which are indicated in Fig. 1.

**Dataset S12**

**Vrijenhoek Field Records**

Excerpts of records from field surveys in the Rio De La Concepción performed by R.C. Vrijenhoek and collaborators between 1981 and 2001.

**Dataset S13**

**Casey B. Dillman’s Collection Records**

Excerpts from records of collections made by Casey B. Dillman, Richard L. Mayden and John Hatch in April 2009 at two sites in the Rio De La Concepción.\

**Dataset S14\**

**Supporting Photographs/Videos Descriptions**

Descriptions and weblinks (figshare repository) of photographs and videos of the study area.\

**Dataset S15**

**Findable, Accessible, Interoperable, Reusable eDNA (FAIRe) metadata**

Completed FAIRe checklist following [44] in Excel® (.xlsx) format.

**Dataset S16**

**Alternative formats for data in Table S4**

Compressed folder containing alternative file format for Table S4 Collective evidence used to infer presence/absence of each of 13 teleost taxa (see included metadata file).\

**Dataset S17**

**Supporting Sanger Sequencing Chromatograms**

Compressed folder containing the original sequence chromatograms (.ab1 format) generated in this study. Each subfolder is labeled with the corresponding specimen voucher ID, which can be matched to specimens and GenBank IDs in Table S3.

## Supporting Protocols Descriptions

**Supporting Protocol S1**

**Description of Wet-lab protocols and conditions.**

A. Description of Clean Lab for pre-PCR processing of samples.
B. Protocols for PCR Reaction first round.
C. Protocols for PCR Reaction second round

**Supporting Protocol S2**

**Description of Bioinformatics pipelines.**

Compressed folder with several text files describing the steps command lines to process data in this study, including:

-cutadapt to demultiplex and trim Illumina reads
-Qiime2 to infer eDNA features and their frequences
-inference and visualization of distance-based trees of eDNA features and reference taxa
-read mapping protocol to determine that no reads from *Poeciliopsis jackschultzi* were present in the eDNA reads.

