## Supplementary material for "Environmental DNA analysis to search for an endangered desert fish": Medatata_Dataset_S1_19May2026_1.txt

Metadata for Dataset S1. Detailed sample information in Excel format

This Workbook contains the following sheets 
"main_convert_to_text" has the same content as Table S2, but some of its fields were populated via formulas dependent on the other sheets
	"collect_date_geog_coord" 
	"manifest_73plus"
	"IDs_from_TableS1"
	"index_names"
	"read_stats"


Metadata for Table S2 and sheet "main_convert_to_text"

Each row represents a field sample (replicate or blank) or a lab negative control
Explanation of column headings (if applicable):
	Site Number (in Map): the number assigned to sample in the Map (1-12)
	Field Collection Site ID: label to identify the site/year sample
	Filter ID: label to identify the filter (blank, 1, 2, or 3) at each site/year sample
	Alternative Collection ID: alternative label to identify the site/year sample
	Alternative Filter ID: alternative label to identify the filter (blank, 1, 2, or 3) at each site/year sample
	Field Collection Site and filter ID: concatenation of Field Collection Site ID and Filter ID
	Sample number in QIIME pipeline
	Locality name and sample ID in Tables and Figures
	Locality Full Name (as in Fieldnotes)
	Latitude
	Longitude
	Volume filtered (L):seconds of air (for blanks) or volume of water filtered for each filter
	Collection Date: date eDNA sample was collected in the field
	Extraction Date: date DNA was isolated from filter
	Library Prep batch: Batch of samples processed (1 or 2)
	Positive Control in PCRs: Species whose DNA was used as positive control for PCRs (Gambusia or Heterandria)
	The following "Qiime..." columns are direct output of the Qiime2 pipeline
		Qiime input: number of reads used as input for Qiime (extracted from file “denoising-stat_73_samples.qzv”)
		Qiime filtered: number of reads that passed Qiime’s filter (extracted from file “denoising-stat_73_samples.qzv”)
		Qiime percentage of input passed filter: percentage of reads that passed Qiime’s filter (extracted from file “denoising-stat_73_samples.qzv”)
		Qiime denoised: number of reads retained by Qiime after denoising step (extracted from file “denoising-stat_73_samples.qzv”)
		Qiime merged: number of denoised reads that were merged by Qiime (extracted from file “denoising-stat_73_samples.qzv”)
		Qiime percentage of input merged: percentage of denoised reads that were merged by Qiime(extracted from file “denoising-stat_73_samples.qzv”)
		Qiime non-chimeric: number of merged reads that were found to be non-chimeric and thus retained by Qiime (extracted from file “denoising-stat_73_samples.qzv”)
		Qiime percentage of input non-chimeric: percentage of merged reads that were found to be non-chimeric and thus retained by (extracted from file “denoising-stat_73_samples.qzv”)
	Index name: label used to identify the combination of unique Index F and Index R
	Index F: unique 10bp barcode sequence added to the 5’ end of the MiFish-U-F_primer of the second-round PCR. Within each library batch, each sample had a different Index F.
	Index R: unique 10bp barcode sequence added to the 5’ end of the MiFish-U-R_primer of the second-round PCR. Within each library batch, each sample had a different Index R.
	MiFish-U-F_primer (First-round Forward Primer): Sequence (5'-3' direction) of first-round PCR Forward Primer (Miya et al. 2015; doi: 10.1098/rsos.150088); also used for amplification and Sanger sequencing of reference specimens.
	MiFish-U-R_primer (First-round Reverse Primer): Sequence (5'-3' direction) of first-round PCR Reverse Primer (Miya et al. 2015; doi: 10.1098/rsos.150088); also used for amplification and Sanger sequencing of reference specimens.
	Second-round forward primer: Sequence (5'-3' direction) of Forward primer used in second-round PCR. It is a concatenation of Index F and MiFish-U-F_primer. These concatenated sequences can be used by cutadapt to demultiplex samples and trim primers prior to submitting to Qiime2 pipeline.
	Second-round reverse primer: Sequence (5'-3' direction) of Reverse primer used in second-round PCR. It is a concatenation of Index R and MiFish-U-R_primer. These concatenated sequences can be used by cutadapt to demultiplex samples and trim primers prior to submitting to Qiime2 pipeline.

Rows 76–84 represent sums/means/standard deviations of sets of rows for a subset of columns (see corresponding description in Column X)
