## Supplementary material for "Environmental DNA analysis to search for an endangered desert fish": DatasetS3_metadata.txt

Metadata for "Table Feature Frequencies & taxon assignment"

Contains the following nine worksheets, several of which are connected to each other through formulas:
working copy all
counts_by_final_taxon2
for_final_taxonomic_assign_MM
working copy -filtering steps
counts_by_final_taxon
counts_by_final_taxon2_ro
cnts_by_final_taxon2_ro_no_blnk
features_passed_filters1&2
features_passed_filters1&2_cnts


Metadata for sheet "working copy all": Contains all the features (2925 total) and their frequencies by sample
	feature: feature ID assigned by QIIME
	Top blast hit taxonomic ID
	Teleost (Yes/No)
	Passed filter for large indels/putative chimera/PCR artifact: No = did not pass filter and thus was discarded. Other values = passed filter
	Passed filter for putative lab contaminant: No = did not pass filter. Other values = passed filter
	Passed filter for field contaminant: No = did not pass filter. Other values = passed filter
	Passed filter for very low frequency/present at few sites: this column is meaningless; ignore
	Final taxonomic assignment scientific name: this column is meaningless/reduntant; ignore
	Final taxonomic assignment common name: this column is meaningless/reduntant; ignore
	Mateos taxonomic assignment: refined taxonomic assignment. Sonora Sucker should be Yaqui Sucker. Gila (not ditaenia) should be Gila minacae.
	Y if passed filters 1&2: Y for samples that passed filters 1&2; otherwise = No
	Mateos taxonomic assignment for those that passed filters 1&2: refined taxonomic assignment for features that passed filters 1&2
	sub-taxonomic assignment 1: this column is meaningless; ignore
	sub-taxonomic assignment 2: this column is meaningless; ignore
	Assigned haplotype?: this column is meaningless; ignore
	Teleost family: Cyprinidae should be Leuciscidae
	Frequency: sum of frequencies of feature in all samples
	The remaining columns correspond to each of the 73 samples"
		row 1 field collection ID
		row 2: locality name plus ID
		row 3: sample number (01-73, as labeled in Qiime)
		all other rows: frequency of each feature per each sample
	

Metadata for sheet "counts_by_final_taxon" for each of 71 preliminary teleost taxa, the collective abundance per sample is shown. The comments column provides rationale for considering discarding (i.e. 50 preliminary taxa). Catosotomus insignis should be Catostomus bernardini. Gila (not ditaenia) should be Gila minacae. 

Metadata for sheet "counts_by_final_taxon2": same as for "counts_by_final_taxon" but limited to the 21 teleost retained teleost taxa.

Metadata for sheet "counts_by_final_taxon2_ro" result of applying Bioinformatics Filter #3: For each of the 13 teleost species considered "non-spurious", we identified the negative control with the highest abundance (hereafter “all blanks threshold”), which ranged from 0 to 1,293. We then converted to zero the abundance of the corresponding species for every sample that did not have an abundance above the “all blanks threshold. Columns CG-FA are the result of Bioinformatics Filter #3. Bar graphs of absolute or relative frequencies of each teleost taxon per sample are provided.

Metadata for sheet "cnts_by_final_taxon2_ro_no_blnk": similar to "counts_by_final_taxon2_ro", with the addition of summation of replicates within year/site, and corresponding bar graphs.

Metadata for sheet "for_final_taxonomic_assign_MM": this sheet is redundant; ignore.

Metadata for sheet "working copy -filtering steps": this sheet is redundant; ignore.

Metadata for sheet "features_passed_filters1&2": List of features that passed filters 1&2, their taxon assignment and collective abundance. These data were used to plot the frequency histograms of each feature for each of the 13 teleost taxa deemed "non-spurious" 
 
Metadata for sheet "features_passed_filters1&2_cnts": Data and graphs showing the relationship between collective abundance and number of features per each of the 13 teleost taxo deemed non spurious.
