## Supplementary material for "Environmental DNA analysis to search for an endangered desert fish": Metadata_Dataset_S4.txt

Metadata for Supporting Dataset S4

Names and descriptions of files used to generate Supporting Figure S3

1. Leuciscidae_805_02Apr2026.nex:   Nexus-formatted file containing the aligned sequences of references and eDNA features assigned to Leuciscidae, prior to trimming with Genenious positions that have at least 75% of taxa with gaps.


2. Leuciscidae_805_02Apr2026_75_gapsstripped.nex: Nexus-formatted file containing the aligned sequences of references and eDNA features assigned to Leuciscidae after gapped positions (i.e. those in which at leat 75% of taxa had gaps) removed.

3. Leuciscidae_805_02Apr2026_75_gapsstripped_p_nj_pars_brlens.tree: Text file containing the a Neighbor-joining tree based on Jukes-Cantor distances. Branch lengths represent parsimony steps

	Commands used in paup to generate the above tree:
		execute Leuciscidae_805_02Apr2026_75_gapsstripped.nex
		set criterion = distance
		dset dist=p missDist=ignore
		nj
		set criterion = parsimony
		savetrees file = Leuciscidae_805_02Apr2026_75_gapsstripped_p_nj_pars_brlens.tree brlens=yes

4. Leuciscidae_805_02Apr2026_75_gapsstripped_p_nj_pars_brlens_colors.tree: tree file processed in FigTree v1.4.4, where TipLabels were replaced according to "TipLabel_clean_1_02Apr2026.txt" file, and colored. This tree was exported as an image to generate Supporting Figure S5

5. TipLabel_clean_1_02Apr2026.txt
