## Supplementary material for "Environmental DNA analysis to search for an endangered desert fish": Metadata_Dataset_S5.txt

Metadata for Supporting Dataset S5

Names and descriptions of files used to generate Supporting Figure S5

1. Poeciliidae_features_plus_references09Mar2024_737taxa178chars_trimmed.nex: Nexus-formatted file containing the aligned sequences of references and eDNA features assigned to Poeciliidae

2. Poeciliidae_features_plus_references09Mar2024_737taxa_NJ_JC_brlens_pars.tree: Text file containing the a Neighbor-joining tree based on Jukes-Cantor distances. Branch lengths represent parsimony steps

	Commands used in paup to generate the above tree:
		execute Poeciliidae_features_plus_references09Mar2024_737taxa178chars_trimmed.nex
		set criterion = distance
		dset dist=p missDist=ignore
		nj
		set criterion = parsimony
		savetrees file = Poeciliidae_features_plus_references09Mar2024_737taxa_NJ_JC_brlens_pars.tree brlens=yes

3. Poeciliidae_features_plus_references14Nov2025_737taxa_NJ_JC_brlens_pars_colors.tree: tree file processed in FigTree v1.4.4, where TipLabels were replaced according to "TipLabels..." file, and colored. This tree was exported as an image to generate Supporting Figure S5

4. TipLabels14Nov2025.txt
