## Supplementary material for "Environmental DNA analysis to search for an endangered desert fish": Metadata_Dataset_S6.txt

Metadata for Supporting Dataset S6

Names and descriptions of files used to generate Supporting Figure S6

1. SuckerMiFish_125taxa_170bp.nex: Nexus-formatted file containing the aligned sequences of references and eDNA features assigned to Catostominae

2. SuckerMiFish_125taxa_170bp_p_dist_NJ_parsbrlens.tree: Text file containing the a Neighbor-joining tree based on Jukes-Cantor distances. Branch lengths represent parsimony steps

	Commands used in paup to generate the above tree:
		execute SuckerMiFish_125taxa_170bp.nex
		set criterion = distance
		dset dist=p missDist=ignore
		nj
		set criterion = parsimony
		savetrees file = SuckerMiFish_125taxa_170bp_p_dist_NJ_parsbrlens.tree brlens=yes

3. SuckerMiFish_125taxa_170bp_p_dist_NJ_parsbrlens_colors13Nov2025.tree: tree file processed in FigTree v1.4.4, where TipLabels were replaced according to "SuckerTipLabels13Nov2025_2.txt" file, and colored. This tree was exported as an image to generate Supporting Figure S5

4. SuckerTipLabels13Nov2025_2.txt
