## Supplementary material for "Environmental DNA analysis to search for an endangered desert fish": Dataset_S11.txt

Dataset S11 
Localities not sampled in 2022 or 2023 (indicated in Fig. 1)

Site I. Dry in 2022 and 2023. Alisos-Bambuto mainstream 30.912103, -110.863418.

Site II*. Dry in 2023. Alisos-Bambuto mainstream and former adjacent cienega “La Providencia”. 30.919221, -110.860985. This locality has been referred to as La Providencia, Rancho Las Playas, and Rancho Los Alisos. See Vrijenhoek Field Records (Dataset S12).

Site III. Dry 2022 & 2023. Alisos-Bambuto mainstream Real de Catorce subdivision: 31° 0.805'N, 110° 53.561'W dry (2022 and 2023; “Real de Catorce”). This is ~6 km downstream from Bambuto mainstream at Cibuta #1.

Site IV. Dry in 2023. Alisos-Bambuto  mainstream at Cibuta #1  (31° 3.784'N, 110° 54.188'W). ~1 km downstream of Bambuto mainstream at Rancho Arochi

Site V. Dry in 2023. Alisos-Bambuto  mainstream at Rancho Arochi (31° 4.321'N, 110° 54.343'W). ~2.5km downstream of Bambuto mainstream at Alisos Pozo (well) #5.

Site VI*. Dry in 2023. Alisos-Bambuto mainstream Alisos Pozo (well) #5. 31.09144°N,110.91244°W. This is ~ 5 km downstream of our Cibuta sampling site (MV23-07). This site coincides with the location of the dead cienega (“Ciénega Rio Magdalena”) documented in Cole, A. T., and Cole, C. 2015. An overview of aridland ciénagas, with proposals for their classification, restoration and preservation. The New Mexico Botanist.  (hdl.handle.net/2152/30285), 28–56. 

Site VII. Dry 2023. Arroyo La Casona (= Subcuenca Nogales; 31° 10.995'N,110° 58.453'W); just west from where Hwy 15 meets Hwy 43 to Saric. This site is ~1km upstream of where it meets with Arroyo La Mesa, which is this upper stretch of the Alisos-Bambuto  is called in GoogleEarth

Site VIII. Dry in 2023. Arroyo La Mesa: 31° 11.933'N, 111° 0.697'W. This is about 2.5km from a storage dam called “El Muro” according to the 1975 topographic map. 

Site e. 2023. 31.191096, -110.994771. About 2–2.4km downstream of site VIII, we saw water in the stream, but it didn’t flow downstream and had no fish. It just appeared to be a place where the underground stream was briefly exposed. ~1.5 km downstream is our sampled site MV23-01. 

Not shown. Dry in 2023. 31.172758897074708, -111.19468604141781 confluence with the stream that comes down from La Esmeralda was completely dry. This site is ~5.5km downstream from our sample site MV23-06  Ejido La Arizona.
