## Supplementary material for "Environmental DNA analysis to search for an endangered desert fish": Dataset_S12.txt

Dataset S12 
Vrijenhoek Field Records

Excerpts of records from field surveys in the Rio De La Concepción performed by R.C. Vrijenhoek and collaborators between 1981 and 2001

Field No.: VD81-1
Collectors: Robert C. Vrijenhoek and Michael Douglas
Date: 1 August 1981
Time: 3:30PM
Description: Rio Magdalena-cienega on W side of H 15 about 24 miles S of Nogales
Water: clear
Vegetation: marsh
Bottom: mud; about 1-2 ft

Field No.: VS83-1
Collectors: Robert C. Vrijenhoek and R.A. Schenck
Date: 24 March 1983
Time: 1PM
Description: La Atacosa Cienega, 24 miles S of Nogales on Hwy 15. Collected swamp off right side of road at La Alacosa Ranch
Water: clear above swamp			
Vegetation: much vegetation; like Potamogeton
Air: ~24°C
Bottom: mud; this is a bog
Current: none

Field No.: VM84-9
Collectors: Robert C. Vrijenhoek and Gary Meffe
Date: 27 March 1984
Time: 10:58AM
Description: Rio Magdalena at La Providencia just off Hwy 15 at railroad crossing (km mark 223), on W side; ~10 km S of La Atascosa. A clear water cienega with many macrophytes. A good crosscut ran through its center portion and across road. Several photos taken. Site is just N. of La Providencia and Estacion Cumeral. The cienega ranges from running to swamps with rushes. It is bordered by willows and trees that look like aspens.
Water: clear
Vegetation: macrophytes
Bottom: sand and mud
Shore: mud

Field No.: VM84-10
Collectors: Robert C. Vrijenhoek and Gary Meffe
Date: 27 March 1984
Time: 12:09PM
Description: Mainstream of Rio Magdalena running adjacent to La Providencia Cienega. Much current; got almost entirely minnows in current. Got Poeciliopsis in periphery. There were no still pools but the Poeciliopsis were at the edge.
Water: clear
Vegetation: none
Bottom: gravel

Field No.: VD86-6
Collectors: Robert C. Vrijenhoek and Michael Douglas
Date: 15 November 1986
Description: La Providencia Cienega. Took a sample from undercuts of bank of stream as it passes through the cienega												

Field No.: VD86-7
Collectors: Robert C. Vrijenhoek and Michael Douglas
Date: 15 November 1986
Description: La Atascosa Cienega. We collected in an open marshy area about 200m from the road.	

Field No.: VD86-8
Collectors: Robert C. Vrijenhoek and Michael Douglas
Date: 15 November 1986
Description: La Cieneguita.  We stopped at a railroad crossing 30 km south of Nogales. Road goes into river bed of Rio Bambuto.
Time: 5:29PM

Field No.: VQH89-1
Collectors: Robert C. Vrijenhoek,  Joseph Quattro and Dean Hendrickson
Date: 8 November 1989
Time: 1-2:15PM
Description: La Providencia. (Sample a) from marshy area adjacent to mainstream. The fish were small but many pups were observed. (Sample b)  taken in mainstream from an undercut under a tree root. Many large Poeciliopsis were seen both in the mainstream and at the edges. The number of pups was unbelievable.

Field No.: VS94-1
Collectors: Robert C. Vrijenhoek and Oris Sanjur
Date: 29 September 1994
Time: 1-3PM
Description: La Providencia Cienega. It was a warm day with full sun. The fish were not visible in the mainstream. Many small fish in the periphery.  New fry. Saw Poeciliopsis in marginal weeds. We went upstream and picked fish out of the emergent vegetation.  Much Potamogeton.  Most fish were associated with the weeds.  A sample of very small fish was taken off the channel. About 1 km north of marker Km 221, close to sign "Rancho Los Alisos". North of Estacion Cumeral.

Field No.: MVH99-1
Collectors: Mariana Mateos, Robert C. Vrijenhoek and Luis A. Hurtado
Date: 20 April 1999
Description: Off Hwy 15 at Rancho Las Playas, about 50km south of Nogales. The site is on road to Ranch across the railroad tracks. The first pool and spring area at the vado, where prior collections were made, were completely dry. We passed to mainstream area which had water but no fish. We saw cormorant feeding upstream about 200 meters. We crossed through gate and found fish upstream. (Sample a) from under cut at bank in shade of tree. (Sample b) from mainstream pool with little current. This site had deeper water than 1a (~2ft deep) and there were probably more P. occidentalis here; big ones.


Field No.: MV00-1
Collectors: Mariana Mateos and Robert C. Vrijenhoek
Date: 5/1/2000
Time 11:15 AM
Description: La Providencia Cienega, N 30°55.157’ W 110°51.637 Alt 3157 (recorded with handeld GPS). Turn west at railroad crossing by Rancho Alisos. Just before Cumeral to the south. 33.5 miles South of Nogales. We sampled the stream about 100 m above road crossing and again about 400 m upstream. The water was clear and running slowly in the mainstream (MV00-1A). A peripheral undercut covered in duckweed was also sampled (MV00-1B). Green sunfish sampled from 1B, largest about 6”; froze one small individual. Saw Green Sunfish (largest ~6"); saw many large (6”) Rana tadpoles. Upstream site (MV00-1C) was source of the running water. Many Poeciliopsis among floating filamentous algae at periphery of pool. Further upstream, stream was deeper and wider. It had mostly minnows.
Water: Clear and deep (up to 18” in places)
Bottom: gravel. 


Field No.: MV01-1
Collectors: Mariana Mateos and Robert C. Vrijenhoek
Date: 17 January 2001
Time: 8:45AM
Description: La Atascosa 30°58.933’N, 110 52.365’W (recorded with handeld GPS). Small hot spring just beside highway. There were 18 Poeciliopsis and ~3 minnows. Pool is just about 1m in diameter. It drains into a marsh that is very extensive and deep with many reeds. We were unable to sample it.	
																					
Field No.: MV01-2
Collectors: Mariana Mateos and Robert C. Vrijenhoek
Date: 17 January 2001
Description: Semarnap ranch (Criadero exte. de fauna silvestre, Reg, 051, Semarnap/Profepa) at La Atascosa area 30° 58.823’ N, 110° 52.347’W (recorded with handeld GPS). Stream crosses under N and S lanes of Hwy 15. Collected below highway. Couldn’t see fish, until we seined. Water was very cold.

																					
Field No.: MV01-3
Collectors: Mariana Mateos and Robert C. Vrijenhoek
Date: 18 January 2001
Description: same as MV01-2, but also sampled stretch above highway. Ranch looks like a cattle ranch. We saw some people and asked if we could go in. Saw a couple of dark Poeciliopsis males.	
