## Supplementary material for "Environmental DNA analysis to search for an endangered desert fish": Dataset_S13.txt

Dataset S13 
Casey B. Dillman’s Collection Records

Excerpts from collection records Casey B. Dillman, Richard L. Mayden and John Hatch in April, 2009 at two sites in the Rio De La Concepción


CBD09-15
Rio de la Concepcion, Sonora, Mexico (31 11' 01.37'' ; 110 59' 05.18''). Site: at low-water bridge on Hwy 15 to Saric, the water was clear but appeared polluted, local swimming hole. No current, stream width 10', depth 1'. Bottom with sand, gravel and cobble. No cover vegetation. Species collected: Agosia chrysogaster and Gila ditaenia.


CBD09-16
Arroyo Atascosa, Sonora, Mexico (30 57' 48.79'' ; 110 52' 3.18''). Site: next Hwy 15, headed south from Nogales km 233-232. Water clear to tannin stained. Slow current, stream width 3', depth 3'. Bottom sand and cobble. Dense cover vegetation. 
Species collected: Agosia chrysogaster, Gila ditaenia and “Gambusia”.

The Gila ditaenia specimens are catalogued as University of Alabama Ichthylogical Collection, AL, USA. (UAIC) 15299.02 {Schonhuth, 2018 #12452}.

The “Gambusia” specimens (n=10) were originally catalogued as Saint Louis University Museum, MO, USA (STL) 15299.04, but this museum collection was subsequently deaccessioned. It is unclear whether specimens under STL 15299.04 were transferred to the UAIC or Cornell University Museum of Vertebrates (CUMV). Consequently, it is not possible at present to confirm whether these specimens correspond to Gambusia (and thus, the earliest record of Gambusia in the Rio De La Concepcion) or to a native topminnow (e.g. Poeciliopsis occidentalis, Poeciliopsis jackschultzi, Poeciliopsis monacha-occidentalis, or Poeciliopsis monacha-jackschultzi).
