## Supplementary material for "Environmental DNA analysis to search for an endangered desert fish": Dataset_S14.docx

**Supporting Photographs/Videos Descriptions and figshare links**

1. 2022 Rancho el Aribabi Cienega MV22-01; site 2 10.6084/m9.figshare.30644246
2. <https://figshare.com/s/3794a5811772fd721143>
3. 2022 Rancho El Aribabi mainstream MV22-02; site 2 10.6084/m9.figshare.30644342 <https://figshare.com/s/f689658b680ee2476a7a>
4. 2022 Cuitaca MV22-03; site 1
5. 10.6084/m9.figshare.30643943 <https://figshare.com/s/5d44f24ec875add7cd7c>
6. 2022 Imuris-Cocospera area MV22-04; site 4 10.6084/m9.figshare.30644477 <https://figshare.com/s/41ac48bc0db764ee8a44>
7. 2022 Imuris-Bambuto area MV22-05; site 5 10.6084/m9.figshare.30644468 <https://figshare.com/s/c88160d9acedd149a7c2>
8. 2022 La Atascosa area MV22-06; site 7 10.6084/m9.figshare.30653090 <https://figshare.com/s/64bc9e22d7a9ed228639>
9. 2022 Confluence of Cocospera with Alisos-Bambuto area MV22-07; site 4 10.6084/m9.figshare.30653075 <https://figshare.com/s/8144a9187707f646c15d>
10. 2023 Imuris-Bambuto area around pond/sidestream MV23-01; site 5 10.6084/m9.figshare.30653060 <https://figshare.com/s/5f7380d134d5329c58a0>
11. 2023 La Mesa area MV23-02; site 10 10.6084/m9.figshare.30652898 <https://figshare.com/s/f60e8255bd361e9a94d1>
12. 2023 Agua Caliente area MV23-03; site 6 10.6084/m9.figshare.30649319 <https://figshare.com/s/542b4332990efffece62>
13. 2023 Imuris-Bambuto mainstream site MV23-04 and upstream area; site 5 10.6084/m9.figshare.30656780 <https://figshare.com/s/2969da9f2a8b09f3268d>
14. 2023 Parque Mascareñas area MV23-05; site 12 10.6084/m9.figshare.30651926 <https://figshare.com/s/b6e4888f05d4d8eeda8f>
15. 2023 La Arizona area MV23-06; site 11 10.6084/m9.figshare.30652877 <https://figshare.com/s/e2278b8cc8a04451a06e>
16. 2023 El Cíbuta area  MV23-07; site 9 10.6084/m9.figshare.30653027  <https://figshare.com/s/70bc1b1595abf3dc10c4>
17. 2023 PTAR Los Alisos area; upstream of MV23-07 10.6084/m9.figshare.30653048 <https://figshare.com/s/f8ed3266c884c84a6cd8>
18. 2023 Imuris-Cocospera area MV23-08; site 4 10.6084/m9.figshare.30652937 <https://figshare.com/s/349688037b87215a63e8>
19. 2022 Rancho El Aribabi Mainstream area MV23-09; site 2 10.6084/m9.figshare.30652367 <https://figshare.com/s/a75c3a79eba173911200>
20. 2023 Rancho El Aribabi Cienega area MV23-10; site 2 10.6084/m9.figshare.30652307 <https://figshare.com/s/ab00f94c4e6fb2d98490>
21. 2023 La Atascosa area MV23-11; site 7 10.6084/m9.figshare.30649715 <https://figshare.com/s/019f1c33b6d2e9ccc4fa>
22. 2023 La Cieneguita area MV23-12; site 8 10.6084/m9.figshare.30651914 <https://figshare.com/s/9862319b3fc09687c84c>
23. 2023 site e in Fig. 1  = short stretch with water but no fish. 10.6084/m9.figshare.30657080 <https://figshare.com/s/4ff6b0dc7ef350585b7f>
24. 2023 site VIII in Fig. 1 dry 10.6084/m9.figshare.30657158 <https://figshare.com/s/b887941c7b843eeb940b>
25. 2000 photos of dry site II* showing aquatic habitat on 01/May/2000 /10.6084/m9.figshare.30727631 <https://doi.org/10.6084/m9.figshare.30727631>

1999 photos of dry site II* showing aquatic habitat on 20 April 1999 10.6084/m9.figshare.32307975 <https://figshare.com/s/4751c5feaa07bd97bbf8>
