## Supplementary material for "Environmental DNA analysis to search for an endangered desert fish": Protocol_S1.pdf

### Supporting Protocol S1

#### A. Description of Clean Lab for pre-PCR processing of samples

##### Summary

For the clean laboratory, this laboratory has restricted use via approval only and cannot be entered after entering any other laboratory in the building. Purchased supplies have not been used prior or in another laboratory and are directly placed into this laboratory upon delivery receipt. The benchtop in the clean lab is wiped down with bleach before and after every use and the entire floor is additionally bleached every two weeks. There is also a UV hood that is used to sterilize all plastic items and pipetors prior to every use. See below for required users' agreement and maintenance guidelines for the clean laboratory.

##### Copy of Users' agreement of this room

1. Never enter the ancient lab if you have visited a PCR/DNA sequencing lab, or have handled DNA extracts, PCR products, or specimens earlier that same day. You must take a shower and change clothes first, or do ancient DNA work another day. This includes short little visits to turn off a machine or check on the availability of machines.

2. All chemicals, supplies, reagents used in the ancient lab must be ordered new and brought to the ancient lab without first taking a trip into a PCR/sequencing lab. Purchase only presterilized tubes and aerosol resistant pipette tips.

3. All ancient tissue samples should be kept well packaged and isolated in labeled drawers/boxes in freezers while they are in the lab. No modern tissue samples (including food), DNA extracts, or PCR products are allowed in the ancient lab. After you do an ancient DNA extraction, you can store these extracts in the ancient-lab freezer, but all "post-PCR" samples should reside in your PCR/sequencing lab. Keep a separate ancient-lab notebook.

4. When you enter the lab to begin work, wear a designated lab coat and gloves. Begin setting up the UV hood with pipettors, tips, tubes, water, etc., and bleaching surfaces. Using a 10% bleach solution, wipe down the bench area and all equipment you will use and mop the floor (we mop ~every 2 weeks). Bleach is hard on equipment, so be clean but careful. Rinse the sponge and mop well so they don't fall apart every two weeks. After you've finished, fill the dishpan with bleach solution and sanitize all racks, pipettes, etc., and wipe down your work area again. Remove the trash. Any racks that leave the lab must be sanitised before they are returned.
5. Limit tissue-processing and DNA extraction to the area designated for such, i.e. separate from the PCR setup area.

6. Work slowly and thoughtfully to avoid contamination of the lab. Take all necessary precautions to avoid cross-contaminating your own samples: change gloves often, and perform multiple negative controls with each extraction or PCR.

#### Lab Maintenance Instructions

1. Put on lab coat and gloves.
2. Mop floor with bleach once every two weeks. Use large dishpan to mix water and bleach (about 2 tablespoons of bleach). Replace mop head if necessary.
3. Make sure there is fresh 10% bleach to wipe surfaces (change once a month). In bottle, put in about 50 ml of bleach and fill to the top with water to make the 10% solution.
4. Create new trash bag.

DNA Extraction

***Change gloves anytime you feel it is necessary***

5. Turn on the incubator to 55 degrees. Take out any frozen items that may need time to thaw (Yeast tRNA, etc).
6. Get all tubes and tips you will need to set up your extraction. Put the tubes in a rack and place the rack in the PCR hood. Open all tip boxes. Also think about other tools and reagents you may need (forceps, pH strips, etc)
7. Turn UV light on for 12 minutes (manufacturers recommendation).
8. While UV light is on, wipe down all counters with 10% bleach solution.
9. Place kimwipes on any counter space you will be using (pipetors, racks, tips, etc.).

10.  Any tools that are used to move tissue samples: use 10% bleach to rinse the tool. Wipe with a kimwipe.  Rinse again with 10% bleach.  Rinse 2X with water, wiping the tool dry after each rinse.

PCRs
***Change gloves anytime you feel it is necessary***

11.  Get all tubes and tips you will need to set up your PCR.  Put the tubes in a rack and place the rack in the PCR hood.  Open all tip boxes.
12.  Turn UV light on for 12 minutes (manufacturers recommendation).
13.  While UV light is on, wipe down all counters with 10% bleach solution.  Get all your calculations ready for your PCR and pull out reagents to thaw and vortex.
14.  Once UV is done, turn on fluorescent light and set up PCR under PCR hood.
***ALL REAGENTS EXCEPT DNA!  NEVER PUT DNA UNDER PCR HOOD***
15.  Place kimwipes on bench to set anything PCR related on (pipetors, racks, etc.).
16.  After PCR set up, place rack of tubes in fridge and clean up.

Cleanup
17.  Wipe down all surfaces (PCR hood too), pipetors, and anything used outside of PCR hood with 10% bleach.
18.  Place all items back where they belong.
19.  Remove trash.
20.  Take out trash and reaction tubes, turn off light, and shut door.

Rack Cleaning
Use a dishpan from the ancient lab and soak all racks in bleach for 20 minutes (do this in a bathroom). Use the same mixture as you would if you were mopping the floor.

#### B. Protocols for PCR Reaction first round

##### Reagents and volumes:

Forward and reverse primer at 10uM concentrations
DNA template: 4.4 µL
Platinum super Fi II Master Mix: 5 µL
BSA: 1.25 µl
F primer: 0.2 µL
R primer: 0.2 µL

Total PCR volume = 11.05 µL

##### Thermocycler settings:

Initial denaturation: 95°C - 2min

38 cycles of of the following protocol:

1. 95°C - 15s
2. 62°C - 15s
3. 72°C - 15s

Final elongation: 72°C - 2min

### C. Protocols for PCR Reaction second round

#### Reagents and volumes:

DNA template: 2 μL (undiluted first round PCR reaction)
Water: 3 μL
Platinum super Fi II Master Mix: 7.5 μL
F primer: 0.75 μL
R primer: 0.75 μL

Total PCR volume = 14 μL volume

#### Thermocycler settings:

Initial denaturation: 98°C - 2min

10 cycles of the following "touch-down" annealing protocol

1. 98°C - 15s
2. 65°C - 15s (decreased -0.5°C per cycle
3. 72°C - 15s

28 cycles of of the following protocol:

1. 98°C - 15s
2. 60°C - 15s
3. 72°C - 15s

Final elongation: 72°C - 2min
