## Supplementary material for "Environmental DNA analysis to search for an endangered desert fish": Cutadapt_commands_Batch2.txt

#! /bin/bash

# First round of cutadapt
# change filenames in last line to match input sequence file names

py -m cutadapt -j 0 -e 0 --no-indels --pair-adapters -g file:MV23Set1_indexes_anchor_cutadapt_F.fasta -G file:MV23Set1_indexes_anchor_cutadapt_R.fasta -o {name}-round1.1.fastq.gz -p {name}-round1.2.fastq.gz EDNA23_R1_001.fastq.gz EDNA23_R2_001.fastq.gz

# Second round of cutadapt
# takes all sequences not trimmed in the first round and searches in reverse
# identifies and trims sequences from reverse orientation into forward orientation 

py -m cutadapt -j 0 -e 0 --no-indels --pair-adapters -g file:MV23Set2_indexes_anchor_cutadapt_F.fasta -G file:MV23Set2_indexes_anchor_cutadapt_R.fasta -o {name}-round2.1.fastq.gz -p {name}-round2.2.fastq.gz unknown-round1.2.fastq.gz unknown-round1.1.fastq.gz
