## Supplementary material for "Environmental DNA analysis to search for an endangered desert fish": Heterandria_mitochondrion_assembly.txt

Pipeline to assemble Heterandria formosa MiFish region from ERR2814814

-Bowtie2 (usegalaxy.eu) to map Trimmomatic reads to P. occidentalis mt genome NC_028294.1
-Samtoolsview (usegalaxy.eu)was used to filter the mapped reads
-Geneious was used to visualize mapped reads and generated a consensus of 387bp of  MiFish + adjacent nucleotides 
-Bowtie2 to map Trimmomatic reads to new 387bp consensus to verify accuracy. The final consensus of the MiFish region had a coverage of at least 1000X.
-Pipeline to assemble and annotate the mitochondrial genome of Poeciliopsis jackschultzi 
-Bowtie2 was used to map Trimmomatic reads to P. occidentalis mt genome NC_028294.1 Samtoolsview was used to filter the mapped reads
-Mapped reads were de novo assembled with Geneious Assembler (settings =low sensitivity/fastest) to generate strict consensus (draft 1)
-Bowtie2 was used to map Trimmomatic reads to draft 1. Samtoolsview was used to filter the mapped reads
-Mapped reads were de novo assembled with Geneious Assembler (settings =low sensitivity/fastest) to generate strict consensus (draft final)
-The final draft had a coverage of at least 4000X, and was subjected to gene annotation with the webserver Mitofish v2025.06 [1,2,3].

1. Iwasaki, W., Fukunaga, T., Isagozawa, R., Yamada, K., Maeda, Y., Satoh, T. P., Sado, T., Mabuchi, K., Takeshima, H., and Miya, M. 2013. MitoFish and MitoAnnotator: a mitochondrial genome database of fish with an accurate and automatic annotation pipeline. Mol. Biol. Evol. 30, 2531-2540. 
2. Sato, Y., Miya, M., Fukunaga, T., Sado, T., and Iwasaki, W. 2018. MitoFish and MiFish pipeline: A mitochondrial genome database of fish with an analysis pipeline for environmental DNA metabarcoding. Mol. Biol. Evol. 35, 1553-1555.  (10.1093/molbev/msy074)
3. Zhu, T., Sato, Y., Sado, T., Miya, M., and Iwasaki, W. 2023. MitoFish, MitoAnnotator, and MiFish Pipeline: updates in 10 years. Mol. Biol. Evol. 40, msad035. 
