## Supplementary material for "Environmental DNA analysis to search for an endangered desert fish": Instructions_for_cutadapt_example_Batch2.txt

Instructions for using cutadapt to demultiplex dual indexed reads from HiSeq fastq.gz files:
1.	Install cutadapt 
2.	In the Excel spreadsheet “Index conversion for cutadapt.xlsx” set samples names in tabs “Set1_F” and “Set1_R” in column C, “Sample”
3.	Copy FASTA style indexes and names from column J “Indexes for cutadapt”. 
a.	Paste data from worksheet “Set1_F” into a text file named “Set1_indexes_anchor_cutadapt_F.fasta”.
b.	Paste data from worksheet “Set1_R” into a text file named “Set1_indexes_anchor_cutadapt_R.fasta”
4.	Set input file names in the script file “cutadapt_index_anchor.sh”
5.	In the Excel spreadsheet “cat for index.xlsx” set samples names in tabs “cat Set1.1” and “cat Set1.2”
6.	Copy cat commands from column G,  “cat commands” from both worksheet in the file “cat for index.xlsx”. Paste cat commands into the script file “cat_cutadapt.sh”.
7.	Run the script to concatenate reads from both rounds of cutadapt into single fastq.gz files, and discard redundant files.

Batch 2 specific: 
renamed each file of input sequences:
Mateos-lab-eDNA-Dec-2023_S1_L001_R1_001.fastq.qz 
to
EDNA23_R1_001.fastq.qz

Mateos-lab-eDNA-Dec-2023_S1_L001_R2_001.fastq.qz
to
EDNA23_R2_001.fastq.qz

Once all files are edited to account for name change and samples, ran according to the Cutadapt_commands_Batch2.txt  file and proceeded with above instructions.
After running the first file to separate sequences according to the unique barcode (index + primer sequence - this removes the index and primer in the output), then run the concatenate commands, according to the 
Concat_post_cutadapt_Batch2.windows.txt file.
