## Supplementary material for "Environmental DNA analysis to search for an endangered desert fish": P_jackschultzi_reference_filtering.txt

Protocol to map post-cutadapt reads to the MiFish region reference sequence of Poeciliopsis jackschultzi to determine that no reads from Poeciliopsis jackschultzi were present in the eDNA reads

Batch map to Poeciliopsis jackschultzi reference within geneious

Imported cutadapt output files into geneious based on sample read 1 and 2 files, and
paired them during import. 

Ran "batch map" on the sample files separately that had been paired during import.

In some cases, the sample files were too large for geneious to map all of the reads at once - in this case the reads of the sample file were separated into smaller "batches" and then the "batch map" was ran on each of the smaller batches.

Visually examined all reads that mapped to determine whether there were any perfect matches to the reference for both the forward and reverse reads of each read pair mapped.

The following settings were used in Geneious with "custom sensitivity" in the options menu
	Under method - interate up to 5 times
	Advanced settings
		The following boxes were checked:
		Minimum mapping quality - 30
		Trim paired read overhands
		Allow gaps- maximum per read 15%
		Ignore words repeated more than 20 times
		Accurately map reads with errors to repeat regions
		Other settings include:
		Minimum support for structural variant discovery - 2 reads
		Word length - 14
		Maximum mismatches per read - 1%
		Map multiple best matches - To none
		Maximum gap size - 50
		Index word length - 12
		Maximum abiguity - 4
