## Supplementary material for "Environmental DNA analysis to search for an endangered desert fish": Supporting_Protocol_S2_Metadata.txt

Metadata for Supporting Protocol S2. Bioinformatics Pipelines


File Instructions_for_cutadapt_example_Batch2.txt: provides instructions for the order and rationale of implementing commands in files Cutadapt_commands_Batch2.txt and Concat_post_cutadapt_Batch2.windows.txt

File: Cutadapt_commands_Batch2.txt: command line to run the two rounds of cutadapt for Batch 2.

File Concat_post_cutadapt_Batch2.windows.txt: command lines used to concatenate the output from rounds 1 and 2 of cutadapt Batch 2 (in Windows OS)

File Concat_post_cutadapt_Batch1.txt: command lines used to concatenate the output from rounds 1 and 2 of cutadapt for Batch 1 (in Mac OS Terminal)

File QiimePipeline13May2026.txt: Command lines used to run qiime with the concatenated files from cutadapt output of Batch 1 and Batch 2 together

File manifest_73.tsv: file used by Qiime Pipeline; contains the relation of each of the 73 samples and its respective forward and reverse read files

File Heterandria_mitochondrion_assembly.txt: pipeline to assemble the mitochondrial genome of Heterandria formosa to extract the MiFish region for comparison.

File P_jackschultzi_reference_filtering.txt: protocol to map post-cutadapt reads to the MiFish region reference sequence of Poeciliopsis jackschultzi to determine that no reads from Poeciliopsis jackschultzi were present in the eDNA reads
