## Supplementary material for "Environmental DNA analysis to search for an endangered desert fish": Table_S1_19May2026.docx

**Table S1.** Status of native fish taxa in the study area (Gila and Concepcion), as well as several fish taxa native to the neighboring Ríos Yaqui and Sonora, which are relevant to this study. Whether or not at least one MiFish region sequence is available for comparison is indicated. P = endangered (“en peligro de extinción”); A = threatened (“amenazada”); Pr = subject to special protection (“sujetas a protección especial”); n/a = not applicable because species has not been formally described and/or because species is not native to corresponding country.

| **Latin Name** | **Common Name** | **Relevant rivers to which it is native** | **IUCN Red List status** | **Mexico status^15,16^** | **US ESA status^17^** | **Available MiFish reference sequence** |
| --- | --- | --- | --- | --- | --- | --- |
| *Gila ditaenia* | Sonora Chub | Concepcion only | VU^1^ | A | T | yes |
| *Agosia chrysogaster* | Longfin Dace | Concepcion, Gila | LC^2^ | P | Not listed | yes |
| *Agosia sp.* (not formally described) | “Mexican Longfin Dace” | Yaqui, Sonora | n/a | n/a | n/a | no |
| *Poeciliopsis occidentalis s.l.* | Gila, Yaqui, or Sonoran Topminnow | Concepcion, Gila, Yaqui, Sonora, Mátape, Mayo | LC^3^ | A | E | yes |
| *Poeciliopsis jackschultzi* | Rio Concepcion Topminnow | Concepcion (only in a short stretch) | EN^4^ | P | n/a | yes |
| *Gila sp.*^18^ (not formally described) | “Rio Concepcion Chub” | Concepcion (only in a short stretch) | n/a | n/a | n/a | no |
| *Catostomus insignis* | Sonora Sucker | Gila^19^ | LC^5^ | P | Not listed | yes |
| *Pantosteus* formerly *Catostomus*) *clarkii* | Desert Sucker | Gila | LC^6^ | Not listed | Not listed | yes |
| *Gila robusta s.l.*^7^ | Roundtail Chub (and others) | Gila | VU^8^, EN^9^, NT^10^ | P | E | yes |
| *Catostomus bernardini* | Yaqui Sucker | Yaqui | LC^11^ | Pr | Not listed | yes |
| *Gila minacae* | Mexican Roundtail Chub | Yaqui | LC^12^ | Not listed | Not listed | yes |
| *Gila purpurea* | Yaqui Chub | Yaqui | VU^13^ | P | E | yes |
| *Gila eremica* | Desert Chub | Yaqui, Sonora, Mátape | NT^14^ | Not listed | n/a | yes |

^1^[1]; ^2^[2]; ^3^[3]; ^4^[4]; ^5^[5]; ^6^[6]; ^7^see [7-10]; ^8^[11]; ^9^[12];

^10^[13]; ^11^[14]; ^12^[15]; ^13^ [16]; ^14^[17]; ^15^[18]; ^16^[19]; ^17^[20]; ^18^[21, 22];^19^While Branson *et al.* [23] reported *Catostomus insignis* at their site 10 (which appears equivalent to our site 6), we consider this to be an error.
