## Supplementary material for "Environmental DNA analysis to search for an endangered desert fish": Table_S3.pdf

**Table S3.** Sanger-sequenced specimens.
Specimens for which the MiFish region was PCR-amplified and sequenced with Sanger technology, including voucher ID, GenBank Accession number, and collection information.

| GenBank Acc. No. | Voucher ID | Species | Geographic location | Latitude Longitude | Collection Date | Other labels used |
|---|---|---|---|---|---|---|
| PX637987 | CATO_AD_0036 | *Catostomus insignis* | USA: Arizona, Aravaipa Creek, West Site | 32.87328 N 110.589346 W | 20-May-2024 | |
| PX637988 | CATO_AD_0048 | *Pantosteus* (formerly *Catostomus*) *clarkii* | USA: Arizona, Aravaipa Creek, West Site | 32.87328 N 110.589346 W | 20-May-2024 | |
| PX637989 | CATO_SO_0324 | *Pantosteus* (formerly *Catostomus*) *clarkii* | USA: Arizona, The Nature Conservancy (TNC)'s Patagonia-Sonoita Creek Preserve | 31.52489 N 110.77821 W | 30-May-2024 | |
| PX556618 | VD86-1#A1 | *Poeciliopsis jackschultzi* | Mexico: Sonora, Imuris, Arroyo Cocospera-Babasac, Rio De La Concepcion | 30.77444 N 110.85728 W | 13-Nov-1986 | |
| PX556619 | VD86-1#A3 | *Poeciliopsis jackschultzi* | Mexico: Sonora, Imuris, Arroyo Cocospera-Babasac, Rio De La Concepcion | 30.77444 N 110.85728 W | 13-Nov-1986 | |
| PX556620 | VD86-1#A10 | *Poeciliopsis jackschultzi* | Mexico: Sonora, Imuris, Arroyo Cocospera-Babasac, Rio De La Concepcion | 30.77444 N 110.85728 W | 13-Nov-1986 | |
| PX556621 | VD86-1#A11 | *Poeciliopsis jackschultzi* | Mexico: Sonora, Imuris, Arroyo Cocospera-Babasac, Rio De La Concepcion | 30.77444 N 110.85728 W | 13-Nov-1986 | |
| PX746535 | USON-1301-9 | *Gila eremica* | Mexico: Sonora, Bacanuchi River sub–basin, Bacanuchi River at Tahuichopa ford, Arizpe–Cananea road | 30.366572 N 110.156817 W | 24-Jul-2014 | G. eremicaA9 = A9 |

| GenBank Acc. No. | Voucher ID | Species | Geographic location | Latitude Longitude | Collection Date | Other labels used |
|---|---|---|---|---|---|---|
| PX746536 | USON-1376-29 | *Gila eremica* | Mexico: Sonora, Mátape River sub–basin, Mátape River at Mazatán | 29.9989 N 100.14757 W | 22-Nov-2014 | G. eremicaM29 = M29 |
| PX746537 | USON-1300-18 | *Gila cf. eremica* | Mexico: Sonora, Arroyo El Tigre sub–basin, La Balandrona Canyon, Sierra El Aguaje mountains | 28.0439 N 111.07277 W | 28-Jun-2014 | GcanonBA = BA18 |
| PX746538 | USON-1302-1 | *Gila cf. eremica* | Mexico: Sonora, Arroyo El Tigre sub–basin, La Pirinola Canyon, Sierra El Aguaje mountains | 28.092222 N 111.0375 W | 30-Aug-2014 | GcanonP = P1 |
| PX746541 | USON-1224-2 | *Gila minacae* | Mexico, Sonora, Bavispe River sub–basin, Arroyo El Largo, 2.5 km E Ejido Arroyo El Largo | 29.734417 N 108.6135 W | 02-Nov-2008 | Gmincae2 = GM2 |
| PX746539 | USON-1224-5 | *Gila minacae* | Mexico, Sonora, Bavispe River sub–basin, Arroyo El Largo, 2.5 km E Ejido Arroyo El Largo | 29.734417 N 108.6135 W | 02-Nov-2008 | Gmincae5 = GM5 |
| PX746540 | USON-1224-6 | *Gila minacae* | Mexico, Sonora, Bavispe River sub–basin, Arroyo El Largo, 2.5 km E Ejido Arroyo El Largo | 29.734417 N 108.6135 W | 02-Nov-2008 | Gmincae6 =GM6 |
| PX746530 | USON-1455-4 | *Agosia chrysogaster* | Mexico: Sonora, Arroyo Cocóspera, Rancho El Aribabi, Rio De La Concepción | 30.854167 N 110.661944 W | 18-May-2022 | Gdit4=M4x |
| PX746543 | UABC-2921-1 | *Gila sp.* | Mexico: Sinaloa, Arroyo Las Higueras at Badiraguato (469masl), Rio Culiacan | 25.633233 N 107.536022 W | 13-Dec-2011 | Gnsp1 |

| GenBank Acc. No. | Voucher ID | Species | Geographic location | Latitude Longitude | Collection Date | Other labels used |
|---|---|---|---|---|---|---|
| PX746533 | UABC-2919-3 | *Gila sp.* | Mexico: Sinaloa, Arroyo El Rodeo at Tamazula (450masl), Rio Culiacan | 24.911308 N 106.782394 W | 9-Dec-2011 | Gnsp3 |
| PX746531 | USON-1454-2 | *Gila ditaenia* | Mexico: Sonora, Arroyo Cocóspera, Rancho El Aribabi, Rio De La Concepción | 30.854167 N 110.661944 W | 18-May-2022 | GditaeniaM 2 = Gdit2 |
| PX746532 | USON-1454-3 | *Gila ditaenia* | Mexico: Sonora, Arroyo Cocóspera, Rancho El Aribabi, Rio De La Concepción | 30.854167 N 110.661944 W | 18-May-2022 | GditaeniaM 3 = Gdit3 |
| PX746534 | USON-1490-1 | *Gila sp.* | Mexico: Sinaloa,Arroyo San Juan del Llano at crossing with road Arroyo San Juan del Llano, Rio Sinaloa | 25.78211827 N 107.3225 W | 14-Mar-2025 | Gsp1 |
| PX746544 | USON-1490-2 | *Gila sp.* | Mexico: Sinaloa,Arroyo San Jose del Llano at crossing with road Arroyo San Jose del Llano, Rio Sinaloa | 25.78211827 N 107.3225 W | 14-Mar-2025 | Gsp2 |
| PX746542 | USON-1490-3 | *Gila sp.* | Mexico: Sinaloa,Arroyo San Jose del Llano at crossing with road Arroyo San Jose del Llano, Rio Sinaloa | 25.78211827 N 107.3225 W | 14-Mar-2025 | Gsp3 |
| PX746545 | USON-1490-4 | *Gila sp.* | Mexico: Sinaloa,Arroyo San Jose del Llano at crossing with road Arroyo San Jose del Llano, Rio Sinaloa | 25.78211827 N 107.3225 W | 14-Mar-2025 | Gsp4 |
| PX746546 | USON-1490-5 | *Gila sp.* | Mexico: Sinaloa,Arroyo San Jose del Llano at crossing with road Arroyo San Jose del Llano, Rio Sinaloa | 25.78211827 N 107.3225 W | 14-Mar-2025 | Gsp5 |

| GenBank Acc. No. | Voucher ID | Species | Geographic location | Latitude Longitude | Collection Date | Other labels used |
|---|---|---|---|---|---|---|
| PZ418350 | YCS09 | *Gila purpurea* | USA: Arizona, San Bernardino NWR-House Pond | 31.339781 N 109.264560 W | unknown | 09 |
| PZ418349 | YCL039 | *Gila purpurea* | USA: Arizona, Leslie Canyon NWR | 31.591332 N 109.508942 W | unknown | 039 |
| PZ418347 | YCD027 | *Gila purpurea* | USA: Arizona, Douglas High School | 31.351373 N 109.535659 W | unknown | 027 |
| PZ418348 | YCE005 | *Gila purpurea* | USA: Arizona, El Coronado Ranch | 31.868299 N 109.367675 W | unknown | 05 |
