## Supplementary material for "Environmental DNA analysis to search for an endangered desert fish": Table_S4_Detected_Observed_Expected17May2026_1.pdf

Table S4. Collective evidence used to infer presence/absence of each of 13 teleost taxa considered non-spurious. For each site/year, values = relative frequency of the sum of replicates (replicate values above "all blanks" threshold). Spurious taxa excluded from counts; and Aribabi Cienega 2022 excluded. Obs (Y/N) = whether taxon was observed during eDNA sampling. Exp (Y/N) = whether taxon is expected based on prior/nearby records. Inference = whether the taxon is inferred as present (P) or absent (A) based on the collective evidence. Red cells = deemed false positive eDNA results; Green cells = deemed false negative eDNA results.

| Latin Name | Common name (N = Native; I = Introduced) | Cuitaca upstream, MV22-3B | | | | Cuitaca downstream, MV22-3A | | | | Aribabi Cienega, MV23-10 | | | | Aribabi mainstream, MV22-2 | | | |
|---|---|---|---|---|---|---|---|---|---|---|---|---|---|---|---|---|---|
| | | relative eDNA abudance | Obs | Exp | Inf | relative eDNA abudance | Obs | Exp | Inf | relative eDNA abudance | Obs | Exp | Inf | relative eDNA abudance | Obs | Exp | Inf |
| Agosia chrysogaster | Longfin Dace (N) | 100.00 | Y | Y | P | 99.87 | Y | Y | P | 0.00 | N | Y | A | 24.02 | Y | Y | P |
| Gila ditaenia | Sonora Chub (N) | 0.00 | N | Y | A | **0.04** | N | Y | **A** | 0.00 | N | Y | A | 62.14 | Y | Y | P |
| Poeciliopsis occidentalis | Gila Topminnow (N) | 0.00 | N | Y | A | **0.09** | N | Y | **A** | 0.00 | N | Y | A | 0.00 | N | Y | A |
| Gila minacae (= "Gila not ditaenia") | Mexican Roundtail Chub (= "Chub; not Sonora") | 0.00 | N | N | A | 0.00 | N | N | A | 0.00 | N | N | A | 0.00 | N | N | A |
| P. monacha-occidentalis asexual hybrid | Topminnow Asexual Hybrid (N) | 0.00 | N | ? | A | 0.00 | N | ? | A | 0.00 | N | ? | A | 0.00 | N | ? | A |
| Catostomus bernardini | Yaqui Sucker (I?) | 0.00 | N | N | A | 0.00 | N | N | A | 0.00 | N | N | A | 0.00 | N | N | A |
| Lepomis cyanellus | Green Sunfish (I) | 0.00 | N | Y | A | 0.00 | N | Y | A | 59.02 | N | Y | P | 11.82 | N | Y | P |
| Cichlidae sp. | Tilapia (I) | 0.00 | N | Y | A | 0.00 | N | Y | A | 0.00 | N | Y | A | 0.00 | N | Y | A |
| Ameiurus melas | Black Bullhead (I) | 0.00 | N | Y | A | 0.00 | N | Y | A | 0.00 | N | Y | A | 0.00 | N | Y | A |
| Gambusia affinis | Mosquitofish (I) | 0.00 | N | Y | A | 0.00 | N | Y | A | 0.00 | N | Y | A | 0.00 | N | Y | A |
| Lepomis macrochirus | Bluegill (I) | 0.00 | N | Y | A | 0.00 | N | Y | A | 24.62 | N | Y | P | 2.03 | N | Y | P |
| Micropterus salmoides | Largemouth Bass (I) | 0.00 | N | Y | A | 0.00 | N | Y | A | 16.36 | N | Y | P | 0.00 | N | Y | A |
| Xiphophorus hellerii | Green Swordtail (I) | 0.00 | N | N | A | 0.00 | N | N | A | 0.00 | N | N | A | 0.00 | N | N | A |
| | proportion of natives (%) | 100% | | | | 100% | | | | 0% | | | | 86% | | | |
| | mean proportion of natives | 69% | | | | | | | | | | | | | | | |
| | Standard deviation of proportion of natives | 32% | | | | | | | | | | | | | | | |

| Latin Name | Common name (N = Native; I = Introduced) | Aribabi mainstream, MV23-9 | | | | Imuris-Cocospera, MV22-4 | | | | Imuris-Cocospera, MV23-8 | | | | Bambuto-Cocospera, MV22-7 | | | | Imuris-Bambuto mainstream, MV22-5A | | | |
|---|---|---|---|---|---|---|---|---|---|---|---|---|---|---|---|---|---|---|---|---|---|
| | | relative eDNA abudance | Obs | Exp | Inf | relative eDNA abudance | Obs | Exp | Inf | relative eDNA abudance | Obs | Exp | Inf | relative eDNA abudance | Obs | Exp | Inf | relative eDNA abudance | Obs | Exp | Inf |
| Agosia chrysogaster | Longfin Dace (N) | 35.35 | Y | Y | P | 3.13 | Y | Y | P | 3.53 | Y | Y | P | 28.55 | N | Y | P | 3.16 | Y | Y | P |
| Gila ditaenia | Sonora Chub (N) | 19.35 | Y | Y | P | 41.72 | Y | Y | P | 30.32 | Y | Y | P | 19.09 | N | Y | P | 34.68 | Y | Y | P |
| Poeciliopsis occidentalis | Gila Topminnow (N) | 0.00 | N | Y | A | 15.81 | Y | Y | P | 15.31 | Y | Y | P | 34.23 | Y | Y | P | 0.95 | N | Y | P |
| Gila minacae (= "Gila not ditaenia") | Mexican Roundtail Chub (= "Chub; not Sonora") | 0.00 | N | N | A | 0.00 | N | N | A | 0.00 | N | N | A | 0.00 | N | N | A | 0.00 | N | N | A |
| P. monacha-occidentalis asexual hybrid | Topminnow Asexual Hybrid (N) | 0.00 | N | ? | A | 11.75 | Y | Y | P | 7.54 | Y | Y | P | 10.09 | Y | Y | P | 1.17 | N | Y | P |
| Catostomus bernardini | Yaqui Sucker (I?) | 0.00 | N | N | A | 0.00 | N | N | A | 0.00 | N | N | A | 0.00 | N | N | A | 0.00 | N | N | A |
| Lepomis cyanellus | Green Sunfish (I) | 41.02 | N | Y | P | 3.83 | N | Y | P | 14.83 | N | Y | P | 0.27 | N | Y | P | 34.48 | Y | Y | P |
| Cichlidae sp. | Tilapia (I) | 0.00 | N | Y | A | 17.60 | N | Y | P | 13.78 | N | Y | P | 5.06 | N | Y | P | 18.60 | Y | Y | P |
| Ameiurus melas | Black Bullhead (I) | 0.00 | N | Y | A | 0.45 | N | Y | P | 0.45 | N | Y | P | 0.72 | N | Y | P | 6.61 | N | Y | P |
| Gambusia affinis | Mosquitofish (I) | 0.00 | N | Y | A | 0.04 | N | Y | P | 0.95 | N | Y | P | 1.43 | N | Y | P | 0.29 | N | Y | P |
| Lepomis macrochirus | Bluegill (I) | 4.12 | N | Y | P | 0.00 | N | Y | A | 0.00 | N | Y | A | 0.00 | N | Y | A | 0.00 | N | Y | A |
| Micropterus salmoides | Largemouth Bass (I) | 0.15 | N | Y | P | 0.00 | N | Y | A | 0.00 | N | Y | A | 0.00 | N | Y | A | 0.00 | N | Y | A |
| Xiphophorus hellerii | Green Swordtail (I) | 0.00 | N | N | A | 5.69 | Y | Y | P | 13.29 | Y | Y | P | 0.56 | Y | Y | P | 0.06 | Y | Y | P |
| | proportion of natives (%) | 55% | | | | 72% | | | | 57% | | | | 92% | | | | 40% | | | |
| | mean proportion of natives | | | | | | | | | | | | | | | | | | | | |
| | Standard deviation of proportion of natives | | | | | | | | | | | | | | | | | | | | |

| Latin Name | Common name (N = Native; I = Introduced) | Imuris-Bambuto mainstream, MV23-4 | | | | Imuris-Bambuto sidestream, MV22-5C | | | | Imuris-Bambuto sidestream, MV23-1 | | | | La Atascosa, MV22-6 | | | | La Atascosa, MV23-11 | | | |
|---|---|---|---|---|---|---|---|---|---|---|---|---|---|---|---|---|---|---|---|---|---|
| | | relative eDNA abudance | Obs | Exp | Inf | relative eDNA abudance | Obs | Exp | Inf | relative eDNA abudance | Obs | Exp | Inf | relative eDNA abudance | Obs | Exp | Inf | relative eDNA abudance | Obs | Exp | Inf |
| Agosia chrysogaster | Longfin Dace (N) | 4.38 | Y | Y | P | 6.90 | Y | Y | P | 4.41 | Y | Y | P | 31.00 | Y | Y | P | 17.43 | Y | Y | P |
| Gila ditaenia | Sonora Chub (N) | 41.55 | Y | Y | P | 22.12 | Y | Y | P | 27.30 | Y | Y | P | 52.24 | Y | Y | P | 66.69 | Y | Y | P |
| Poeciliopsis occidentalis | Gila Topminnow (N) | 0.00 | N | Y | A | 1.17 | Y | Y | P | 0.03 | Y | Y | P | 4.81 | Y | Y | P | 2.36 | Y | Y | P |
| Gila minacae (= "Gila not ditaenia") | Mexican Roundtail Chub (= "Chub; not Sonora") | 0.00 | N | N | A | 0.00 | N | N | A | 0.00 | N | N | A | 0.00 | N | N | A | 0.00 | N | Y | A |
| P. monacha-occidentalis asexual hybrid | Topminnow Asexual Hybrid (N) | 0.05 | N | Y | P | 4.01 | Y | Y | P | 0.45 | Y | Y | P | 6.33 | Y | Y | P | 2.69 | Y | Y | P |
| Catostomus bernardini | Yaqui Sucker (I?) | 0.00 | N | N | A | 0.00 | N | N | A | 0.00 | N | N | A | 0.00 | N | N | A | 0.00 | N | N | A |
| Lepomis cyanellus | Green Sunfish (I) | 33.77 | Y | Y | P | 27.78 | Y | Y | P | 21.65 | Y | Y | P | 5.57 | N | Y | P | 9.25 | N | Y | P |
| Cichlidae sp. | Tilapia (I) | 19.74 | Y | Y | P | 28.87 | Y | Y | P | 35.73 | Y | Y | P | 0.00 | N | Y | A | 0.00 | N | Y | A |
| Ameiurus melas | Black Bullhead (I) | 0.05 | N | Y | P | 0.78 | Y | Y | P | 0.03 | Y | Y | P | 0.06 | N | Y | P | 0.03 | N | Y | P |
| Gambusia affinis | Mosquitofish (I) | 0.01 | N | Y | P | 8.37 | Y | Y | P | 9.41 | Y | Y | P | 0.00 | N | Y | A | 0.04 | N | Y | P |
| Lepomis macrochirus | Bluegill (I) | 0.26 | N | Y | P | 0.00 | N | N | A | 0.00 | N | N | A | 0.00 | N | Y | A | 1.51 | N | Y | P |
| Micropterus salmoides | Largemouth Bass (I) | 0.00 | N | N | A | 0.00 | N | N | A | 0.02 | N | N | P | 0.00 | N | Y | A | 0.00 | N | Y | A |
| Xiphophorus hellerii | Green Swordtail (I) | 0.19 | Y | Y | P | 0.00 | Y | ? | A | 0.96 | Y | Y | P | 0.00 | N | N | A | 0.00 | N | N | A |
| | proportion of natives (%) | 46% | | | | 34% | | | | 32% | | | | 94% | | | | 89% | | | |
| | mean proportion of natives | | | | | | | | | | | | | | | | | | | | |
| | Standard deviation of proportion of natives | | | | | | | | | | | | | | | | | | | | |

| Latin Name | Common name (N = Native; I = Introduced) | Agua Caliente, MV23-13 | | | | La Cieneguita, MV23-12 | | | | Cibuta, MV23-7 | | | | La Mesa, MV23-2 | | | | La Arizona, MV23-6 | | | |
|---|---|---|---|---|---|---|---|---|---|---|---|---|---|---|---|---|---|---|---|---|---|
| | | relative eDNA abudance | Obs | Exp | Inf | relative eDNA abudance | Obs | Exp | Inf | relative eDNA abudance | Obs | Exp | Inf | relative eDNA abudance | Obs | Exp | Inf | relative eDNA abudance | Obs | Exp | Inf |
| Agosia chrysogaster | Longfin Dace (N) | 0.00 | N | Y | A | 0.00 | ? | Y | A | 51.00 | Y | Y | P | 78.01 | Y | Y | P | 42.57 | Y | Y | P |
| Gila ditaenia | Sonora Chub (N) | 1.17 | N | Y | P | 76.21 | ? | Y | P | 36.02 | Y | Y | P | 21.85 | Y | Y | P | 44.98 | Y | Y | P |
| Poeciliopsis occidentalis | Gila Topminnow (N) | 0.00 | N? | ? | A | 0.65 | ? | Y | P | **0.00** | Y | Y | **P** | 0.06 | Y | Y | P | 1.02 | Y | Y | P |
| Gila minacae (= "Gila not ditaenia") | Mexican Roundtail Chub (= "Chub; not Sonora") | 0.00 | N | N | A | 0.00 | N | N | A | 0.00 | N | N | A | 0.00 | N | N | A | 0.00 | N | N | A |
| P. monacha-occidentalis asexual hybrid | Topminnow Asexual Hybrid (N) | 13.07 | Y | Y | A | 16.75 | Y? | Y | P | 0.00 | ? | Y | A | 0.00 | ? | ? | A | 0.00 | ? | ? | A |
| Catostomus bernardini | Yaqui Sucker (I?) | 0.00 | N | N | A | 0.00 | N | N | A | 0.00 | N | N | A | 0.00 | N | N | A | 0.00 | N | N | A |
| Lepomis cyanellus | Green Sunfish (I) | 51.88 | N | Y | P | 4.29 | N | Y | P | 12.97 | N | Y | P | 0.07 | N | Y | P | 0.00 | N | Y | A |
| Cichlidae sp. | Tilapia (I) | 0.00 | N | Y | A | 0.00 | N | Y | A | 0.00 | N | Y | A | 0.00 | N | Y | A | **0.00** | Y | Y | **P** |
| Ameiurus melas | Black Bullhead (I) | 33.88 | N | Y | P | 1.05 | N | Y | P | 0.00 | N | Y | A | 0.00 | N | Y | A | 0.00 | N | Y | A |
| Gambusia affinis | Mosquitofish (I) | 0.00 | N | Y | A | 1.05 | N | Y | P | **0.00** | Y | Y | **P** | 0.01 | Y | Y | P | 1.47 | Y | Y | P |
| Lepomis macrochirus | Bluegill (I) | 0.00 | N | Y | A | 0.00 | N | Y | A | 0.00 | N | Y | A | 0.00 | N | Y | A | 0.65 | N | Y | P |
| Micropterus salmoides | Largemouth Bass (I) | 0.00 | N | Y | A | 0.00 | N | Y | A | 0.00 | N | Y | A | 0.00 | N | Y | A | 9.31 | Y | Y | P |
| Xiphophorus hellerii | Green Swordtail (I) | 0.00 | N | N | A | 0.00 | N | N | A | 0.00 | N | N | A | 0.00 | N | N | A | 0.00 | N | N | A |
| | proportion of natives (%) | 14% | | | | 94% | | | | 87% | | | | 100% | | | | 89% | | | |
| | mean proportion of natives | | | | | | | | | | | | | | | | | | | | |
| | Standard deviation of proportion of natives | | | | | | | | | | | | | | | | | | | | |

| Latin Name | Common name (N = Native; I = Introduced) | Sta. Cruz-Mascareñas, MV23-5 | | | | minimum relative frequency where present | maximum relative frequency | average relative frequency where present | standard deviation of relative frequency where present |
|---|---|---|---|---|---|---|---|---|---|
| | | relative eDNA abudance | Obs | Exp | Inf | | | | |
| Agosia chrysogaster | Longfin Dace (N) | 73.04 | Y | Y | P | 3.13 | 100.00 | 35.67 | 33.73 |
| Gila ditaenia | Sonora Chub (N) | **0.01** | N | N | **A** | 1.17 | 76.21 | 37.34 | 19.83 |
| Poeciliopsis occidentalis | Gila Topminnow (N) | 8.08 | Y | Y | P | 0.03 | 34.23 | 7.04 | 10.26 |
| Gila minacae (= "Gila not ditaenia") | Mexican Roundtail Chub (= "Chub; not Sonora") | 0.01 | N | Y | P | | | | |
| P. monacha-occidentalis asexual hybrid | Topminnow Asexual Hybrid (N) | 18.56 | Y? | N? | P | 0.05 | 18.56 | 7.70 | 6.36 |
| Catostomus bernardini | Yaqui Sucker (I?) | 0.08 | N | Y | P | | | | |
| Lepomis cyanellus | Green Sunfish (I) | 0.03 | N | Y | P | 0.03 | 59.02 | 19.56 | 18.71 |
| Cichlidae sp. | Tilapia (I) | 0.00 | N | ? | A | 5.06 | 35.73 | 19.91 | 9.96 |
| Ameiurus melas | Black Bullhead (I) | 0.00 | N | Y | A | 0.03 | 76.21 | 37.34 | 19.83 |
| Gambusia affinis | Mosquitofish (I) | 0.18 | N | Y | P | 0.01 | 9.41 | 1.94 | 3.30 |
| Lepomis macrochirus | Bluegill (I) | 0.00 | N | Y | A | 0.26 | 24.62 | 5.53 | 9.45 |
| Micropterus salmoides | Largemouth Bass (I) | 0.00 | N | Y | A | 0.02 | 16.36 | 6.46 | 7.90 |
| Xiphophorus hellerii | Green Swordtail (I) | 0.00 | N | N | A | 0.06 | 13.29 | 3.46 | 5.26 |
| | proportion of natives (%) | 100% | | | | | | | |
| | mean proportion of natives | | | | | | | | |
| | Standard deviation of proportion of natives | | | | | | | | |
