## Supplementary figures and images for "Environmental DNA analysis to search for an endangered desert fish"

### Figure_S1_OriginHybridoMOColor20May2026.png

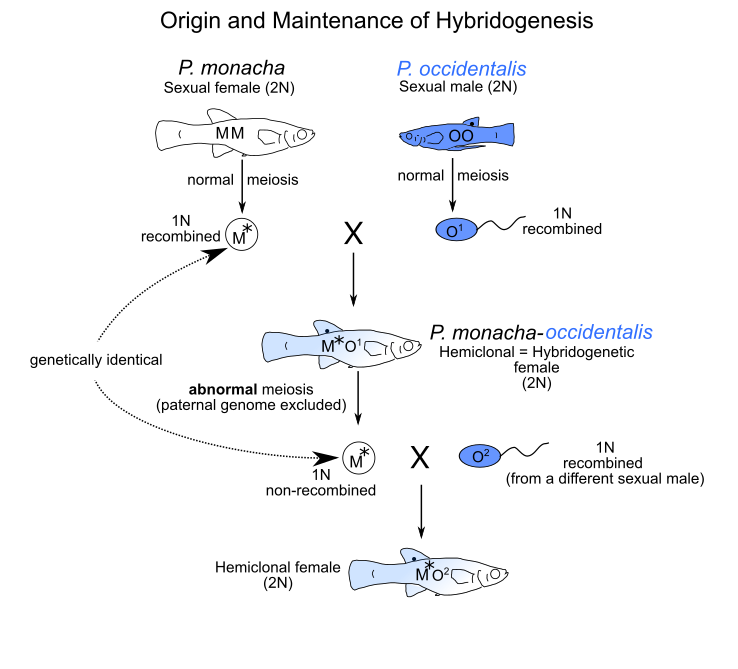

### Figure_S2_Absolute_all_all_03Jul2025.pdf

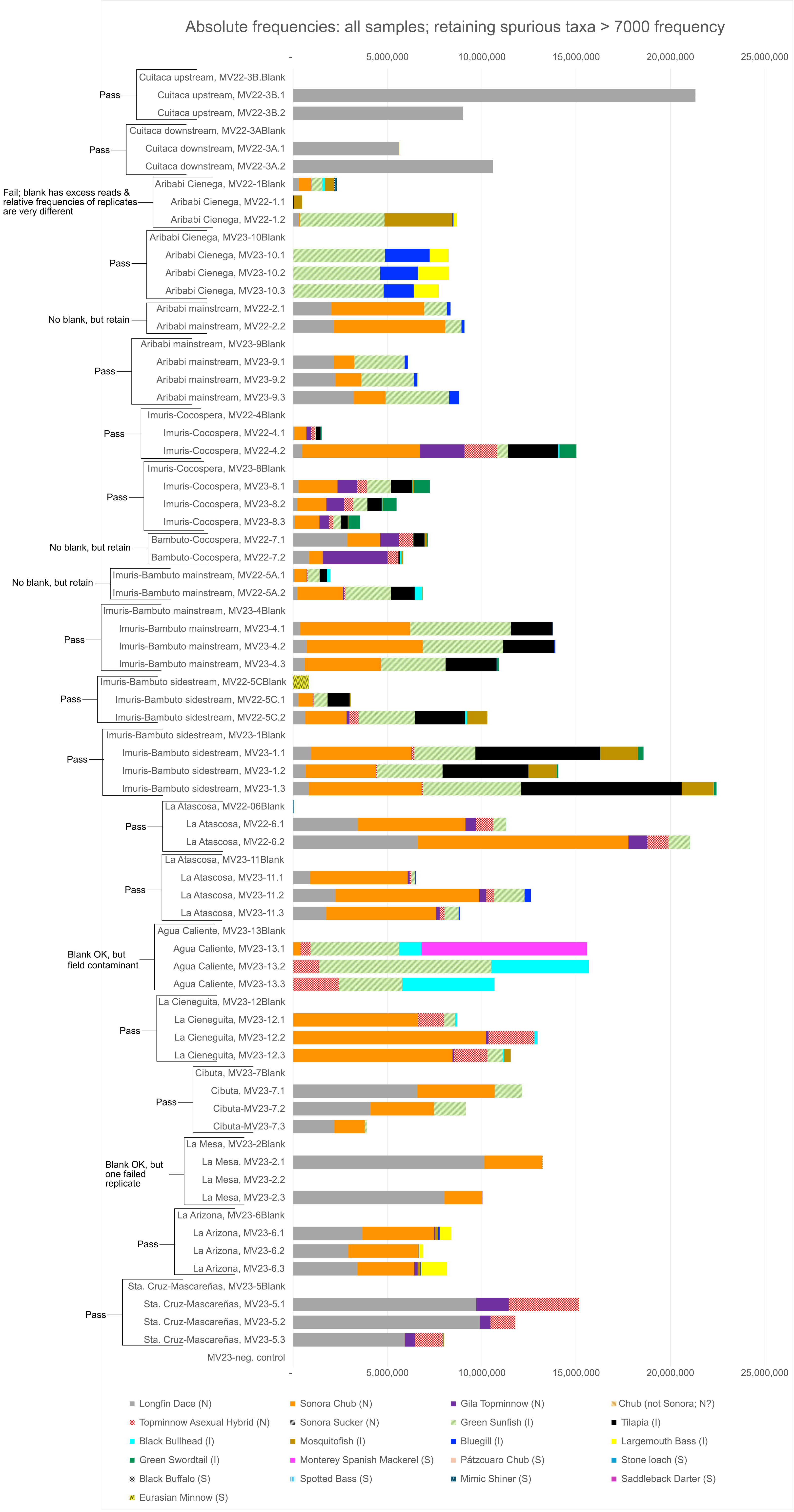

### Figure_S3_Leuciscidae_805_02Apr2026_75_gapsstripped_p_nj_pars_brlens_colors.tree.pdf

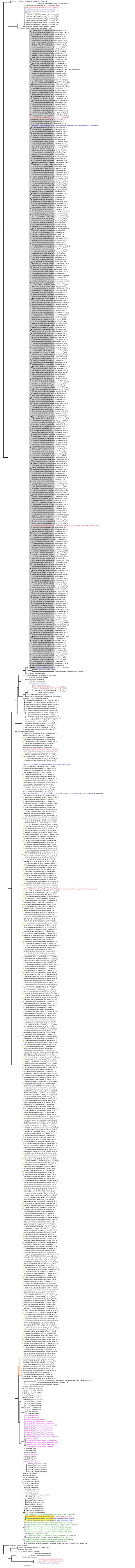

### Figure_S4_Feature_Frequency_Dist_within_species.pdf

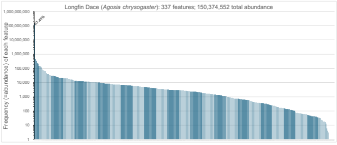

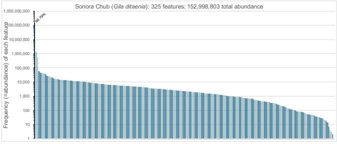

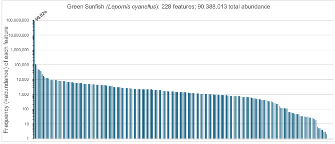

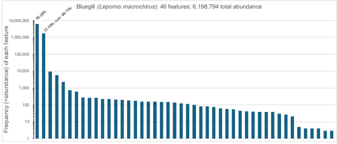

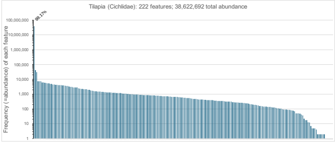

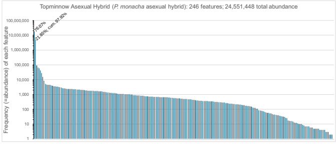

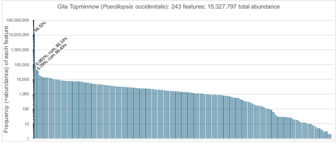

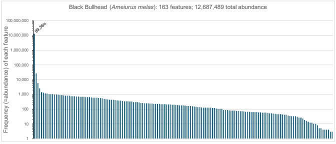

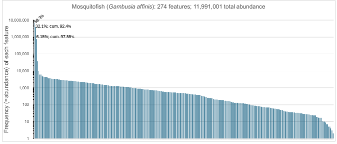

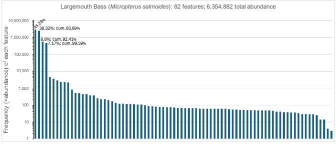

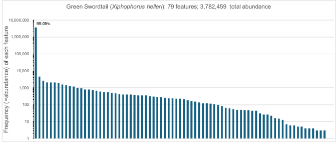

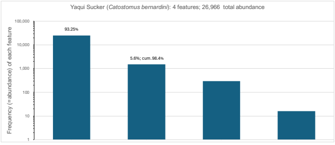

### Figure_S5_Poeciliidae_features_plus_references14Nov2025_737taxa_NJ_JC_brlens_pars_colors.trees.pdf

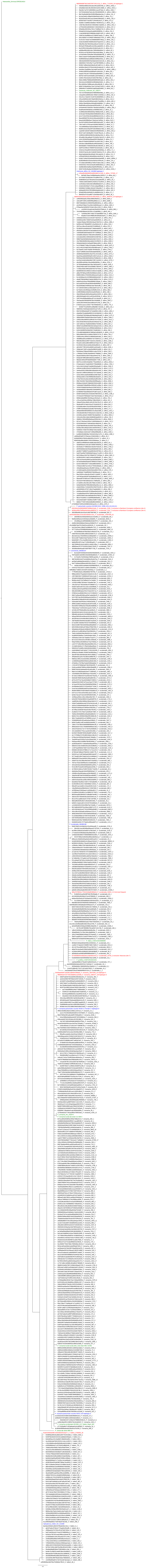

### Figure_S6_SuckerMiFish_125taxa_170bp_p_dist_NJ_parsbrlens_colors13Nov2025.tree.pdf

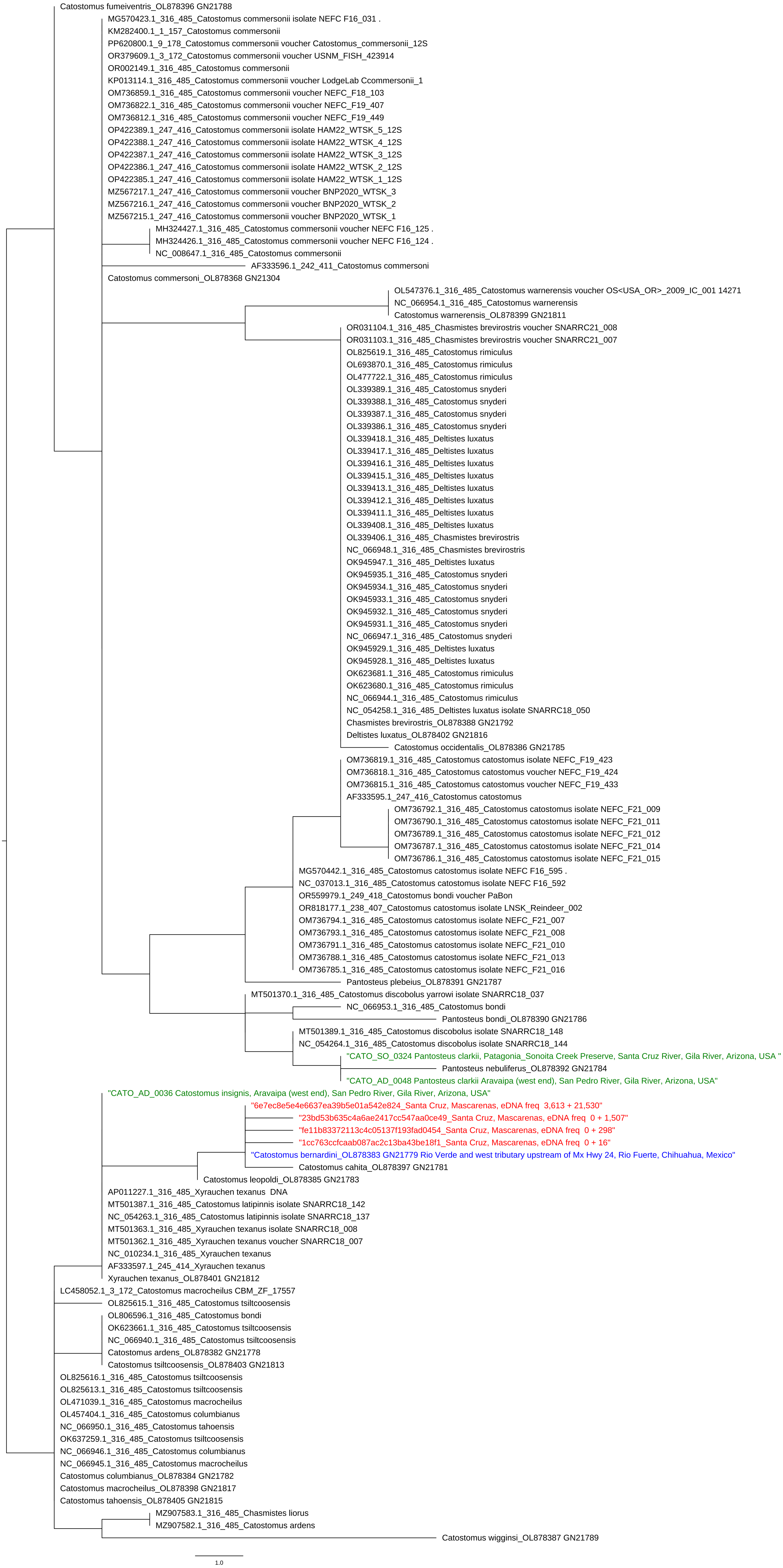

### Figure_S7_monacha_hap_a_d_freqs29Jul2025.pdf

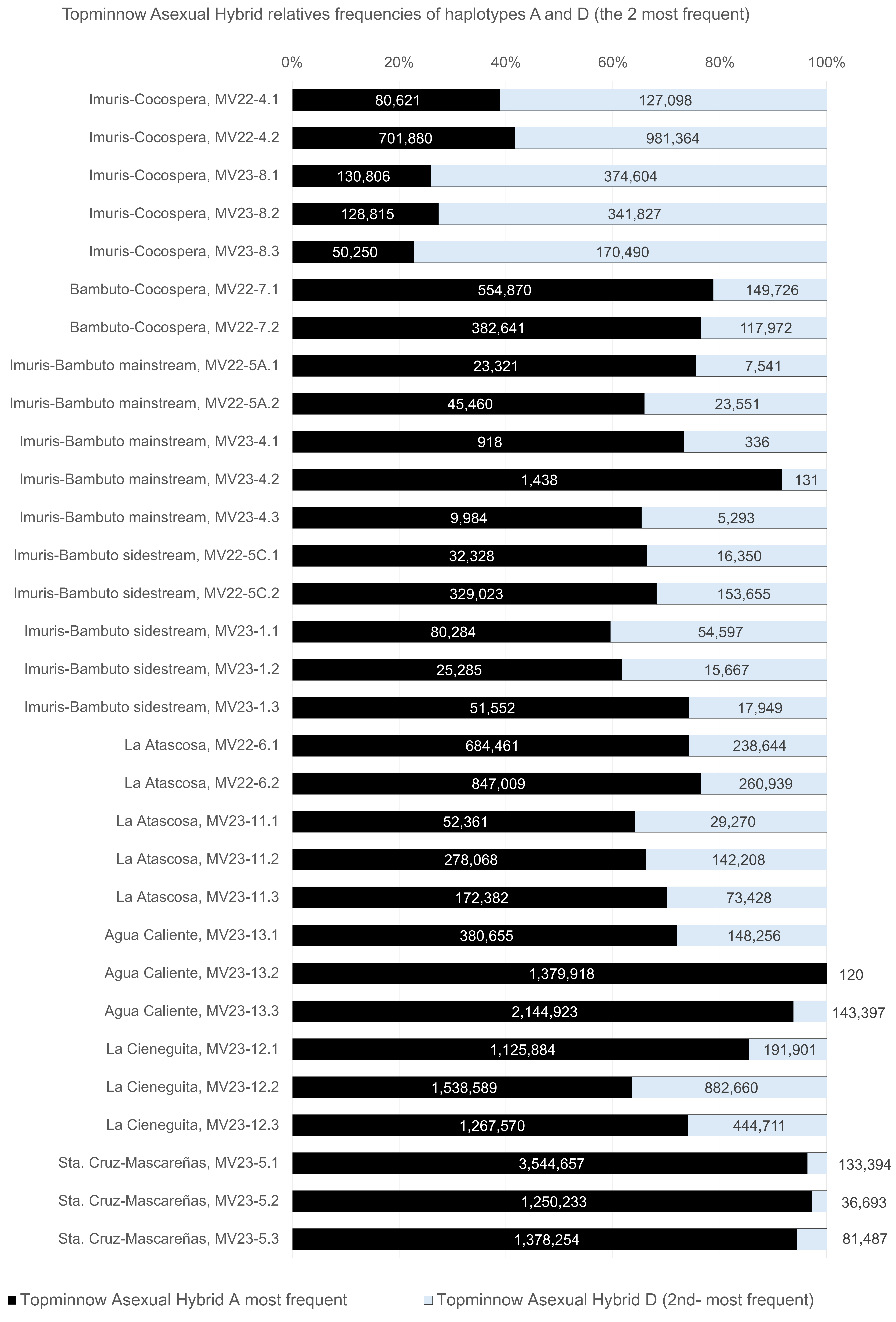

### Figure_S8_occidenta_top_3_feature_freqs29Jul2025.pdf

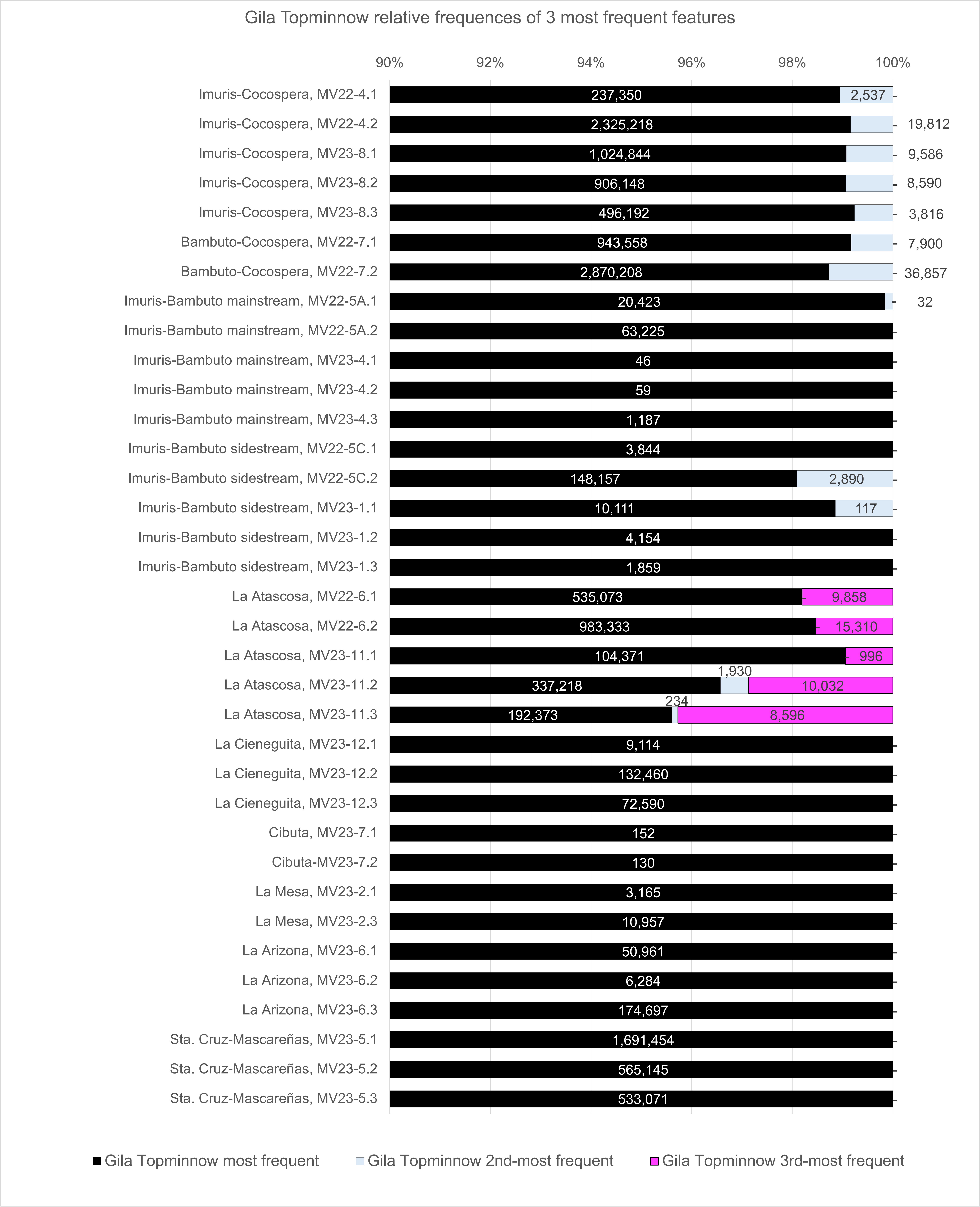

### Figure_S9_Agosia_most_frequent_feat25Nov2025.pdf

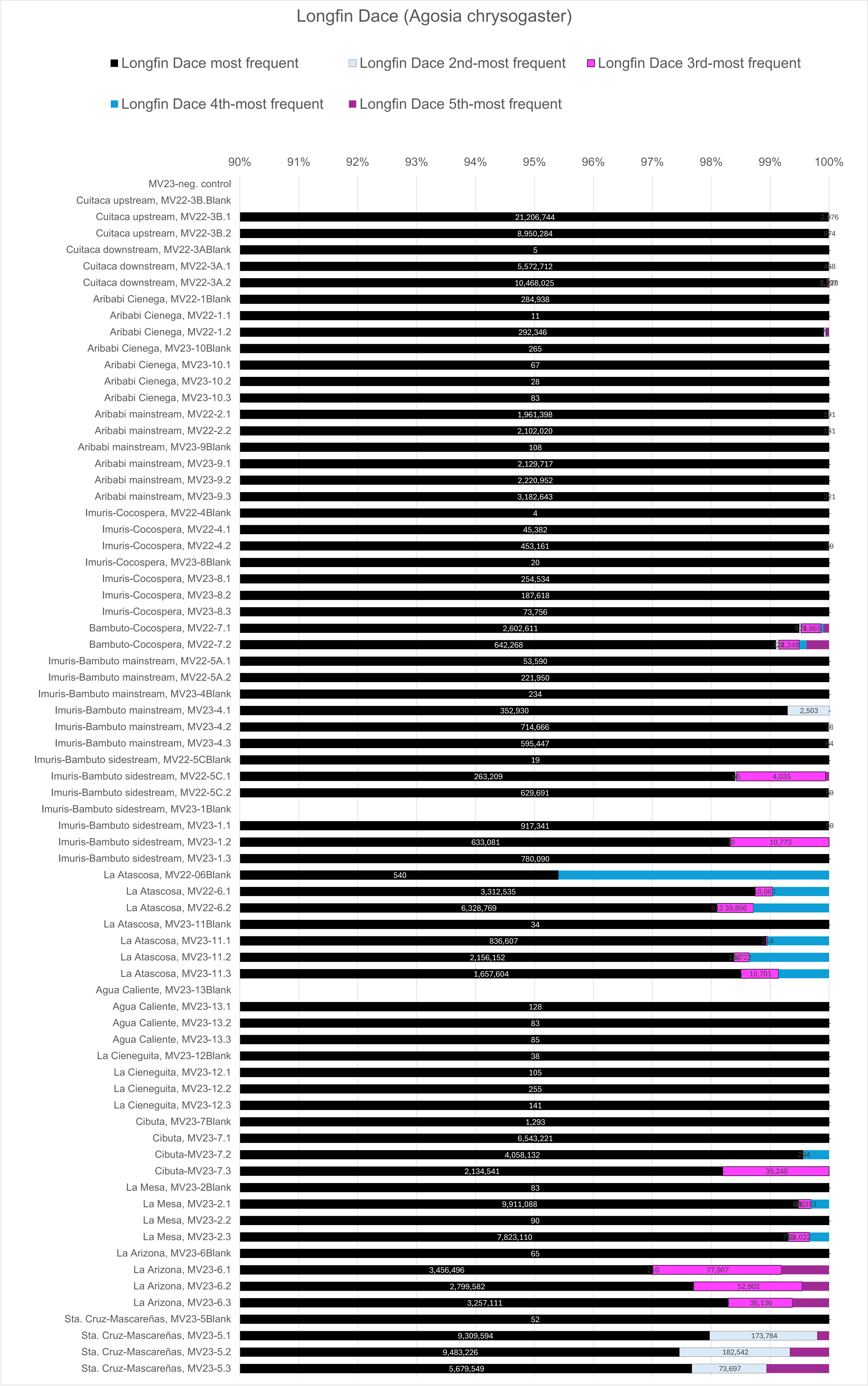

### Figure_S10_Gila_most_frequent_feat25Nov2025.pdf

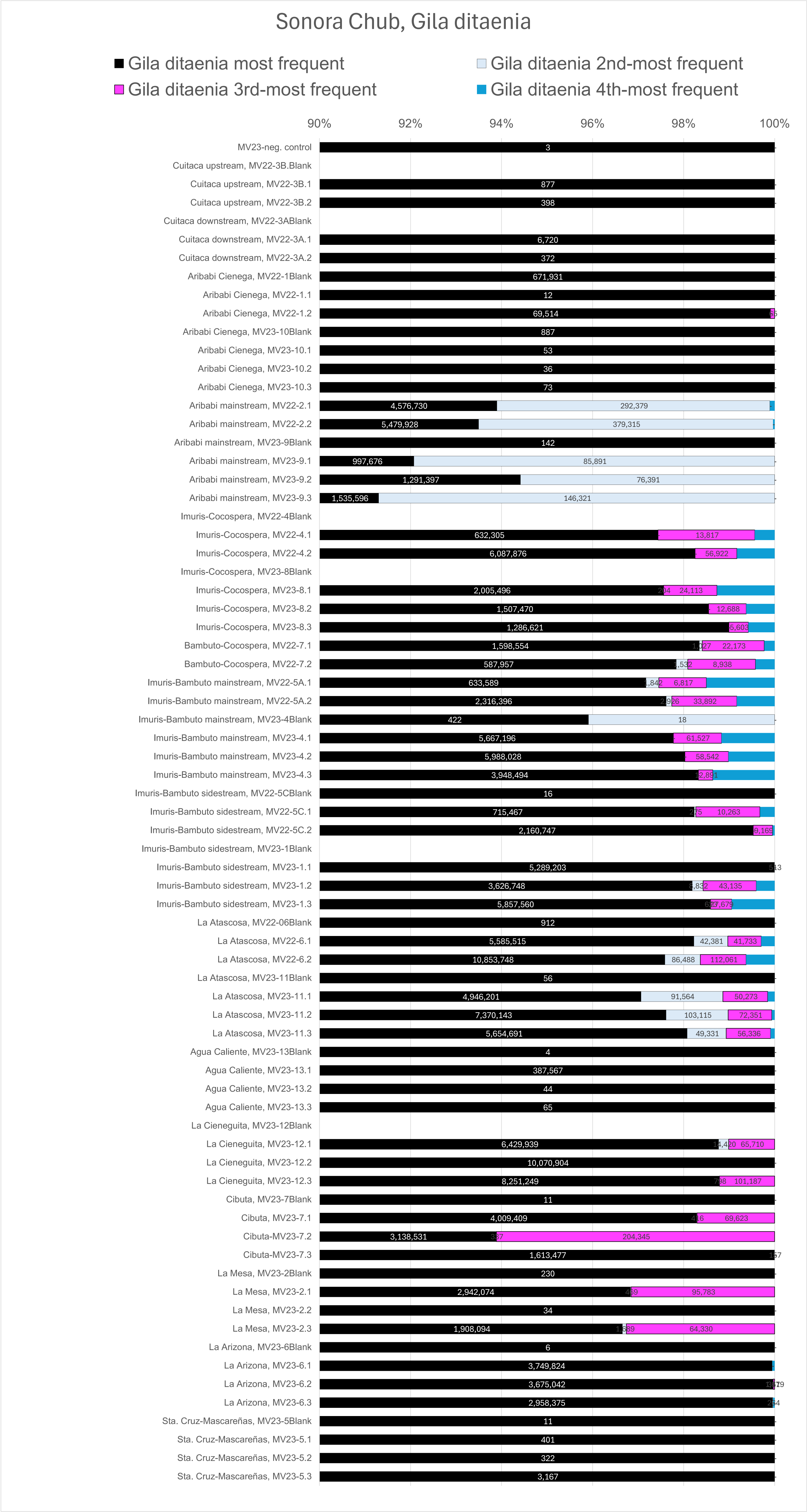

### Figure_S12_14May2026.pdf

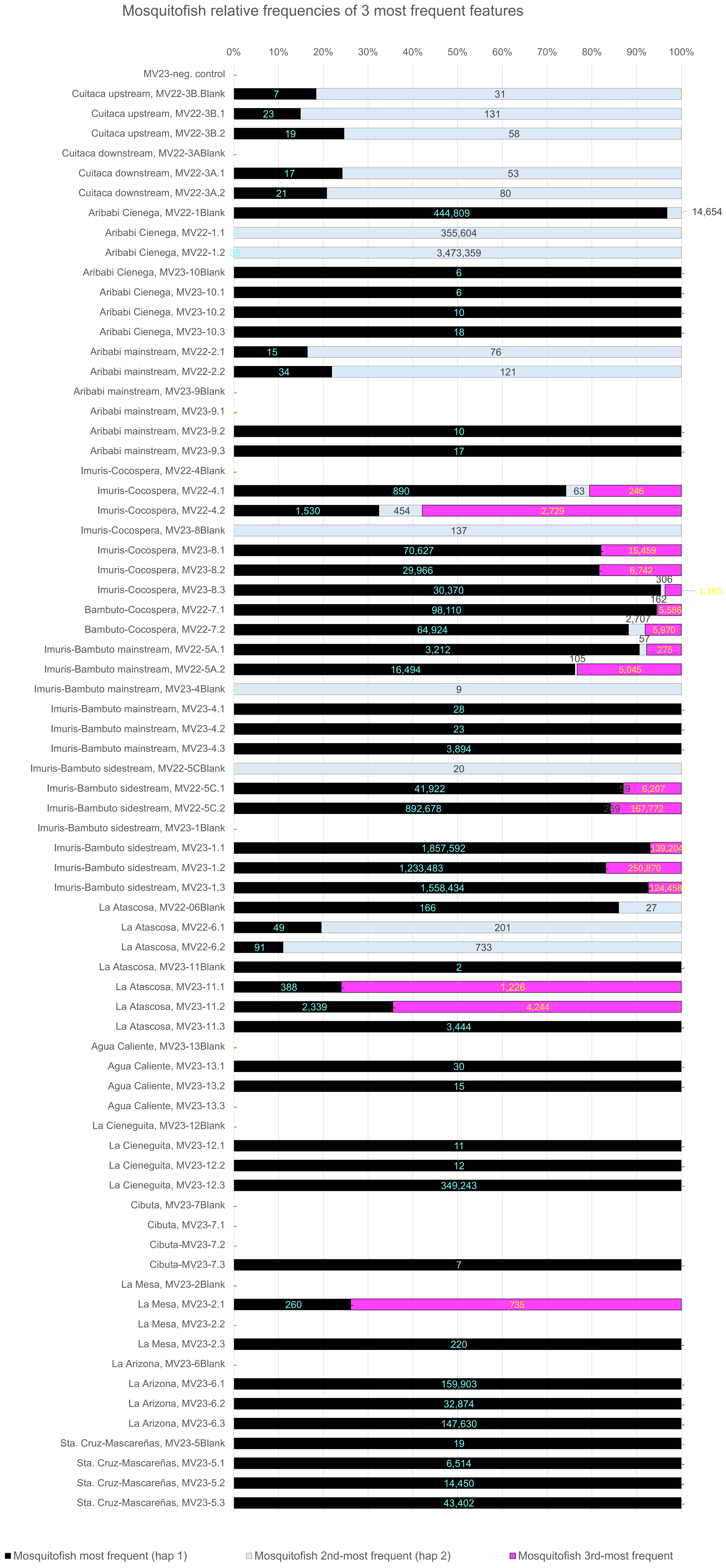
